# The *in vivo* endothelial cell translatome is highly heterogeneous across vascular beds

**DOI:** 10.1101/708701

**Authors:** Audrey C.A. Cleuren, Martijn A. van der Ent, Hui Jiang, Kristina L. Hunker, Andrew Yee, David R. Siemieniak, Grietje Molema, William C. Aird, Santhi K. Ganesh, David Ginsburg

## Abstract

Endothelial cells (ECs) are highly specialized across vascular beds. However, given their interspersed anatomic distribution, comprehensive characterization of the molecular basis for this heterogeneity *in vivo* has been limited. By applying endothelial-specific translating ribosome affinity purification (EC-TRAP) combined with high-throughput RNA sequencing analysis, we identified pan EC-enriched genes and tissue-specific EC transcripts, which include both established markers and genes previously unappreciated for their presence in ECs. In addition, EC-TRAP limits changes in gene expression following EC isolation and *in vitro* expansion, as well as rapid vascular bed-specific shifts in EC gene expression profiles as a result of the enzymatic tissue dissociation required to generate single cell suspensions for fluorescence-activated cell sorting (FACS) or single cell RNA sequencing analysis. Comparison of our EC-TRAP to published single cell RNA sequencing data further demonstrates considerably greater sensitivity of EC-TRAP for the detection of low abundant transcripts. Application of EC-TRAP to examine the *in vivo* host response to lipopolysaccharide (LPS) revealed the induction of gene expression programs associated with a native defense response, with marked differences across vascular beds. Furthermore, comparative analysis of whole tissue and TRAP-selected mRNAs identified LPS-induced differences that would not have been detected by whole tissue analysis alone. Together, these data provide a resource for the analysis of EC-specific gene expression programs across heterogeneous vascular beds under both physiologic and pathologic conditions.

**Significance:** Endothelial cells (ECs), which line all vertebrate blood vessels, are highly heterogeneous across different tissues. The present study uses a genetic approach to specifically tag mRNAs within ECs of the mouse, thereby allowing recovery and sequence analysis to evaluate the EC-specific gene expression program directly from intact organs. Our findings demonstrate marked heterogeneity in EC gene expression across different vascular beds under both normal and disease conditions, with a more accurate picture than can be achieved using other methods. The data generated in these studies advance our understanding of EC function in different blood vessels and provide a valuable resource for future studies.

## Introduction

Endothelial cells (ECs) form the inner lining of all vertebrate blood vessels, providing a critical barrier between circulating blood and parenchymal cells. In addition to maintaining blood fluidity and regulating vascular tone, ECs control the transport of nutrients and metabolites to and from underlying tissues while also contributing to host defense and the control of inflammatory processes. Consistent with their diverse microenvironments and specialized functions, ECs display remarkable morphologic and structural heterogeneity across organs (1–5). Initial studies of well-established EC markers including von Willebrand factor (*Vwf*), platelet endothelial cell adhesion molecule (*Pecam*) and *Cd34* demonstrated heterogeneous expression of these genes across the vascular tree and in distinct vascular beds (6, 7). However, more comprehensive characterization of the EC transcriptome has been largely limited by the challenge of purifying ECs, given their highly interspersed anatomic distribution among parenchymal cells in various tissues.

Previous studies have focused on human umbilical vein endothelial cells (HUVECs) expanded in cell culture, or isolation and *in vitro* expansion of ECs from distinct vascular beds such as the brain or liver (8–10). Although ECs expanded *in vitro* may retain some tissue-specific characteristics, the loss of microenvironmental cues could result in shifts in the expression program that no longer faithfully reflect the *in vivo* EC transcriptome (11–14). Several *in vivo* phage display approaches have been used to identify vascular bed-specific markers or ‘vascular zip codes’, though generally resulting in the identification of only a limited number of EC binding peptides (15–18). In addition, fluorescence-activated cell sorting (FACS) of cells endogenously marked with a fluorescent protein (19–23) or via intravital staining (24), and more recently single cell RNA sequencing (RNASeq) analyses (25–30), have been performed to determine cell type-specific transcriptomes. However, these latter methods rely on the preparation of single cell suspensions following enzymatic tissue dissociation often followed by flow sorting under high shear stress, which may affect gene expression programs.

Mouse transgenic approaches have enabled transcriptional profiling directly *in vivo* via translating ribosome affinity purification (TRAP) (31–36). This method relies on the inducible expression of an epitope-tagged ribosomal protein in a cell type-specific pattern, thus facilitating isolation of ribosome-associated transcripts, also known as the translatome (37), for particular cellular subsets directly from a complex mixture of cells. Here we report the application of EC-specific TRAP (EC-TRAP) combined with high-throughput RNASeq analysis to characterize EC translatomes across multiple vascular beds. Our findings demonstrate a high degree of tissue-specific endothelial heterogeneity, as well as notable shifts in EC mRNA content during the process of enzymatic tissue dissociation. Using EC-TRAP we also demonstrate vascular bed-specific variation in endothelial reactivity in response to lipopolysaccharide (LPS) exposure, which would not have been identified by whole tissue RNASeq data analysis.

## Results and Discussion

### In vitro expansion of ECs leads to phenotypic drift

Microarray analysis was performed on ECs isolated from heart or kidney, either directly after isolation (day 0) or after additional expansion in culture for 3 days. As shown by principal component analysis (Figure 1), *in vitro* expansion of primary ECs results in major shifts in gene expression profiles, with the expression profiles of heart and kidney more closely resembling each other after 3 days in culture, than compared to freshly isolated ECs at day 0. These data suggest that removal of the native environment results in phenotypic drift with regression toward a common EC transcription profile. This finding supports previous reports (11–14) demonstrating an important role for microenvironmental cues in maintaining the molecular heterogeneity of ECs.

**Figure 1:**
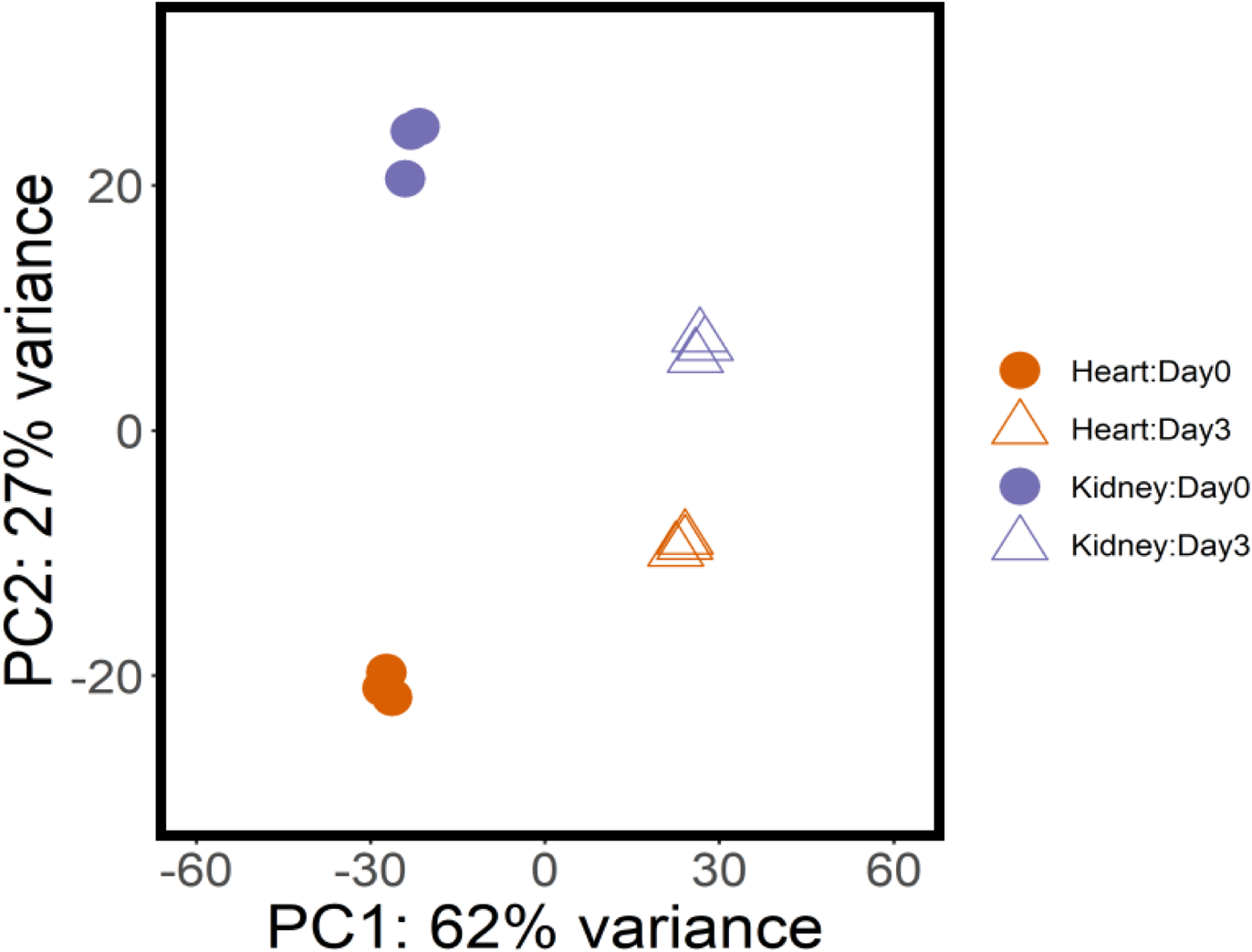
*In vitro* expansion of primary ECs leads to changes in expression profiles. Principal component analysis of expression profiles obtained from heart and kidney EC samples immediately after isolation (day 0) identified 4783 differentially expressed genes (FDR<10%) between tissue origins. *In vitro* expansion for 3 days resulted in significant shifts in expression programs, with the number of differentially expressed genes being reduced to 2397 between heart versus kidney ECs, indicating phenotypic drift occurring in the absence of a native microenvironment; n=3 biologic replicates per condition.

### Rpl22 isoform analysis to assess tissue-specific EC content

To probe EC heterogeneity directly *in vivo*, we applied TRAP by taking advantage of the RiboTag mouse, which carries a conditional *Rpl22* allele (35). We, and others, have previously shown effective EC targeting using *Cre*-recombinase driven by the *Tek* promoter in evaluating the cellular origin of coagulation factor VIII (38, 39). As shown in Figure 2A-D, analysis of mice carrying a *Tek-Cre* transgene (*Tek-Cre^+/0^*) (40, 41) together with the *Rosa^mTmG^* reporter (42) confirms efficient targeting of ECs *in vivo*. The high degree of EC sensitivity and specificity observed for the *Tek-Cre* transgene suggests that the extent of genomic *Rpl22* excision in organs from *Rpl22^fl/fl^, Tek-Cre^+/0^* mice should accurately reflect the relative fraction of ECs within those tissues. Quantitative analysis of *Rpl22* targeting by direct high-throughput genomic sequence analysis demonstrated the lowest fraction of ECs in brain (5.5±0.9%), followed by liver (13.5±1.7%), kidney (16.3±2.6%), and heart (38.6±3.8), with the highest fraction observed in the lung (48.4±6.7%) (Figure 2E). Although we cannot exclude a minor contribution from non-ECs (transiently) expressing *Tek* by this approach (see below), the quantitative *Rpl22* data are consistent with our microscopic images from *Rosa^mT/mG^, Tek-Cre^+/0^* mice (Figure 2A-D) and comparable to previous estimates of EC content based on histologic examination and FACS analysis (20, 43).

**Figure 2:**
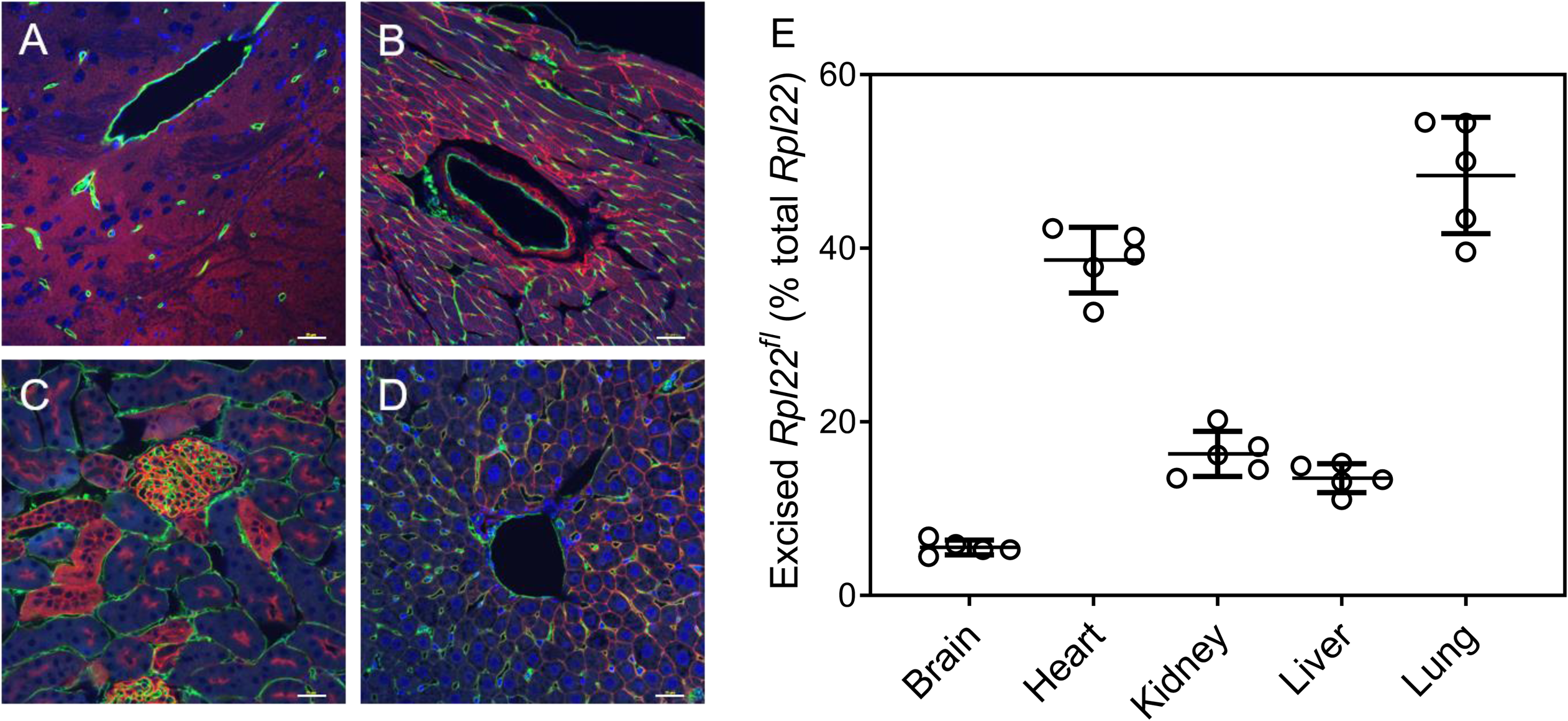
I*n vivo* targeting of *Tek*-positive cells to determine tissue-specific EC content. (A-D) Histologic analysis of *Rosa^mT/mG^,Tek-Cre^+/0^* brain (A), heart (B), kidney (C) and liver (D); *Tek*-positive cells are identified by the expression of membrane-bound green fluorescent protein, while *Tek*-negative cells express membrane-bound Tomato red fluorescent protein; scale bar = 25 µm. (E) Percentage of *Tek*-positive cells, as a proxy for EC content in each tissue, as determined by high-throughput DNA sequencing of *Rpl22* isoforms in tissues from *Rpl22^fl/fl^, Tek-Cre^+/0^* animals; n=5 biologic replicates (mean ± SD).

### Evaluation of EC-specific translatomes by TRAP

TRAP specifically captures actively translated mRNAs (translatome), which may differ quantitatively from the total cellular mRNA pool (transcriptome). To determine if tissue transcriptomes obtained prior to EC-TRAP are comparable to tissue translatomes, or if corrections are needed for potential disproportional mRNA capture due to differences in ribosome density across mRNAs, we performed TRAP and RNAseq analysis on tissues of mice expressing HA-tagged ribosomes in all cell types (*Rpl22^fl/fl^, EIIa-Cre^+/0^*). Before tissue isolation, animals were perfused with cycloheximide to block translation and stabilize tagged ribosomes on their cognate mRNAs to minimize potential shifts in ribosome distribution during tissue collection and immunoprecipitation (44). Expression profiles obtained before and after TRAP were highly reproducible between biologic replicates, with distinct patterns evident across organs. Only minor changes were observed between the total mRNA and actively translated mRNA within a given tissue (Figure 3A), resulting in a limited number of transcripts that were significantly different between the 2 mRNA pools (Supplemental Figure S1; Supplemental Table S1). These data demonstrate that signatures obtained after EC-TRAP should primarily reflect cell-specific gene expression programs, with little or no contribution from shifts in ribosome distribution, and that the transcriptome of a tissue lysate can serve as a proxy for the corresponding tissue translatome.

**Figure 3:**
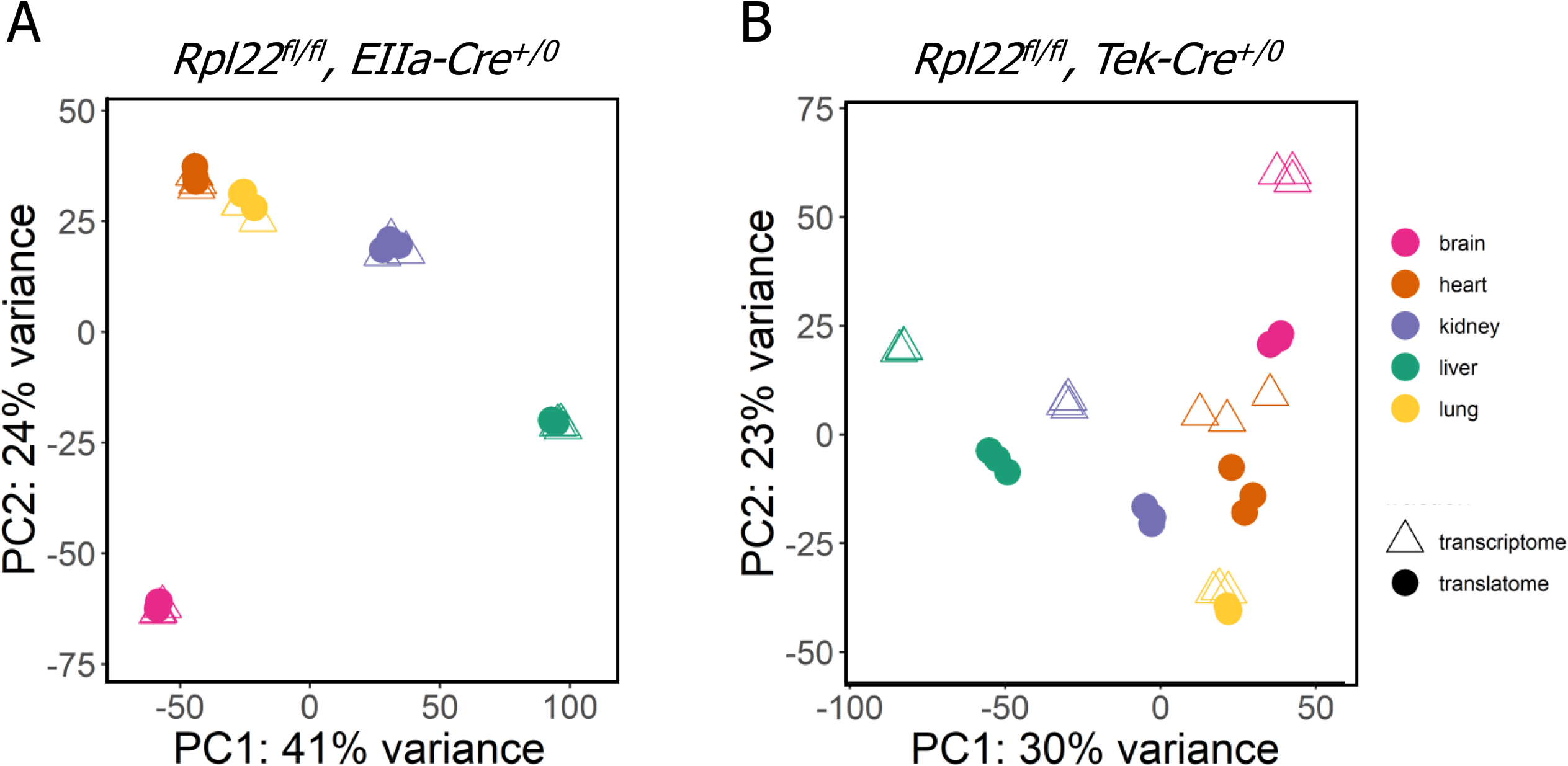
Evaluation of transcriptome versus translatome data after (EC) TRAP. (A) Mice expressing HA-tagged *Rpl22* in all cells show only minor differences between tissue-specific transcriptomes and translatomes from brain, heart, kidney, liver and lung by genome-wide principal component analysis. (B) EC translatomes obtained after TRAP are highly distinct from the unselected tissue transcriptomes, with the exception of lung. n=3 biologic replicates per tissue.

We next analyzed TRAP-selected mRNA fractions collected from multiple tissues of *Rpl22^fl/fl^, Tek-Cre^+/0^* mice after cycloheximide perfusion. To avoid contamination of TRAP-selected mRNAs with transcripts originating from parenchymal cells (35), we also evaluated the accompanying unselected tissue transcriptomes by RNAseq analysis, and calculated relative enrichment scores. These scores identify those transcripts that are more abundant in *Tek^+^* cells (enriched genes, log2 fold change >0), or more abundant in *Tek^−^* cells (depleted genes, log2 fold change <0), where the former should represent EC-specific genes and the latter parenchymal cell-specific genes (Supplemental Figure S1). As expected, GO analysis of TRAP-enriched transcripts showed highly significant associations with endothelial and hematopoietic biologic processes including “vasculature development”, “immune response” and “hemopoiesis”, while depleted gene sets showing signals consistent with parenchymal, tissue-specific processes (Supplemental Figure S2).

### Correction of TRAP-enriched genes for hematopoietic cell content

Although the *Tek-Cre* transgene provides an efficient and specific marker of the EC compartment, the *Tek* gene is also transiently expressed in hematopoietic cells (45), which could result in the false identification of mRNAs as EC-specific as a result of blood contamination of tissues. For example, despite exsanguination and cardiac perfusion prior to organ harvest to remove circulating blood cells, the genes encoding hemoglobin α and β (*Hba-a1*, *Hba-a2*, *Hbb-bs* and *Hbb-bt*) were among the most enriched transcripts across all tissues and although α-hemoglobin gene expression has been reported in arterial ECs, β-hemoglobin has not directly been associated with the endothelium (46, 47). Thus, the similar degree of enrichment observed for both α- and β-hemoglobin in our EC-TRAP suggests a contribution of HA-tagged polysomes derived from circulating blood cells that were not completely removed by perfusion prior to tissue isolation. To eliminate transcripts originating from hematopoietic cells, we applied additional computational filters based on data obtained from TRAP performed directly on peripheral blood samples from *Rpl22^fl/fl^, Tek-Cre^+/0^*, as well as EC-TRAP on tissues collected from an *Rpl22^fl/fl^, Tek-Cre^+/0^* recipient following bone marrow transplant using an *Rpl22^fl/fl^, Tek-Cre^−^* donor (see Supporting Information). In addition to removing the hemoglobin genes and other known lymphoid and myeloid cell markers, including a number of established EC markers known to be expressed by megakaryocytes/platelets (48), applying these filters provides the most specific set of EC-enriched genes. GO analysis of this more restricted EC gene set now showed enrichment limited to vascular-related biologic processes (Figure 4A) with loss of the GO terms for marrow-derived cell populations identified in the broader analysis (Supplemental Figure S2).

**Figure 4:**
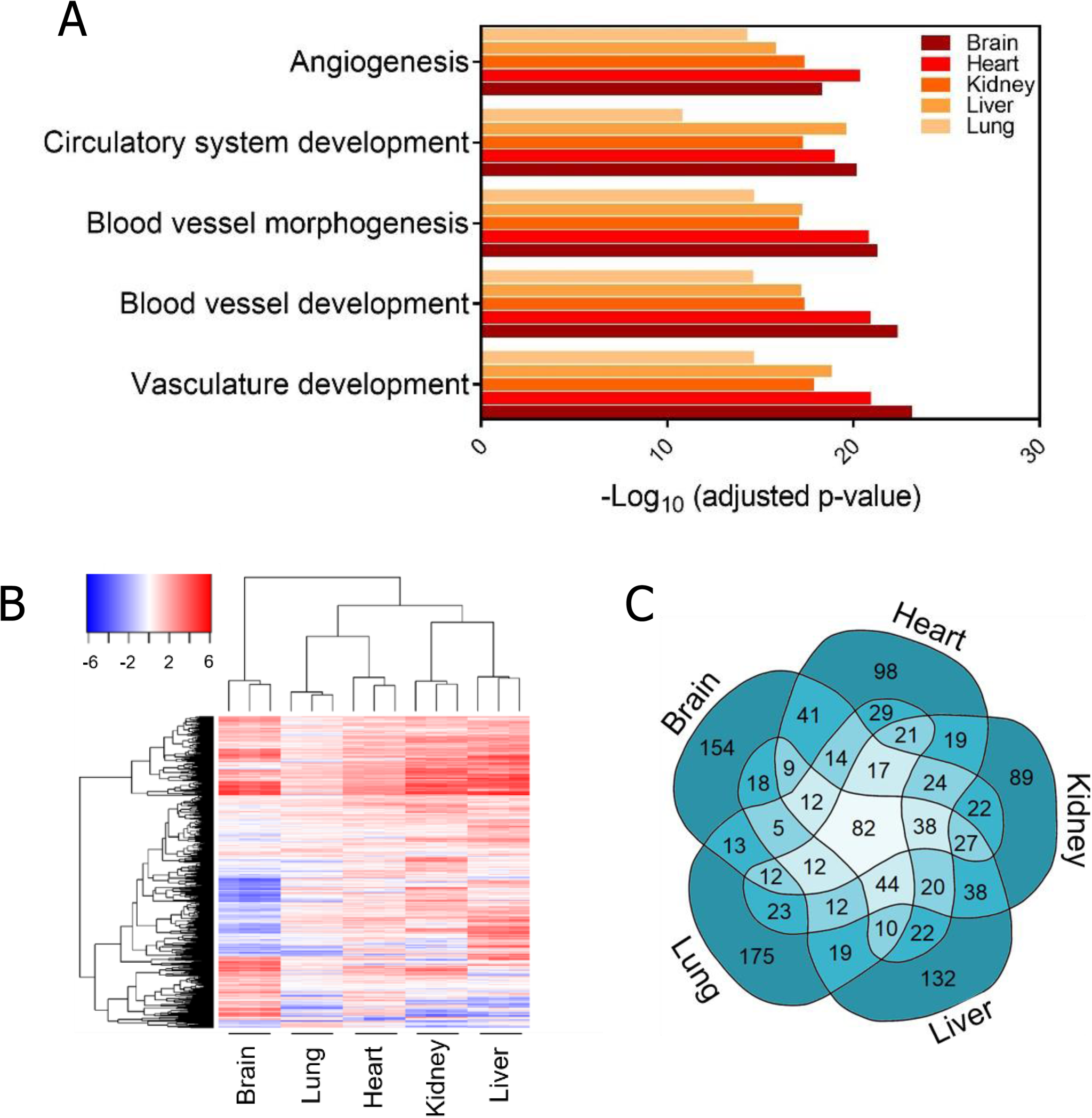
Identification of enriched transcripts after EC-TRAP. (A) Of the putative EC-enriched genes in each tissue (FDR<10%), GO analysis of the 500 top-ranked genes with the highest enrichment scores shows overrepresentation of transcripts involved in vascular-related processes. (B) Unsupervised hierarchical clustering of EC-enriched genes (log2 fold change >2 in at least 1 tissue) shows distinct, highly heterogeneous vascular bed-specific EC expression patterns. (C) Comparison of the top 500 most enriched genes per tissue identifies a group of pan-endothelial and subsets of tissue-specific EC-enriched genes. Data based on n=3 biologic replicates per tissue.

### TRAP identifies EC heterogeneity across vascular beds

Principal component analysis of EC translatomes demonstrates distinct, reproducible patterns for each organ (Figure 3B). This observation is confirmed by hierarchical clustering of enrichment scores where, consistent with the high percentage of ECs in the lung and heart, the degree of enrichment for EC transcripts in these organs is generally lower than that observed in brain, kidney and liver (Figure 4B).

Overlap in expression across the different vascular beds for the 500 top-ranked genes based on enrichment scores for each tissue is shown in Figure 4C, identifying 82 genes shared among all 5 vascular beds, including *Tek*, as expected, as well as other established pan-EC markers such as *Cdh5*, *Nos3*, *Eng* and *Robo4*. This list also includes *Ephb4*, *Dll4* and *Flt4*, specifically marking arterial, venous and lymphatic ECs, respectively, demonstrating efficient targeting of all 3 EC subtypes (49, 50) (Table 1). In addition to the pan-EC markers that have previously been associated with EC function and/or localization (23, 32, 51–53), our EC-TRAP analysis also identified potential novel pan-EC genes, including *Gm20748* (also known as *Bvht*) and *Eva1b*, with *in situ* hybridization confirming colocalization of these transcripts with *Tek* transcripts in ECs of brain, kidney and liver (Figure 5A-B).

**Figure 5:**
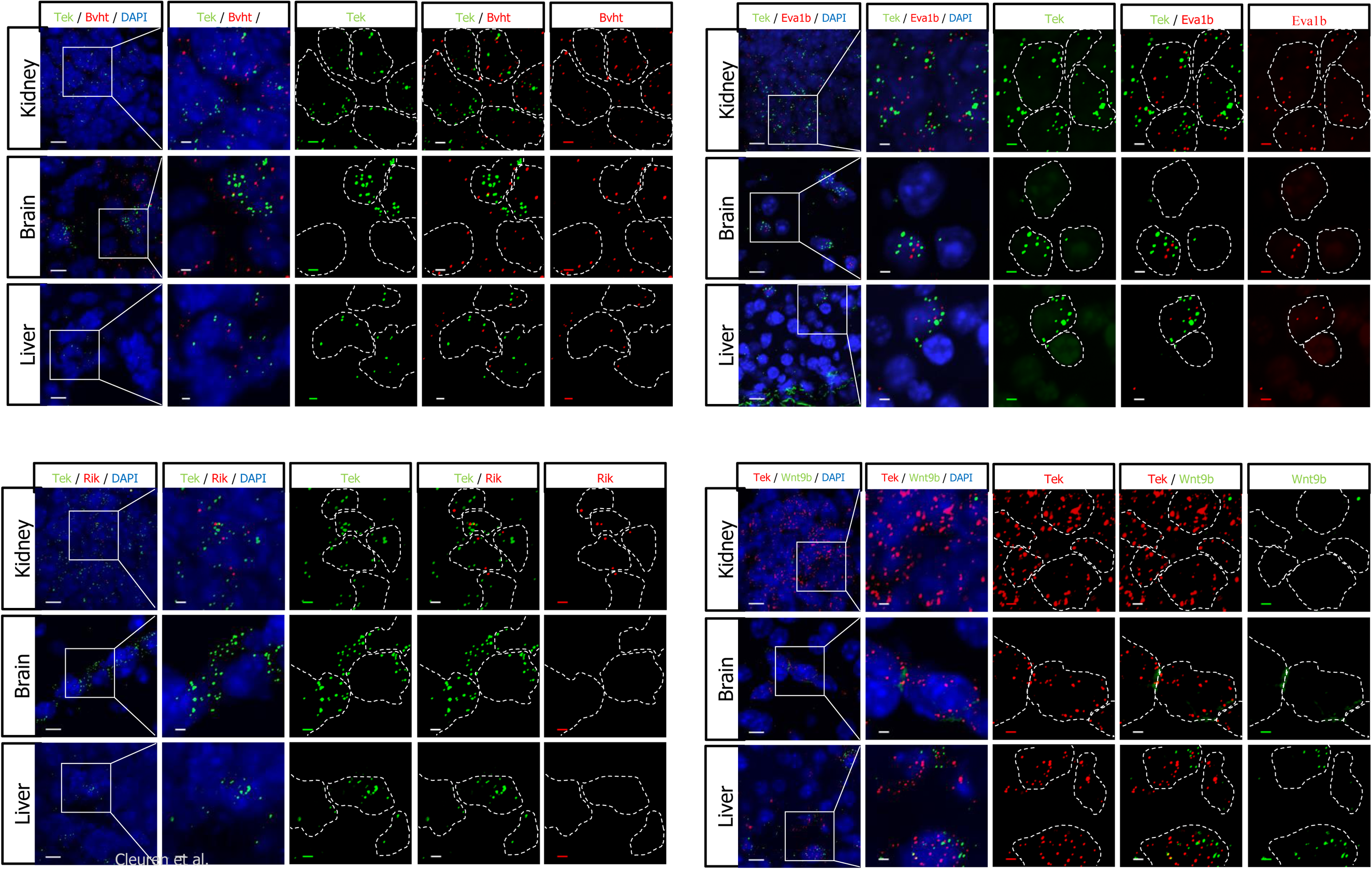
Single-molecule *in situ* hybridization validates EC-enriched transcripts identified by TRAP. (A-B) Colocalization of *Bvht* and *Eva1b*, both identified as pan-EC markers by EC-TRAP, with *Tek* by *in situ* hybridization in kidney, brain and liver. (C-D) *3110099E03Rik* and *Wnt9b* were identified as kidney- and liver-specific EC markers, respectively and were only present in the corresponding tissues, where they also colocalized with *Tek*, confirming their EC origin. Dotted lines in the insets outline the cell nuclei as indicated by DAPI. Scale bars are 10 µm in overviews (left column in each panel), and 2.5 µm in insets (remaining columns).

**Table 1:**
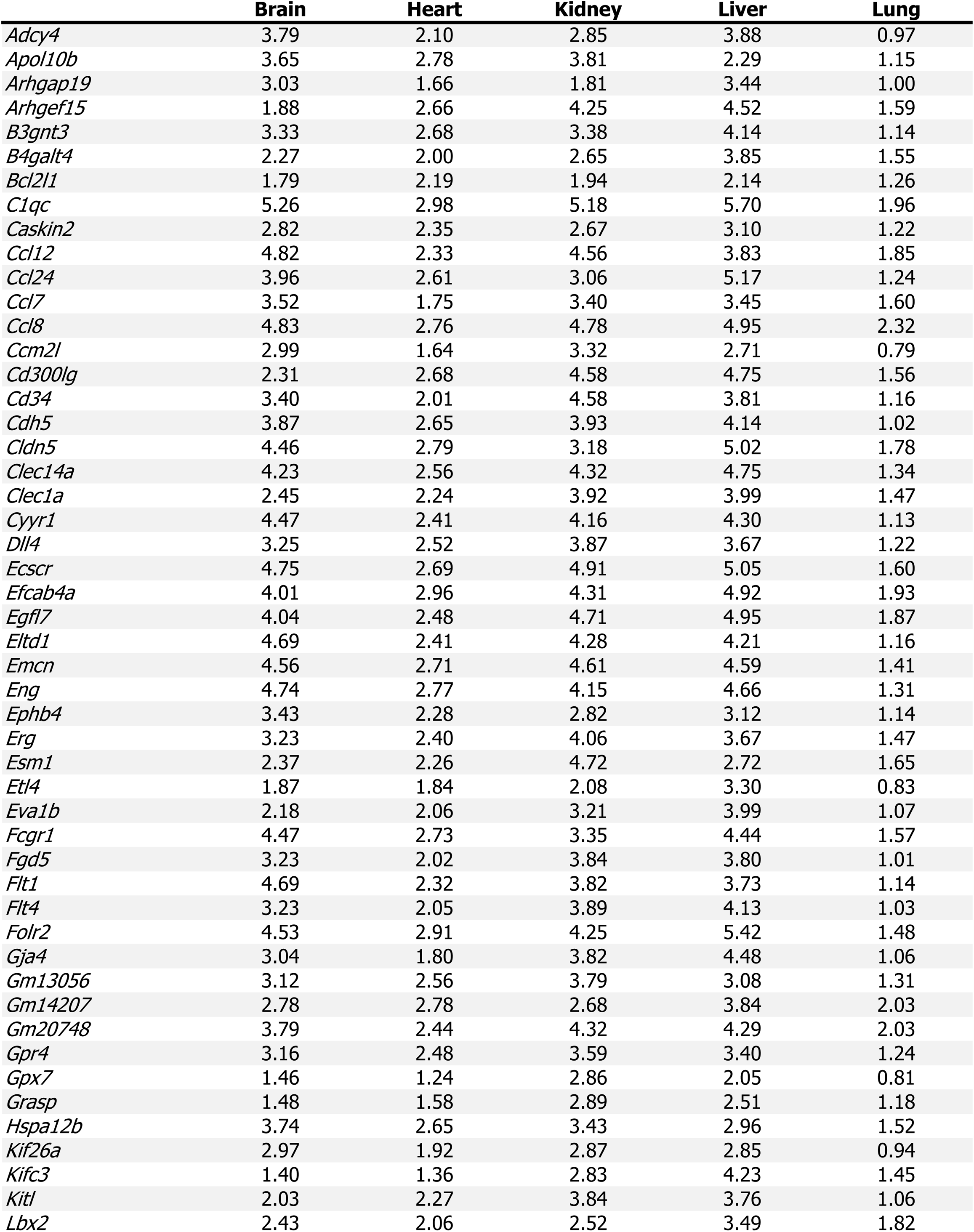
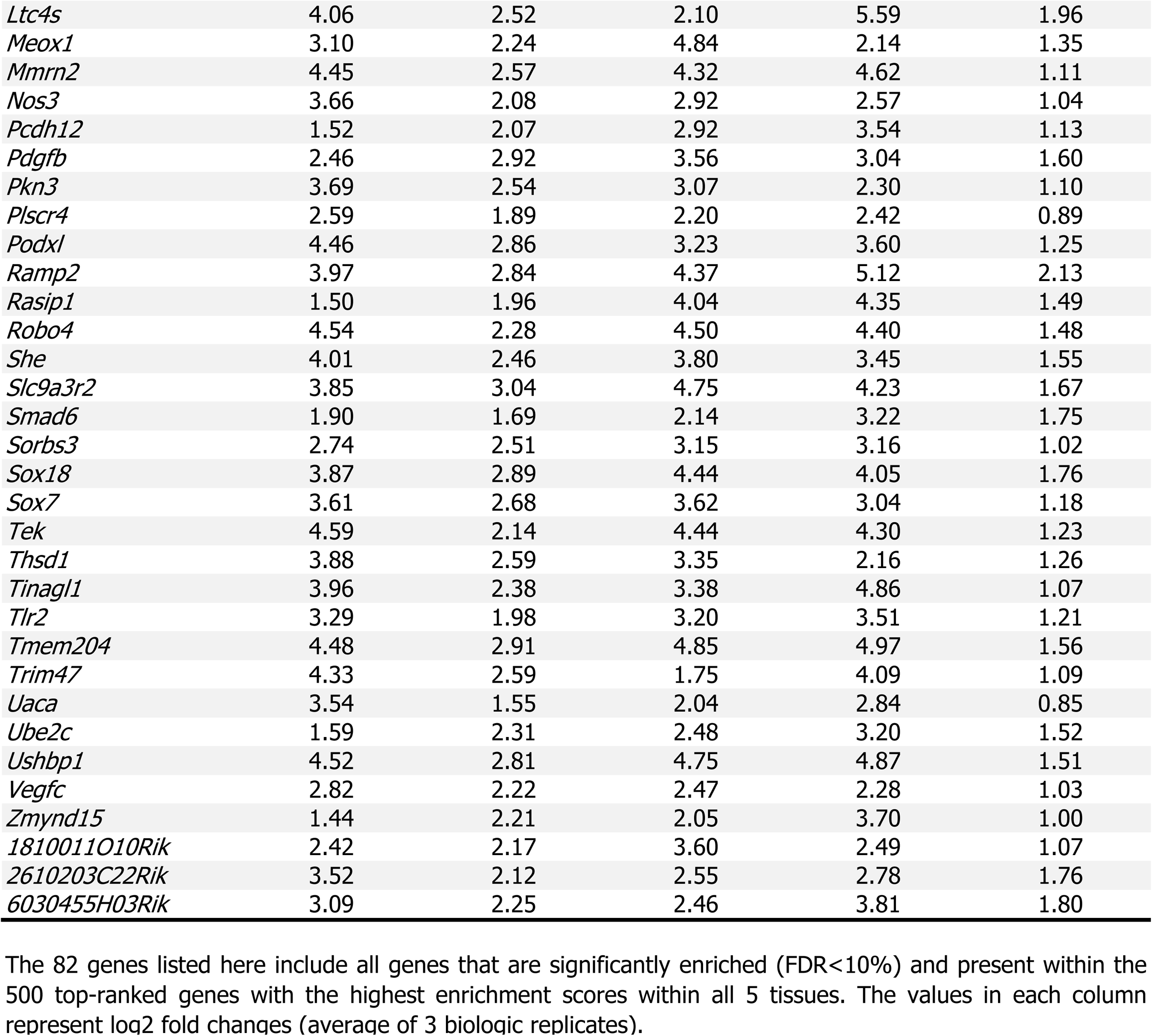
82 pan-endothelial genes present in all 5 tissues after EC-TRAP.

The EC-TRAP data identify marked heterogeneity of gene expression levels for EC-enriched genes across tissues, including genes restricted to a single or several vascular beds. An interesting subset of genes appears to be EC-enriched in one or more tissue(s), with expression levels more abundant in non-ECs in another tissue (Supplemental Figure S3; Supplemental Table S2). For example, whereas several solute carriers play important roles in proximal tubules in the kidney, and are thus depleted after kidney EC-TRAP, a number of these genes are significantly enriched in brain ECs (Supplemental Table S2). Focusing on EC-specific genes with expression restricted to a single vascular bed identified the largest number of unique markers for brain ECs, including several genes previously associated with the blood brain barrier (e.g. *Rad54b*, *Zic3* and *Slco1c*) (20, 25, 27, 54). Smaller subsets of tissue-specific EC markers were also identified for heart, kidney, liver and lung (Table 2), with specific kidney EC expression of lincRNA *3110099E03Rik* and liver EC expression of *Wnt9b* confirmed by *in situ* hybridization (Figure 5C-D).

**Table 2:**
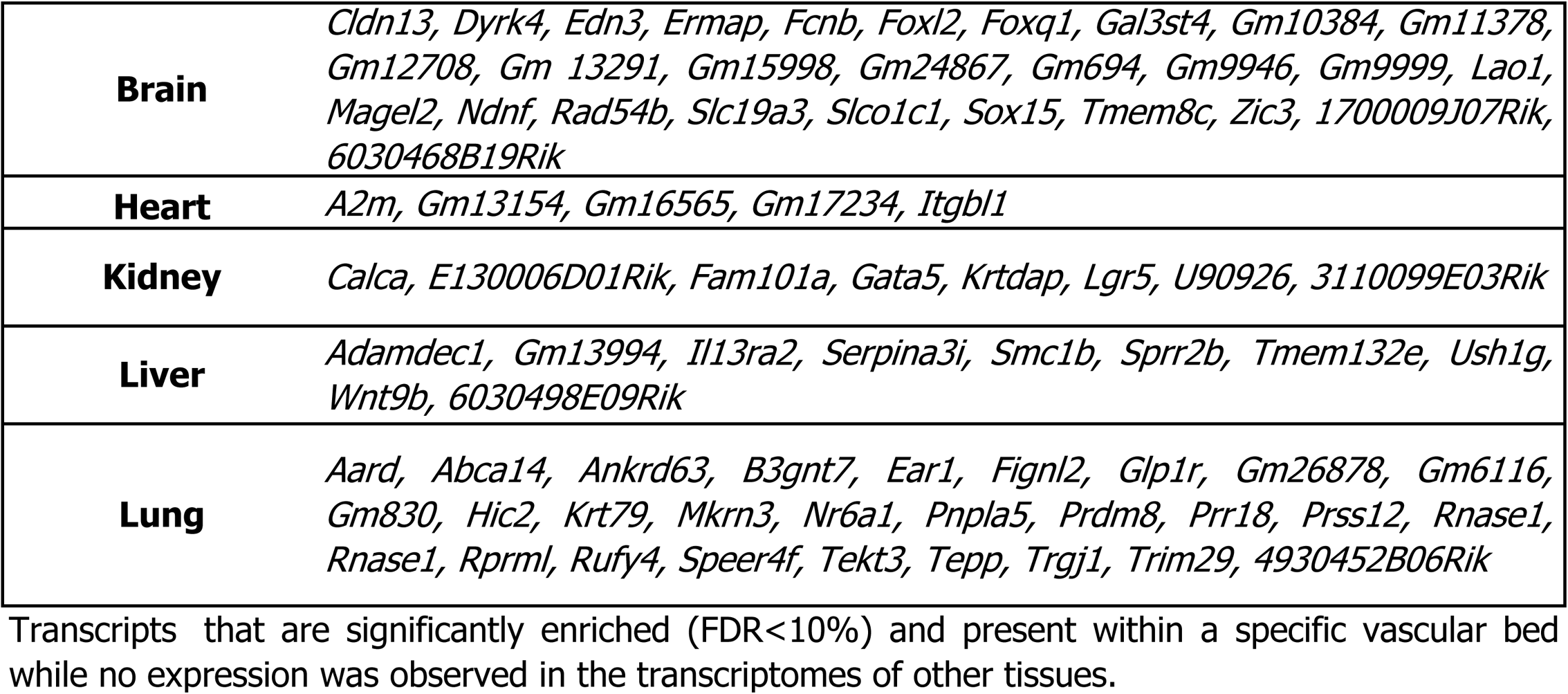
Tissue-specific, EC restricted transcripts identified by EC-TRAP.

### Comparison between EC-TRAP and single cell RNASeq

Recently developed single cell RNASeq methods make it possible to distinguish cellular subtypes within complex mixtures of cells, including multiple subsets of ECs, as previously reported for the brain (23, 27). However, several studies raise the concern that the isolation procedures required to obtain individual cells for RNASeq analysis may perturb relative mRNA abundance (55–57). To evaluate the potential impact of such changes in EC expression programs across tissues, perfusion and enzymatic tissue dissociation were performed in the absence of cycloheximide, to mimic the effects of single cell isolation procedures (including continuing changes in mRNA transcription and translation), until the addition of cycloheximide-containing buffer prior to TRAP, approximately 90 minutes after organ harvest.

Although some degree of vascular bed-specific gene expression patterns persist, substantial changes are observed when compared to the profiles obtained from cycloheximide-perfused animals (Figure 6A). Whereas the kidney endothelium showed relatively small shifts in expression profiles between isolation methods, liver and brain ECs exhibit larger deviations as well as lower correlations between biologic replicates. Indeed, 48.2% of genes expressed in brain ECs (3794/7866) and 21.6% expressed in liver ECs (2306/10696) are differentially expressed when comparing the 2 preparation methods, with only 7.4% for kidney EC genes (779/10475). As a result, hierarchical clustering of EC translatomes identify liver and brain ECs as more similar based on sample preparation method than by tissue of origin (Figure 6B).

**Figure 6:**
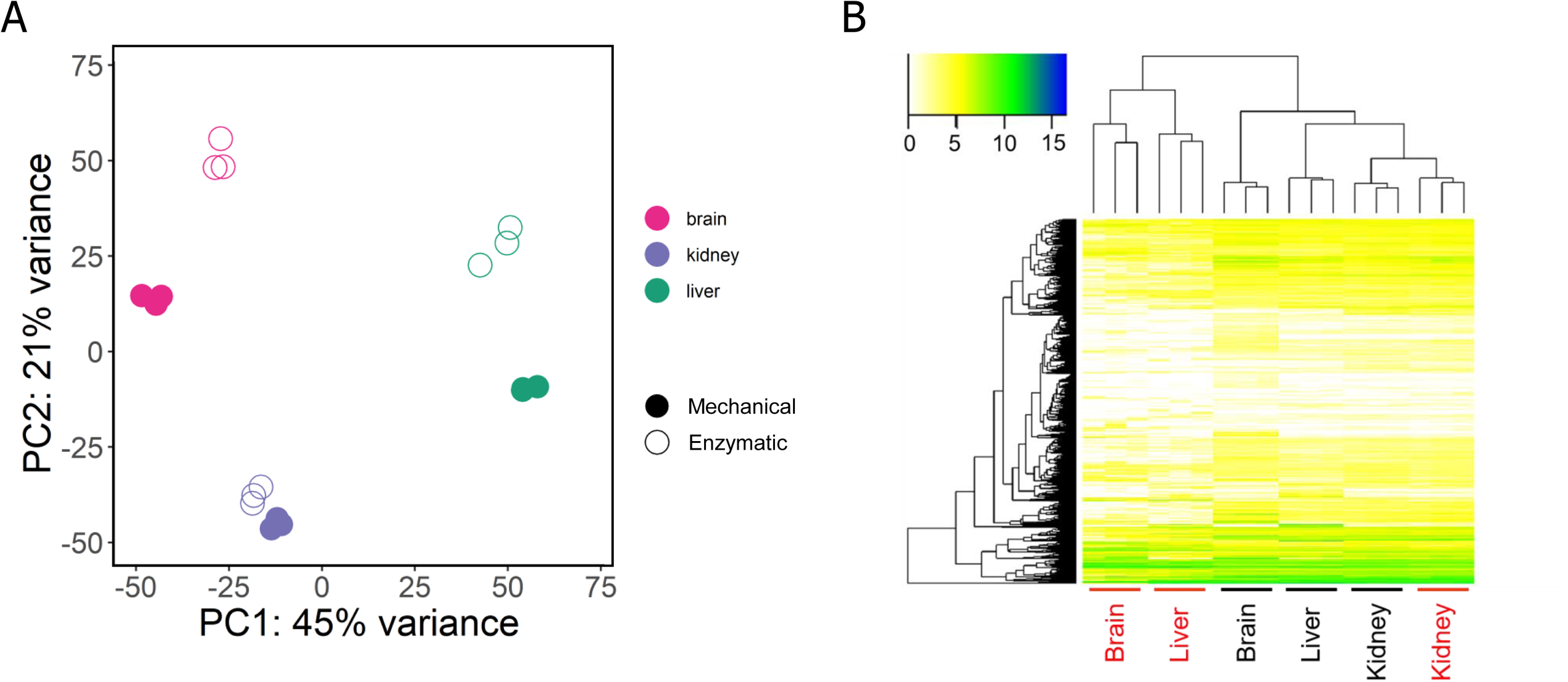
Dissociation-induced changes in EC gene expression programs. (A) PCA analysis of EC translatomes obtained from TRAP samples after cycloheximide perfusion and mechanical dissociation versus samples following enzymatic tissue dissociation without cycloheximide perfusion in brain, liver and kidney. (B) mRNA abundance presented as log2(TPM) shows clustering of brain and liver EC translatomes by preparation method (black: mechanical, red: enzymatic) rather than by tissue type. Data from n=3 biologic replicates per tissue.

In addition to the potential for shifts in the EC expression program during the preparation of single cells, current single cell RNASeq methods exhibit a relatively limited sensitivity to detect low abundance transcripts (58, 59). Consistent with these previous reports, of the 100 most abundant EC-enriched transcripts per tissue identified in our EC-TRAP data set, many have also been identified in ECs based on single cell RNASeq data (median 50, range 28-77; Supplemental Table S3). In contrast, only a few of the 100 lowest abundant genes were detected by single cell RNAseq (median 2, range 0-24) (25–27, 29).

Taken together, the EC-TRAP approach combined with *in vivo* cycloheximide perfusion prior to mRNA preparation reported here, should result in a more accurate snapshot of native EC profiles across different vascular beds, including low abundant transcripts, than is possible with current single cell isolation and RNASeq methods.

### Endotoxemia induces tissue-specific changes in the EC gene expression program

The endothelium provides a key line of host defense in response to microbial pathogens. To examine the tissue-specific response in an animal model for bacterial endotoxemia, *Rpl22^fl/fl^, Tek-Cre^+/0^* mice were injected with lipopolysaccharide (LPS). Analysis of whole organ transcriptomes (including both ECs and parenchymal cells) demonstrated marked changes in gene expression in response to LPS, with gene ontology analysis yielding highly significant values for terms associated with host defense and immune response (Supplemental Figure S4A). The extent of gene changes was highly variable across tissues, with a markedly reduced number of LPS responsive mRNAs evident for intact brain tissue (Figure 7A). Analysis of the EC-specific response, however, showed similar numbers of genes affected by LPS treatment across all tissues, including brain (Figure 7B). These data suggest a dramatic reduction in the effect of LPS on brain parenchymal cells compared to other tissues, consistent with a high degree of protection provided by the blood-brain barrier (60). In addition, this also illustrates a limitation of RNASeq of whole tissues, where cell type-specific responses can be masked by changes in more abundant cellular subsets. Interestingly, whereas most vascular bed-specific EC translatomes exhibited considerably more downregulated than upregulated transcripts in response to LPS, the opposite pattern was observed for brain ECs (Figure 7B, Supplemental Figure S4B-C).

**Figure 7:**
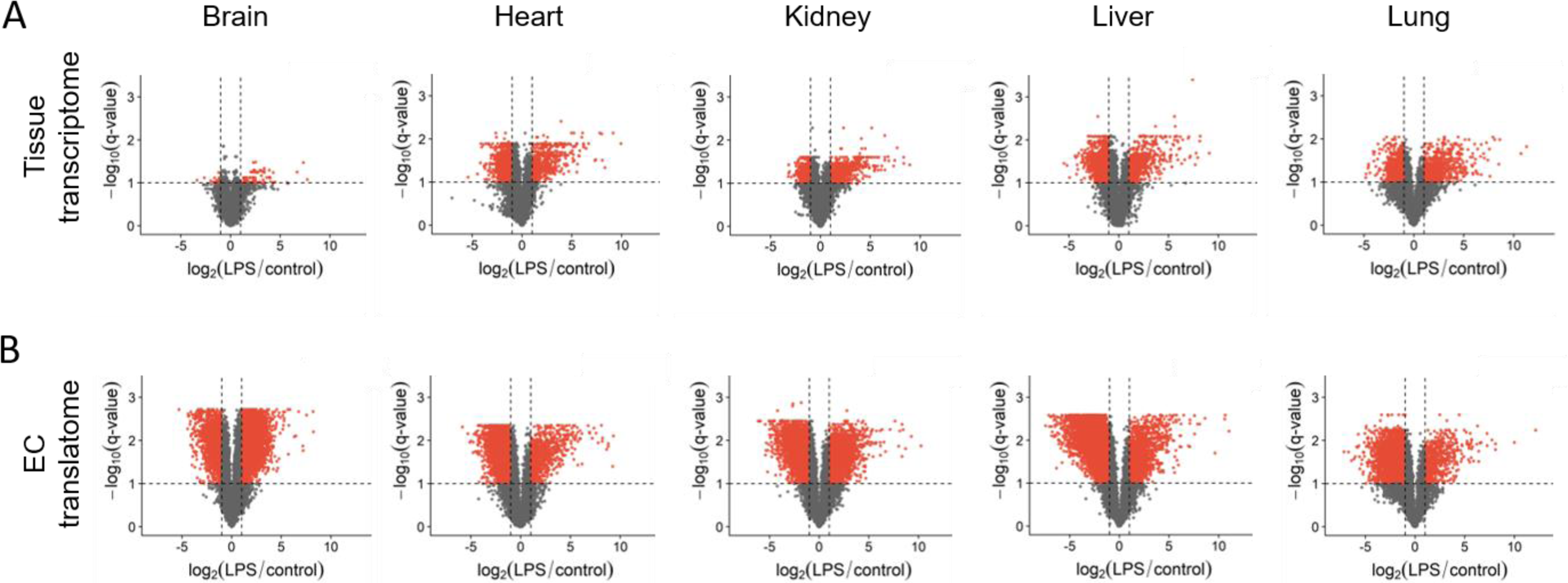
*In vivo* LPS-induced changes uncovered by EC-TRAP. (A-B) Volcano plots indicate significantly affected genes by LPS treatment (red), either for tissue transcriptomes (A) or in the EC translatome identified by EC-TRAP (B). The horizontal dotted lines mark an FDR cut-off <10%, with the vertical dotted lines denoting log2 fold changes of >1 or <-1. Data averaged from n=3 biologic replicates per tissue.

Consistent with previous reports (61–63), EC-enriched genes upregulated by LPS include several adhesion molecules required for leukocyte recruitment and migration, while genes involved in maintenance of the EC barrier were downregulated. Furthermore, a shift toward a procoagulant state was observed, with reduced expression of anticoagulant and profibrinolytic genes (Supplemental Table S4). Within this limited set of genes, expression of the protein C receptor (*Procr*) provides an example of vascular bed-specific variation in EC reactivity in response to LPS, with significant downregulation of *Procr* in brain, kidney and heart, while expression in liver and lung was increased. These data could explain the organ-specific susceptibility to increased vascular leakage after LPS challenge previously reported for mice with reduced *Procr* expression (64).

## Conclusion

The interspersed anatomic distribution of ECs and marked dependence of their gene expression programs on microenvironmental cues have constituted major challenges to the study of EC function and responses to (patho)physiological stimuli *in vivo*. The EC-TRAP strategy described here provides a powerful tool for the analysis of the EC translatome across multiple vascular beds and in response to pathologic challenges in the whole animal. Our findings demonstrate that this method offers an accurate *in vivo* snapshot of tissue-specific EC expression profiles, resulting in better cellular resolution than whole tissue RNASeq, while maintaining a high degree of sensitivity to detect low abundant transcripts. In addition, these data provide a database of tissue-specific EC gene expression that should be a useful reference for other studies, including the identification of tissue-specific targets for drug delivery. Future refinement of EC-TRAP taking advantage of intersectional genetic approaches based on combined expression of 2 or more EC markers (65) may lead to greater resolution of EC subsets within a tissue, particularly in combination with single cell RNAseq methods. Finally, future applications of this approach should significantly advance our understanding of EC function *in vivo* under both physiologic and pathophysiologic conditions, potentially leading to the identification of new disease-specific biomarkers as well as potential novel therapeutic targets.

## Material and Methods

All animal experiments were approved by the Institutional Animal Care and Use Committee of the University of Michigan. Detailed descriptions of the mice and procedures used in these studies, including sample preparations for TRAP and high-throughput RNA sequencing, data analysis and follow-up validation by single-molecule fluorescent *in situ* hybridization assays, are provided in the Supporting Information Materials and Methods.

## Data availability

All RNASeq data supporting these studies will be made available through the Gene Expression Omnibus.

## Acknowledgements

We would like to thank Lauren Janes, Kate Spokes and Peter Zwiers for their expert technical assistance with the *in vitro* experiments, and Eliane Popa for the helpful discussions. This research was supported by grants from the National Institutes of Health (R35HL135793 to D.G. and R01HL122684 to S.G.), and the American Heart Association (16POST27770093 to A.C.). D.G. is an investigator of the Howard Hughes Medical Institute.

## Supplementary Material and Methods

### Mice

All mice were housed under standard conditions according to the University of Michigan’s Unit of Laboratory Animal Medicine guidelines. Mice were originally obtained from The Jackson Laboratory and bred to C57BL/6 for at least 5 generations, either by the donating institute or The Jackson Laboratory. Upon arrival, colonies were subsequently maintained on a C57BL/6J background, with experimental animals being backcrossed for at least 3 additional generations. Genotyping was performed on DNA isolated from tail clips using the PCR primers listed in Supplemental Table S5.

RiboTag mice (1) (*Rpl22^fl/fl^*; stock 011029: B6N.129-*Rpl22^tm1.1Psam^*/J) carrying a conditional hemagglutinin (HA)-epitope tagged ribosomal protein *Rpl22* gene were crossed to *Tek-Cre* recombinase transgenes (2, 3) (stock 004128: B6.Cg-Tg(Tek-cre)12Flv/J) to generate animals with HA-tagged polysomes in endothelial and hematopoietic cells, while other cell types are expected to express only the wildtype RPL22. However, *Tek-Cre* has previously also been described to target *loxP*-flanked sequences in germ cells (2). As this occurs at a lower frequency in males than in females, only *Tek-Cre^+/0^* males were used for matings, and the efficiency and specificity of the *Tek-Cre* recombinase was evaluated by breeding *Tek-Cre^+/0^* to *Rosa^mT/mG^* reporter mice (4) (stock 007676: B6.129(Cg)-*Gt(ROSA)26Sor^tm4(ACTB-tdTomato,-EGFP)Luo^*/J).

*Rpl22^fl/fl^* mice were also bred to *Pf4-Cre* recombinase males (5) (stock 008535:C57BL/6-Tg(Pf4-icre)Q3Rsko/J) or *EIIa-Cre* transgenes (6) (stock 003724: B6.FVB-Tg(EIIa-cre)C5379Lmgd/J) to determine megakaryocyte/platelet specific transcripts, or to generate animals with ubiquitously expressed HA-tagged RPL22, respectively.

### Blood and tissue collection

Ten-week-old *Rpl22^fl/fl^, Tek-Cre^+/0^* males were anesthetized with 50 µg pentobarbital/g body weight and blood was drawn via cardiac puncture into a 9:1 sodium citrate solution (Sigma-Aldrich) containing cycloheximide (Sigma-Aldrich) to a final concentration of 100 µg/mL. Following the blood draw, intracardiac perfusion with 12 mL 100 µg/mL cycloheximide in phosphate buffered saline (PBS) was performed to lock the ribosome onto the mRNA (7) and limit blood contamination of tissue samples. After perfusion, organs were harvested, snap frozen in liquid nitrogen and stored at −80°C until further processing. Blood samples were immediately processed and treated with ammonium-chloride-potassium (ACK) lysis buffer (Thermo Fisher Scientific) according to the manufacturer’s protocol to remove red blood cells. The resulting cell pellet was resuspended in polysome buffer (50 mM Tris pH 7.5, 100 mM KCl, 12 mM MgCl2, 1% Igepal CA-630 (Sigma-Aldrich), 1 mM dithiothreitol, 200 U/mL RNaseOUT (Invitrogen), 1 mg/mL heparin sodium salt (Sigma-Aldrich), 100 µg/mL cycloheximide and an EDTA-free protease inhibitor cocktail (Roche)) followed by immunoprecipitation.

To study the platelet transcriptome, blood samples from five 10 week old *Rpl22^fl/fl^, Pf4-Cre^+/0^* males were pooled and centrifuged at 100 x g for 5 minutes to obtain platelet rich plasma and platelets were subsequently pelleted by centrifugation at 800 x g for 10 minutes and resuspended in polysome buffer.

### Lipopolysaccharide treatment

For the LPS-induced endotoxemia model, *Rpl22^fl/fl^,Tek-Cre^+/0^* mice were injected intraperitoneally with LPS (*E. Coli* 026:B6 from Sigma-Aldrich) at 1 µg/g body weight 4 hours before blood draw, cycloheximide perfusion and tissue collection.

### Enzymatic tissue dissociation

To study the effects of enzymatic tissue dissociation on the EC expression program, tissues were collected after exsanguination by cardiac puncture and intracardiac perfusion with 12 mL PBS. Brain, kidney and liver samples were harvested and immediately minced and incubated with Collagenase A (25 mg/mL; Roche), Dispase II (25 mg/mL; Roche) and DNase I (250 µg/mL; Roche) at 37°C for 30 minutes (8). The subsequent cell suspension was filtered through a 70 µm cell strainer and cells were washed twice with ice-cold PBS before adding polysome buffer and being directly subjected to TRAP.

### Bone marrow transplantation

Bone marrow was harvested from 8-week old *Rpl22^fl/fl^,Tek-Cre^+/0^* or *Rpl22^fl/fl^,Tek-Cre^−^* donors and 5-10 million donor cells were injected into recipient *Rpl22^fl/fl^,Tek-Cre^+/0^* mice via the tail vein as previously described (9). Eight weeks after transplantation, mice were anesthetized and perfused with cycloheximide/PBS after which tissues were harvested, snap frozen in liquid nitrogen and stored at −80° until further processing.

### Translating Ribosome Affinity Purification (TRAP) and RNA isolation

Frozen tissues (25-100 mg) were pulverized using a spring-loaded hammer and added to polysome buffer. Tissue homogenates were subsequently centrifuged to remove cellular debris and clear lysates were incubated with a purified mouse monoclonal antibody against the HA epitope tag (HA.11 clone 16B12, BioLegend; 3µg/400µl lysate) for 1 hour at 4°C after which magnetic protein G beads (New England BioLabs; 100µl/400µl lysate) equilibrated in polysome buffer were added for an additional 30 minute incubation. The magnetic beads were subsequently washed 3 times with high salt buffer (50 mM Tris (pH 7.5), 300 mM KCl, 12 mM MgCl2, 1% Igepal CA-630, 1 mM dithiothreitol and 100 µg/mL cycloheximide). RLT buffer (RNeasy Mini Kit, Qiagen) supplemented with 2-mercapto-ethanol (1% vol/vol) was added to the beads and vigorously vortexed to dissociate the mRNA from the HA-tagged polysomes. Total RNA from tissue lysates and TRAP polysomal complexes was isolated using the Qiagen RNeasy Mini Kit, including an on-column DNase digestion step, and quantified with the Quant-iT RiboGreen RNA assay (Thermo Fisher Scientific).

### cDNA conversion and RNASeq library preparation

RNA from total tissue lysates (5 ng) or TRAP-selected polysomes (10 ng) was converted into cDNA using the SMARTer Ultra Low Input kit, followed by amplification with the accompanying Advantage 2 PCR kit (Clontech) according to the manufacturer’s protocol. The resulting cDNA was bead-purified (Agencourt AMPure XP beads, Beckman Coulter) and mechanically sheared to a 200-250 base pair range (Bioruptor, Diagenode). End repair with T4 DNA polymerase was followed by dA-tailing using Klenow 3’ to 5’ exonuclease (New England BioLabs) and adenosine triphosphate to create a 3’ adenosine overhang to promote adapter ligation (NEXTflex multiplex adapters, Bioo Scientific). After purification (MinElute Reaction Cleanup kit, Qiagen), samples were linearized by PCR before being size-selected in a range of 300-350 bp using Invitrogen’s E-Gel system. Final enrichment of the adapter-modified cDNA fragments was performed by PCR using Phusion High-Fidelity DNA Polymerase (New England BioLabs). Quality of the final libraries was assessed on the Agilent BioAnalyzer and quantified using the Kapa Library quantification kit (Kapa Biosystems). Flow cells were prepared with 8 samples being multiplexed per lane and sequenced as 50 nucleotide reads on the Illumina HiSeq 2500.

### RNASeq data analysis

Demultiplexed fastq files were aligned against the mouse Gencode reference (GRCm38.p2, Wellcome Trust Sanger Institute) (10) using Bowtie2 (11) and quantified with rSeq(12). Given their separate translational machinery (13), genes encoded by mitochondrial DNA are not associated with HA-tagged RPL22-containing polysomes, which was confirmed by reduced abundance of these genes after TRAP (Supplemental Figure S5). Thus, to avoid a biased inflation of the TRAP-selected transcripts, mitochondrial genes were excluded from analysis. Quantified sequencing reads were normalized to transcripts per million mapped reads (TPM) and transcripts with an average TPM >1 after TRAP were used for further analyses. Differential gene expression analysis to calculate enrichment scores (log2 fold changes) based on transcript abundance in the total tissue lysate and HA-selected fraction, or between experimental conditions, were performed in R (version 3.5.2) where *p*-values were calculated using a Welch’s t-test and adjusted *p*-values were determined based on the Benjamini-Hochberg method, with a false discovery rate (FDR) <10% considered to be significant. Significant differentially expressed transcripts were subjected to gene ontology (GO) (14, 15) analyses using the online STRING (Search Tool for the Retrieval of Interacting Genes/Proteins) biological database version 11.0 (16).

### Classification of gene enrichment categories and EC translatomes

Transcripts significantly depleted (enrichment score <0) after TRAP were classified as parenchymal genes, whereas transcripts significantly more abundant (enrichment score >0) were labeled as *Tek*-enriched (Supplemental Figure 6A). However, *Tek* is also transiently expressed in hematopoietic cells, which may lead to the identification of false positives due to blood contamination of tissues. Therefore, *Tek*-enriched genes were additionally filtered based on data obtained from EC-TRAP on tissues collected from an *Rpl22^fl/fl^, Tek-Cre^+/0^* recipient following bone marrow transplantation using a *Rpl22^fl/fl^, Tek-Cre^−^* donor, as well as TRAP performed on peripheral blood samples from *Rpl22^fl/fl^, Tek-Cre^+/0^* to identify genes that were enriched in the EC compartment only (Supplemental Figure 6A). Of note, this stringent filtering approach removes a number of established EC markers known to be expressed by megakaryocytes/platelets (17), which were also identified by TRAP on platelets from *Rpl22^fl/fl^, Pf4Cre^+/0^* mice (Supplemental Table S6).

Translatome profiles of *Tek^+^* cells are defined as all mRNAs with an averaged TPM value >1 after TRAP, excluding significantly depleted parenchymal genes per tissue. Additional exclusion of genes primarily originating from hematopoietic cells identified by the bone marrow transplant data resulted in the identification of tissue-specific EC translatomes (Supplemental Figure S6b).

### Primary EC isolation and microarray analysis

ECs from heart capillaries and kidney cortex peritubular capillaries were isolated as previously described (18). In summary, tissues from 4-6 C57BL/6 mice (2-4 weeks old) were pooled, minced and digested with type I collagenase at 37°C for 30 minutes, followed by separation through a 70 µm cell strainer. Cells were washed and incubated with rat anti-mouse CD31 antibody (BD Pharmingen) and CD31-positive cells were collected and either immediately processed for microarray analysis, or cultured in DMEM containing 15% fetal bovine serum and endothelial mitogen BT-203 (Biomedical Technologies) for 3 days before RNA isolation for microarray analysis using the Illumina MouseRef-8 Expression BeadChip. Quantile normalization was performed using the GeneSpring GX software (version 11.5.1; Agilent) and additional processing and differential expression analyses were performed using the R packages lumi (19) (version 2.34.0) and limma (20) (version 3.38.3). Adjusted *p*-values were determined based on the Benjamini-Hochberg method where an FDR <10% for differentially expressed genes was considered to be significant.

### Tissue specific endothelial cell content

High throughput DNA sequencing of floxed and *Cre*-excised *Rpl22* isoforms was performed to determine the *Tek*-positive cell content within multiple tissues. *Rpl22^fl/fl^,Tek-Cre^+/0^* males were perfused with PBS and tissues were harvested, followed by DNA isolation (DNeasy Blood and Tissue kit, Qiagen). Intron 3 of *Rpl22*, spanning either the LoxP site for the floxed allele or the combined LoxP/FRT site for the *Cre*-excised allele (1) was PCR amplified using the following primers: 5’-CACCTGTGCGGTCTTTCTCT-3’ and 5’ GCAAGCAAGCGTCTATCACA-3’ (Figure S7). After PCR purification (QIAquick PCR purification kit, Qiagen) libraries were prepared for multiplexed high throughput sequencing as above and sequenced on the Illumina MiSeq as 75 nucleotide paired-end reads. Raw reads were aligned to the genomic *Rpl22* isoforms and the fraction *Cre*-excised sequences of the total number of *Rpl22* reads, indicating the *Tek*-positive cell content, was used as a proxy for the fraction of ECs assuming excision occurred in both alleles per cell.

### Histology

*Rosa^mT/mG^,Tek-Cre^+/0^* mice were anesthetized and perfused with 4% paraformaldehyde in PBS. Tissues were isolated and post-fixed overnight at 4°C in 4% paraformaldehyde, followed by overnight cryoprotection in 30% sucrose (4°C) and embedded in optimal cutting temperature medium (OCT, Sakura). Ten micron cryosections were mounted with Prolong Gold antifade reagent including DAPI (4’,6-diamidino-2-phenylindole; Thermo Fischer Scientific) and imaged on a Nikon A1 Confocal microscope using the accompanying Elements Viewer software.

### Single-molecule fluorescent in situ RNA hybridization

Mice were perfused with 12 ml PBS and tissues were immediately embedded in OCT compound. Twelve micron sections were mounted on SuperFrost Plus slides (Thermo Fisher) and dried overnight at −20°C. After fixing and dehydration, slides were subjected to the RNAscope technology (Advanced Cell Diagnostics) for multiplexed fluorescent *in situ* RNA hybridization, which allows visualization of single mRNA transcripts (21). Sections were treated with Protease IV for 30 minutes at room temperature followed by probe hybridization, followed by signal amplification and counterstaining according to the manufacturer’s protocol. In order to determine endothelial localization, probes were multiplexed with *Tek* (Mm-*Tek*-C1) in either channel 2 (Mm-*Eva1b*-C2, Mm-*3110099E03Rik*-C2 and Mm-*Wnt9b*-C2) or channel 3 (Mm-*Bvht*-C3). Slides were imaged on a Nikon A1 Confocal microscope and analyzed using the Elements Viewer software.

---

**Supplemental Figure S1:**
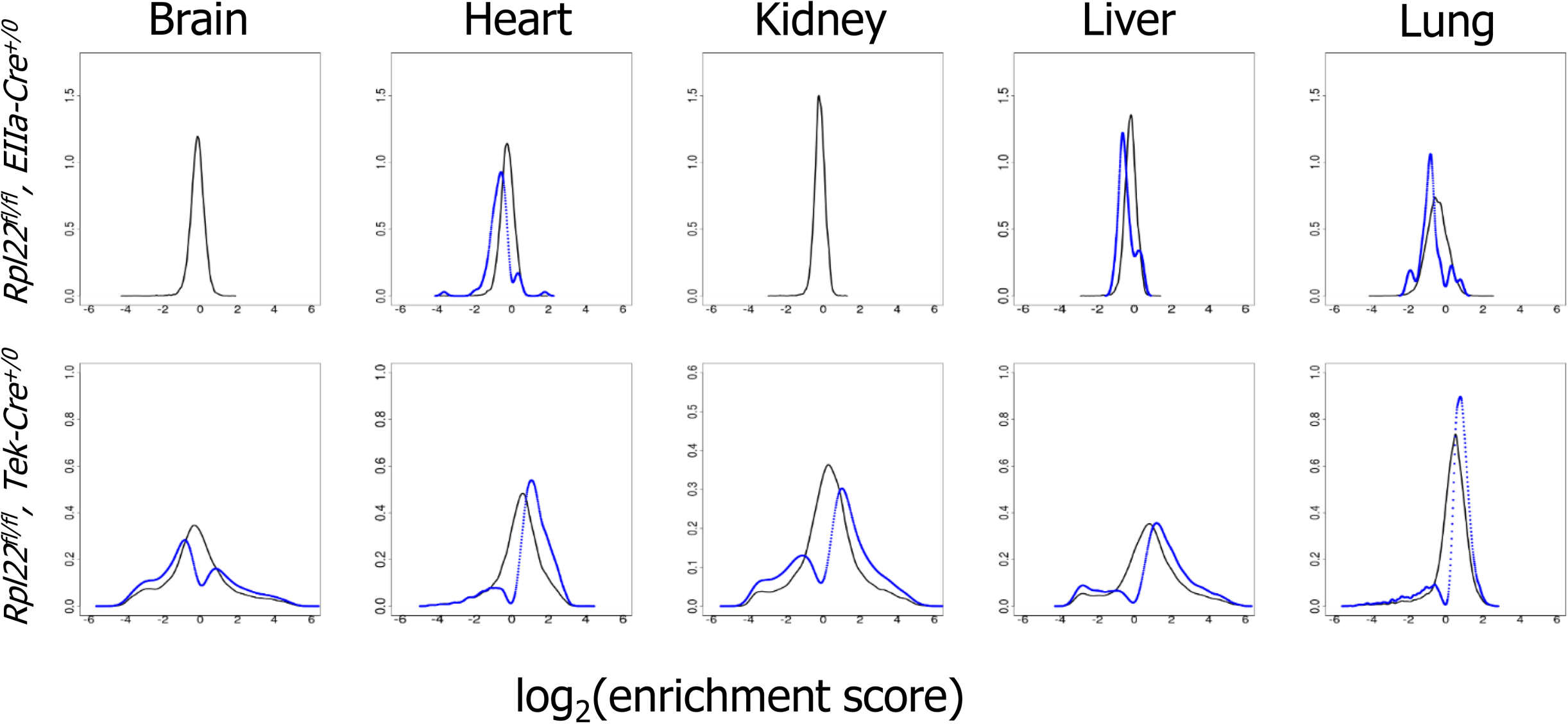
Density plots of enrichment scores for TRAP performed on mice with HA-tagged RPL22 in all cells (*Rpl22^fl/fl^, EIIa-Cre^+/0^*; top panel) show an overall limited dynamic range in the distribution of enrichment scores in tissues (black lines), with few significant differences between the transcriptome and translatome (blue lines, enrichment scores with FDR<10%). EC-TRAP (*Rpl22^fl/fl^, Tek-Cre^+/0^*; bottom panel) identifies genes enriched in ECs (blue line, log_2_ fold change >0) as well as parenchymal genes (log_2_ fold change <0).

**Supplemental Figure S2:**
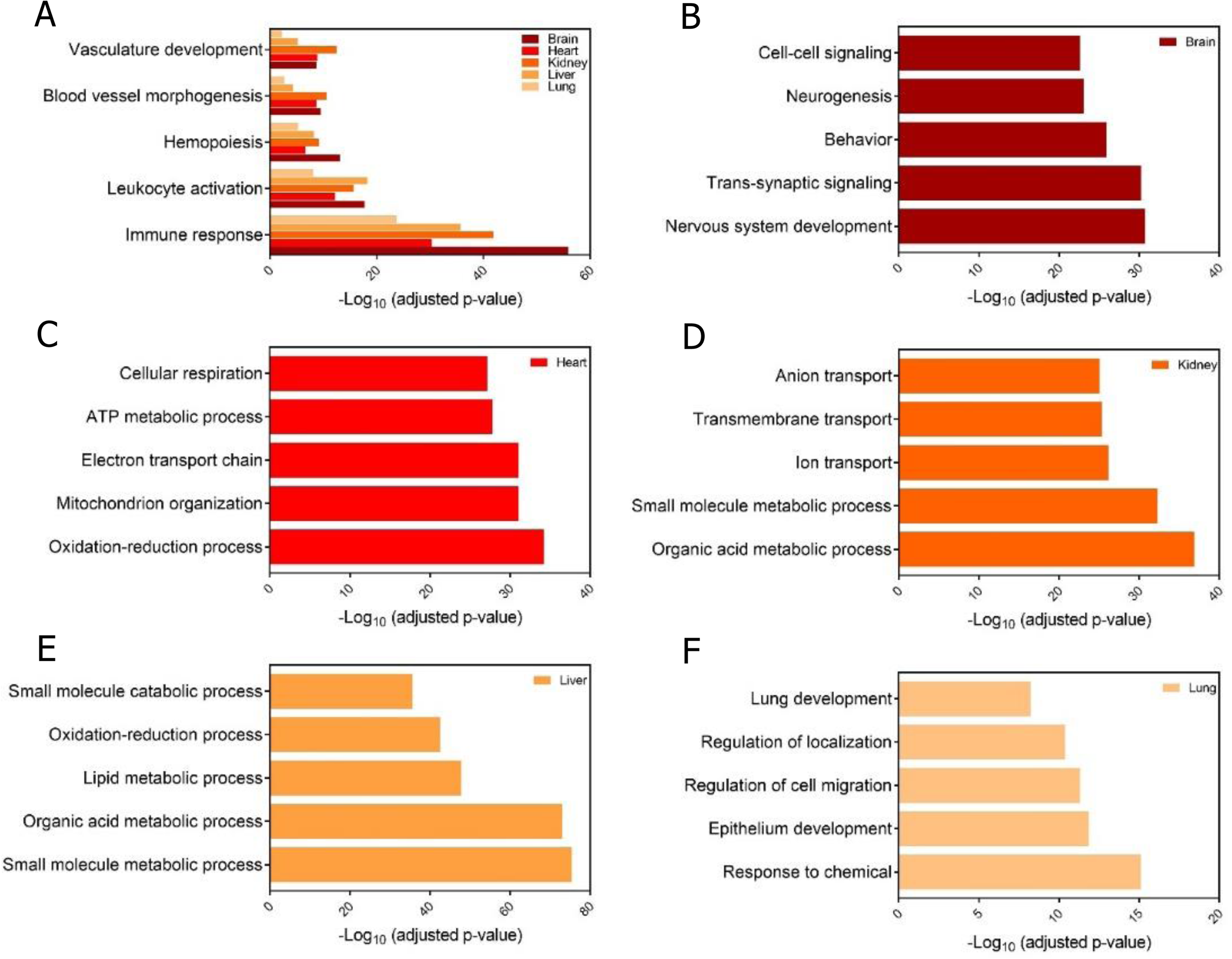
Gene ontology analysis of significantly enriched and depleted genes after TRAP. (A) Top 500 enriched genes after TRAP (FDR<10%, ranked based on enrichment scores) were subjected to GO analysis which identified processes associated with vascular and hematopoietic cells, consistent with the known expression of *Tek*. (B-F) GO analysis on the top 500 most depleted transcripts per tissue identified processes expected to be associated with tissue-specific parenchymal cells, consistent with efficient removal of non-EC mRNAs by the TRAP procedure.

**Supplemental Figure S3:**
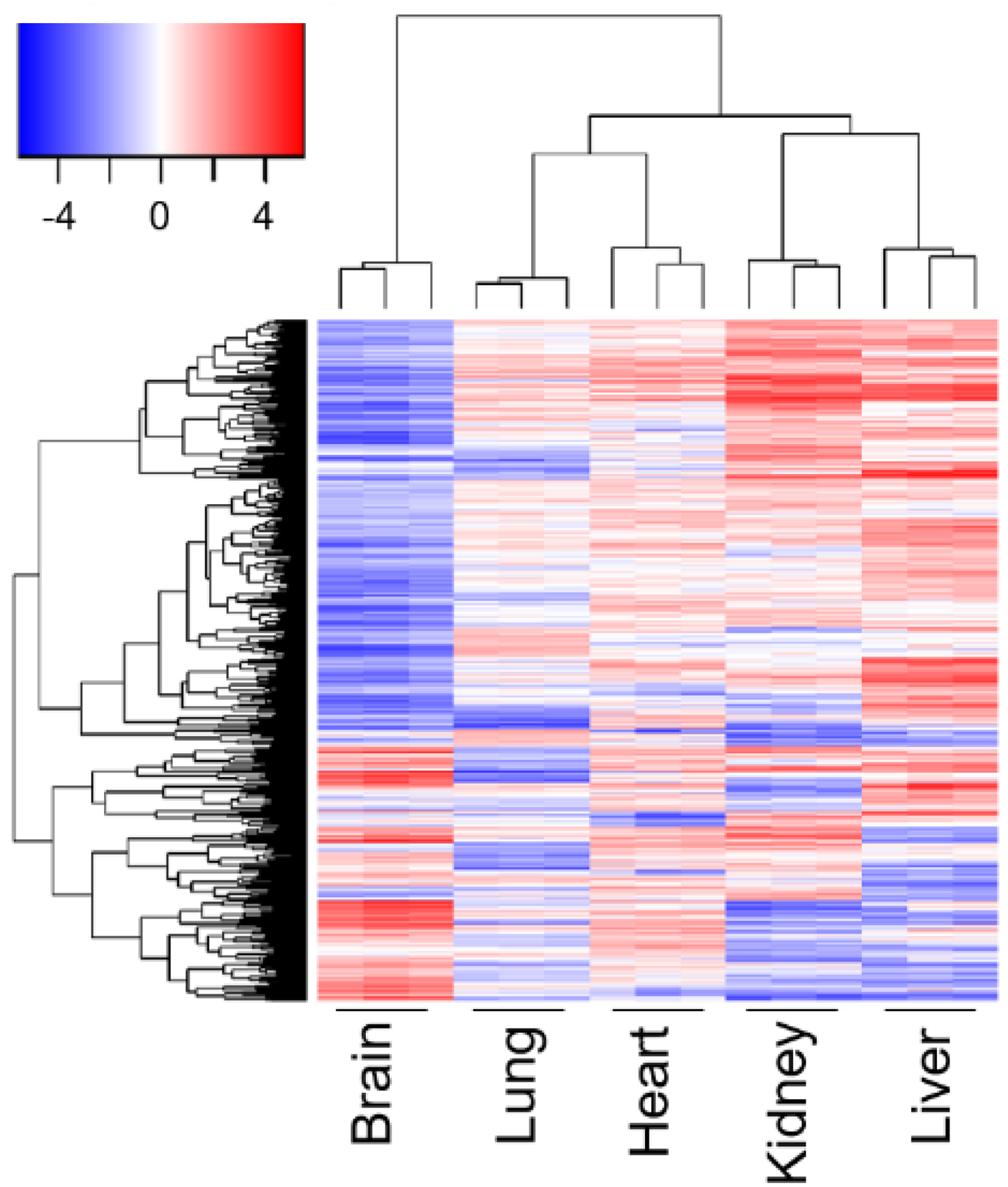
Hierarchical clustering of transcripts identified as EC-enriched in at least 1 tissue (log_2_ fold change >1), while primarily parenchymal-specific in at least 1 other tissue (log_2_ fold change <1). See Supplemental Table S2 for the full gene list and accompanying enrichment scores.

**Supplemental Figure S4:**
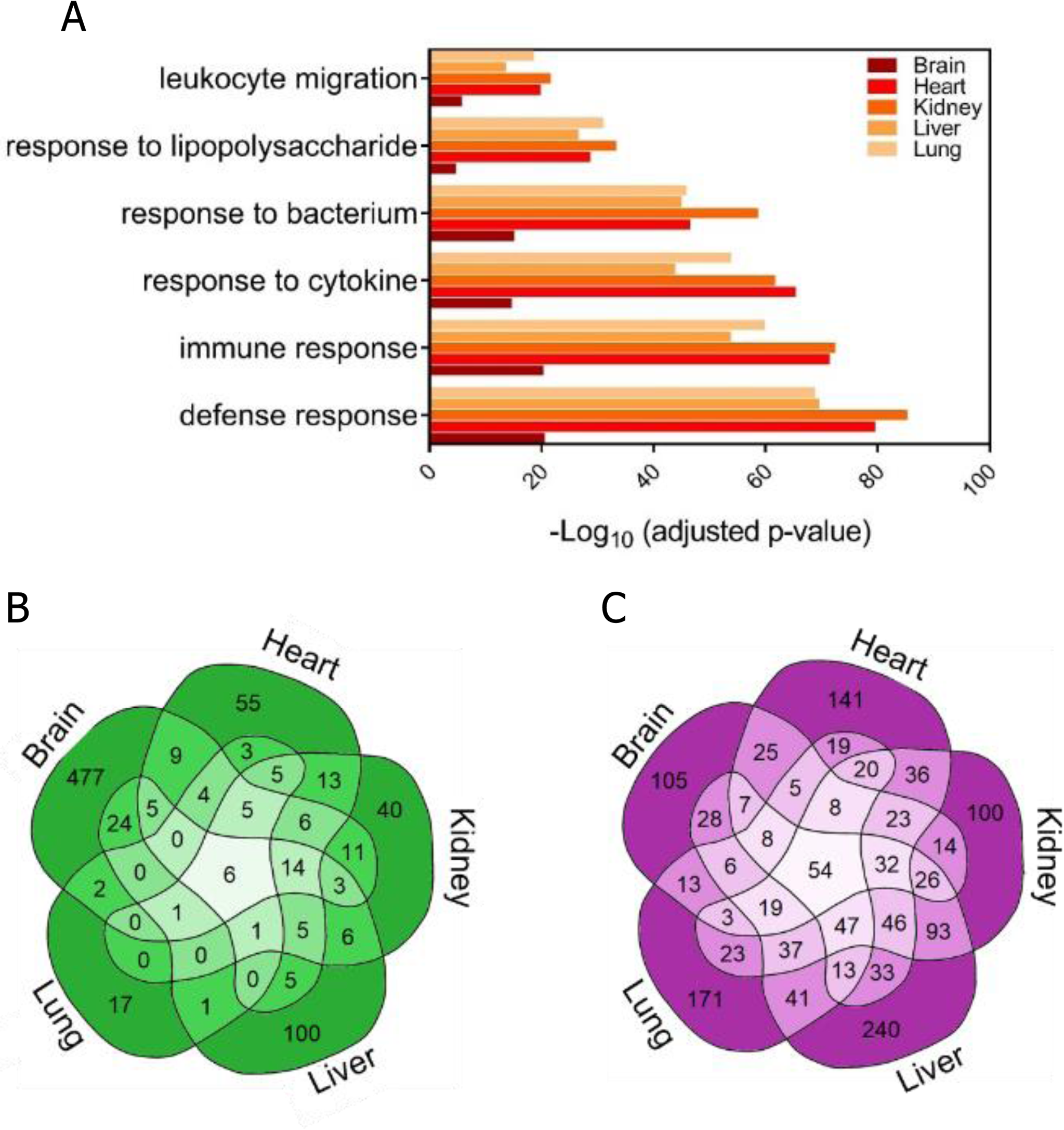
Tissue-specific changes in expression profiles upon LPS exposure. (A) GO analysis of organ-wide transcriptomes following LPS exposure shows upregulation of genes involved in the defense response. (B-C) Venn diagram showing the overlap between EC-enriched genes significantly upregulated (log_2_ fold change >1; C) versus downregulated (log_2_ fold change <-1; D) across tissues after LPS exposure.

**Figure S5:**
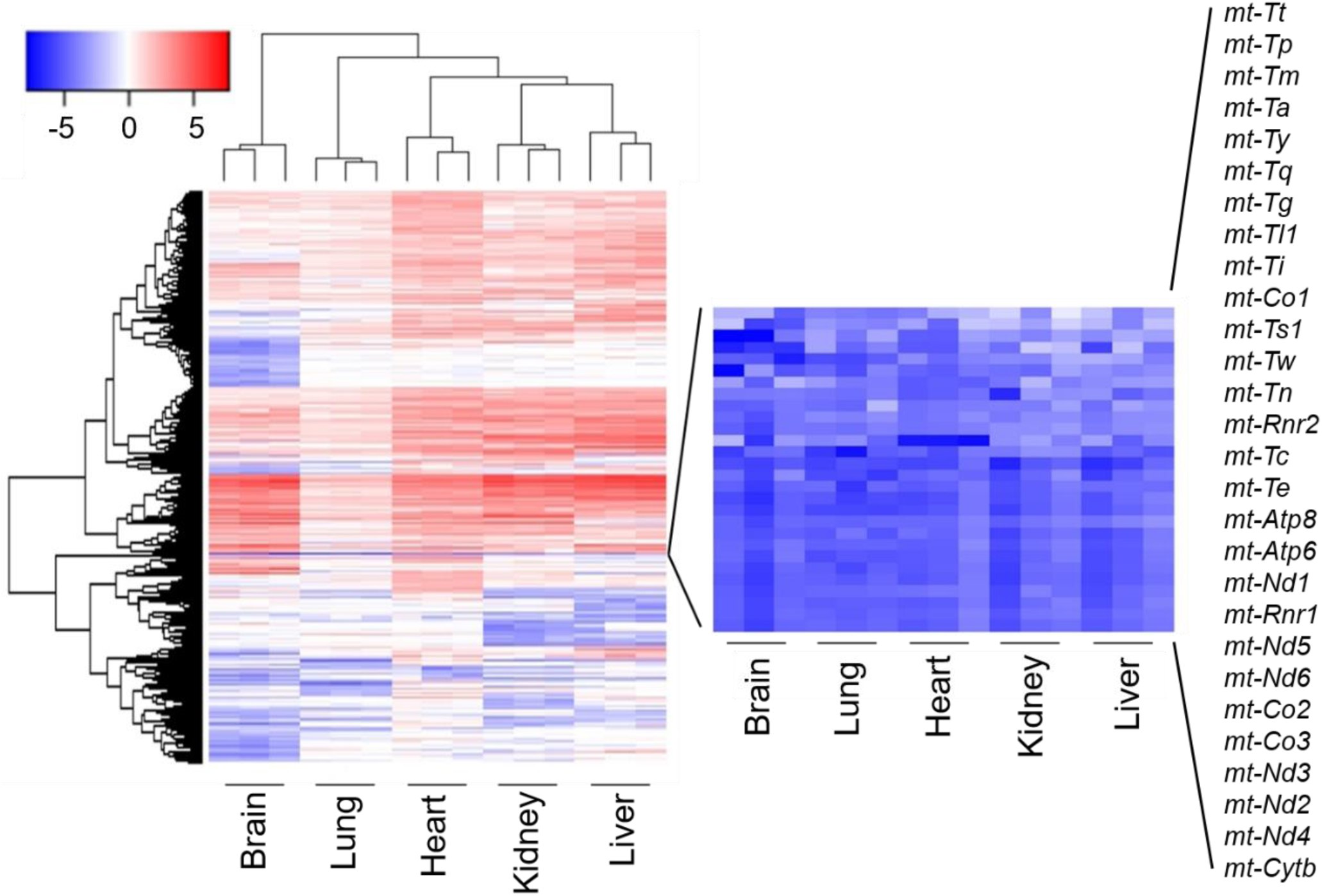
Unsupervised clustering of enrichment scores after EC-TRAP indicates a general and expected depletion of mitochondrial genes as these are not associated with RPL22-containing ribosomes. To avoid bias and inflation of enrichment scores, mitochondrial genes were therefore excluded before performing differential expression analysis after TRAP.

**Figure S6:**
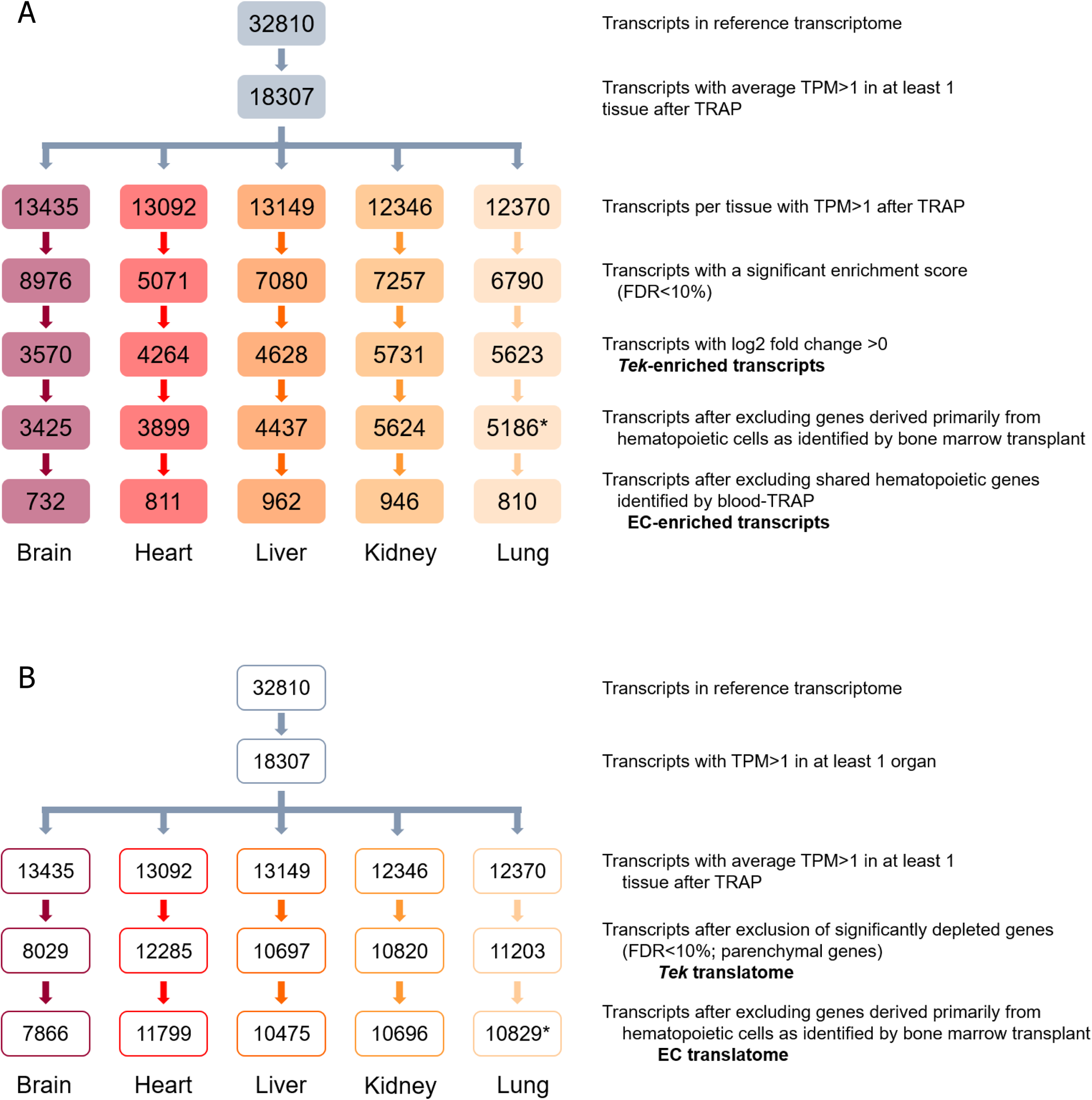
Classification of gene enrichment categories and cell-specific expression profiles. (A) Number of transcripts present after filtering RNASeq data to identify genes that are specifically enriched in ECs, thus excluding genes that are commonly shared with other cell types. (B) Number of all transcripts identified in EC translatomes, which include housekeeping genes as well as transcripts shared, but not primarily derived from, hematopoietic cells. *Due to suboptimal sequencing quality of lung samples obtained from the *Rpl22^fl/fl^, Tek-Cre^+/0^* mouse receiving *Rpl22^fl/fl^, Tek-Cre^−^* bone marrow, hematopoietic transcripts were defined based on the sum of hematopoietic genes identified in brain, heart, kidney and liver.

**Figure S7:**
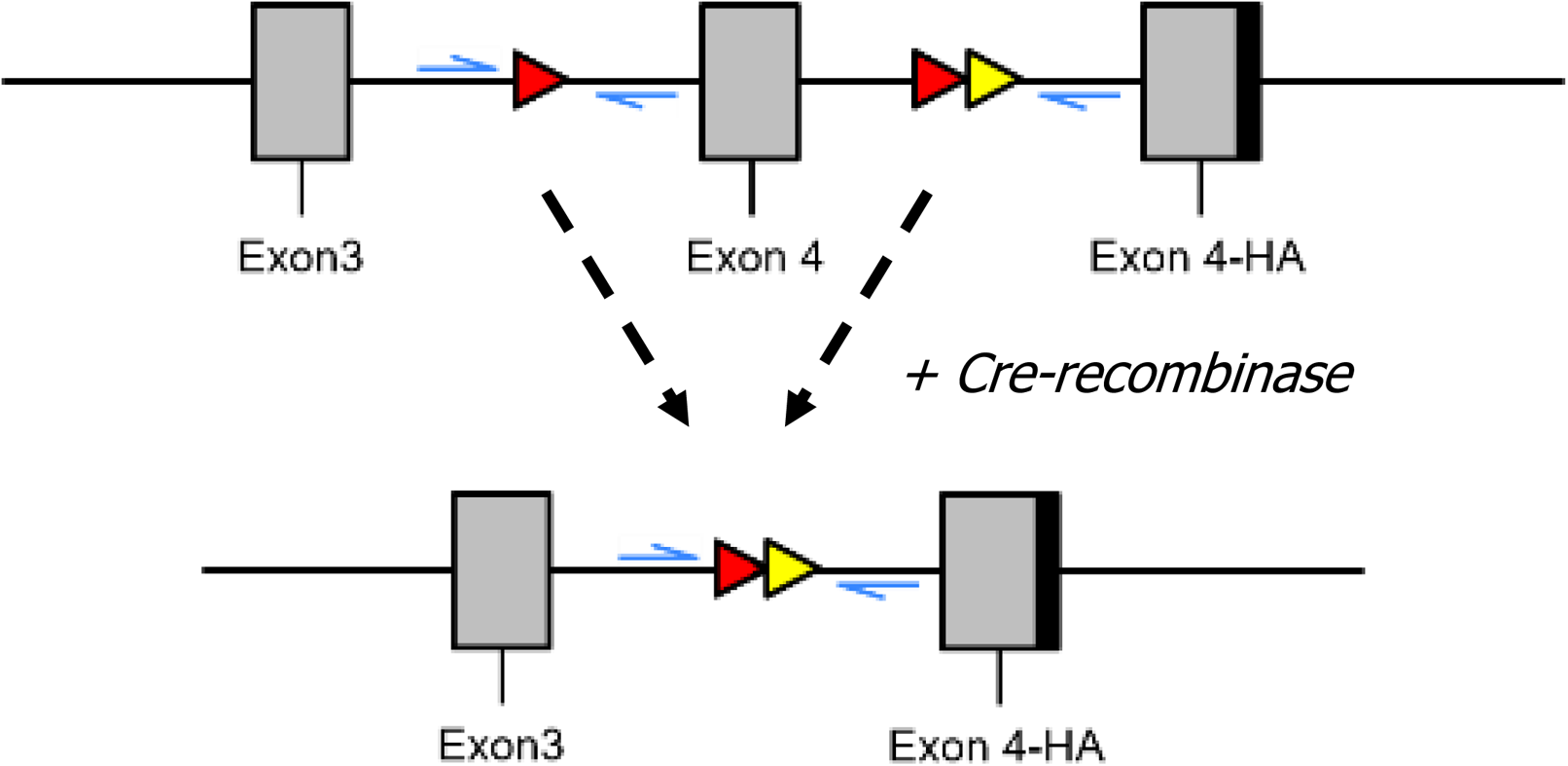
Primer design to identify the tissue-specific EC content based on genomic *Rpl22* isoforms. PCR primers in the floxed allele span the LoxP site (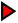) resulting in a 157 basepair amplicon whereas after *Cre-recombinase* and excision of the wildtype exon 4, primers will span both the loxP and FRT site (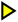) leading to a 240 basepair amplicon. Amplicons were sequenced as paired-end 75 nucleotide reads, allowing the discrimination of the HA-expressing allele from the total amplicons sequenced.

**Supplemental Table S1:**
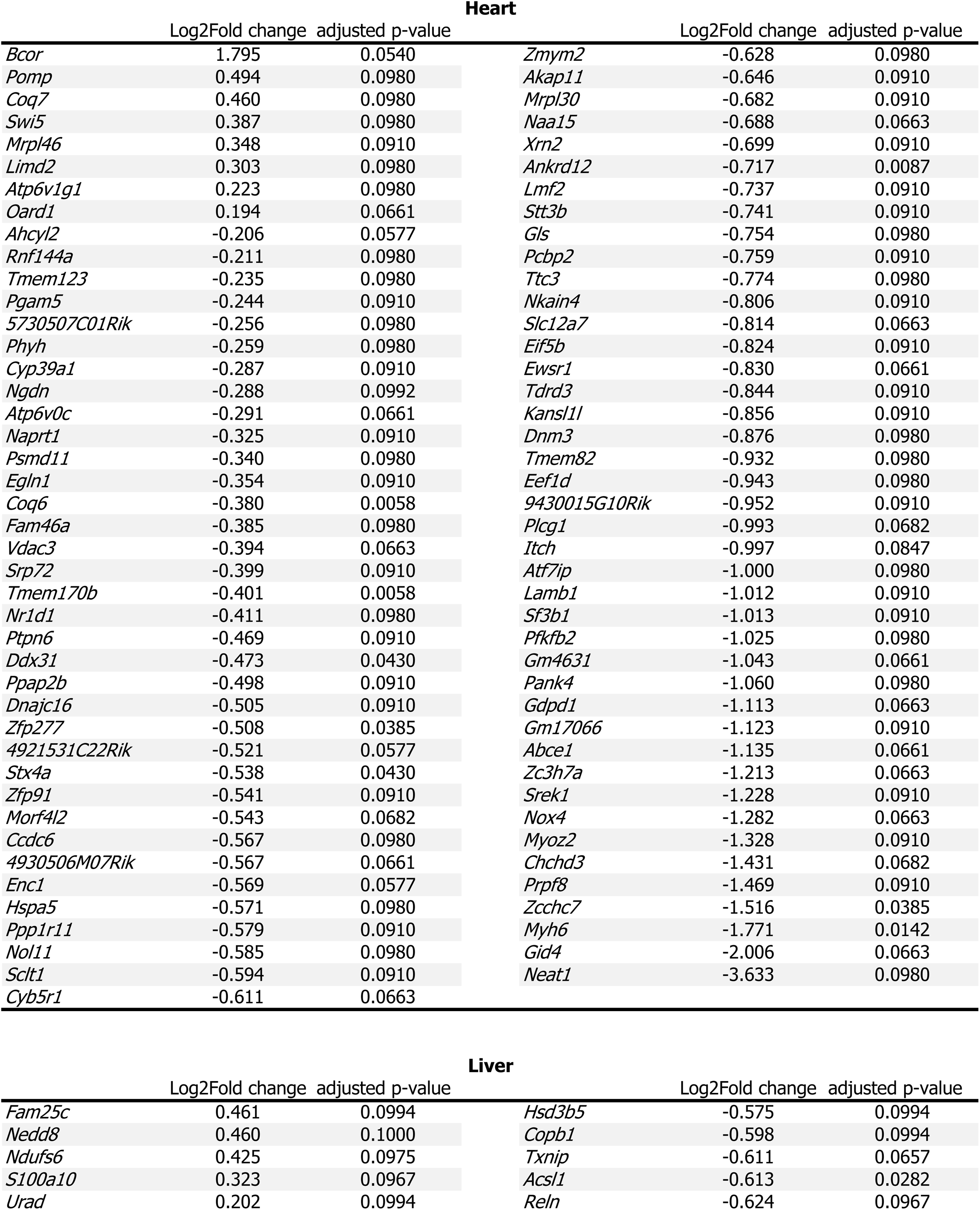
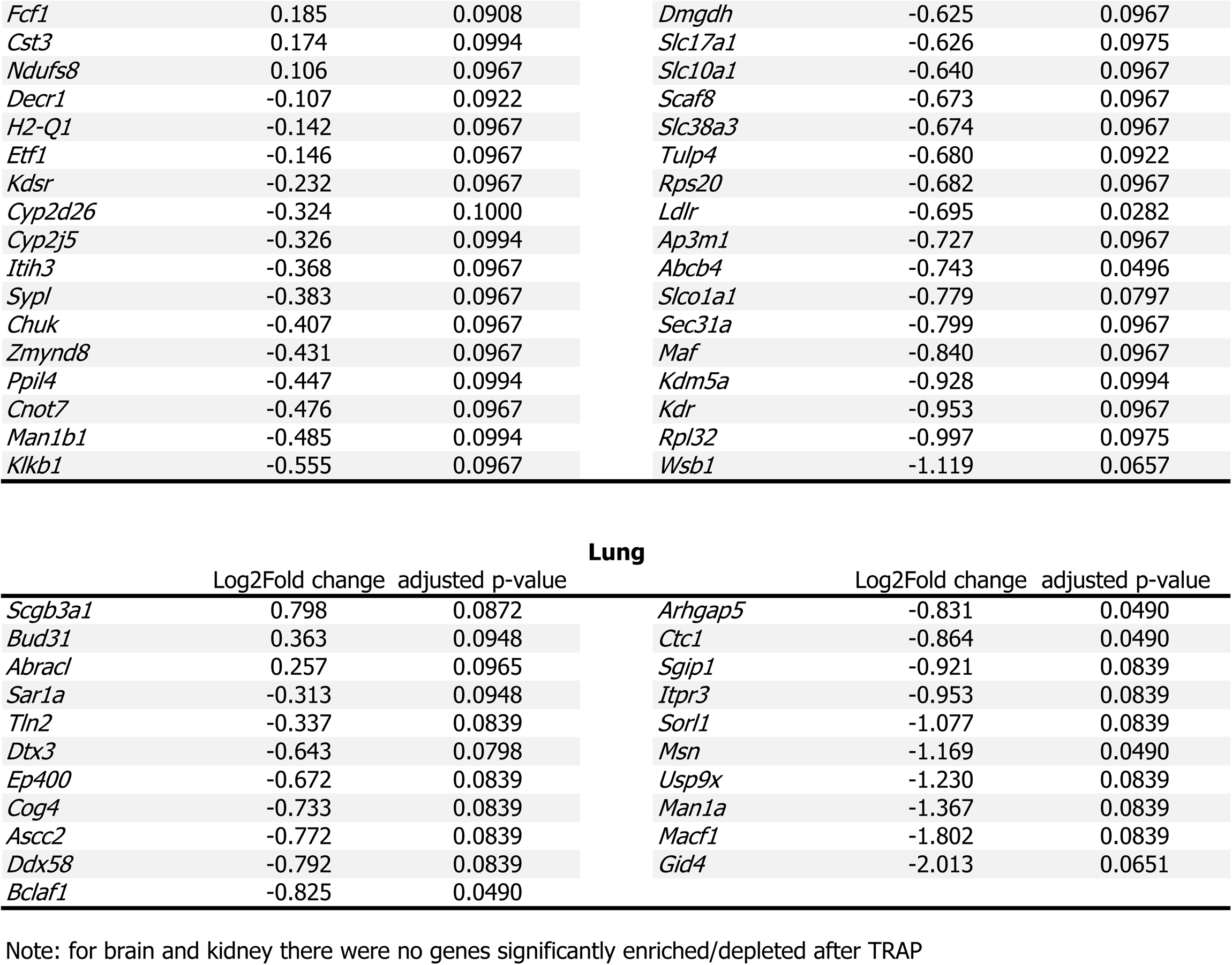
Enrichment scores of genes significantly enriched or depleted in mice ubiquitously expression HA-tagged Rpl22.

**Supplemental Table S2:**
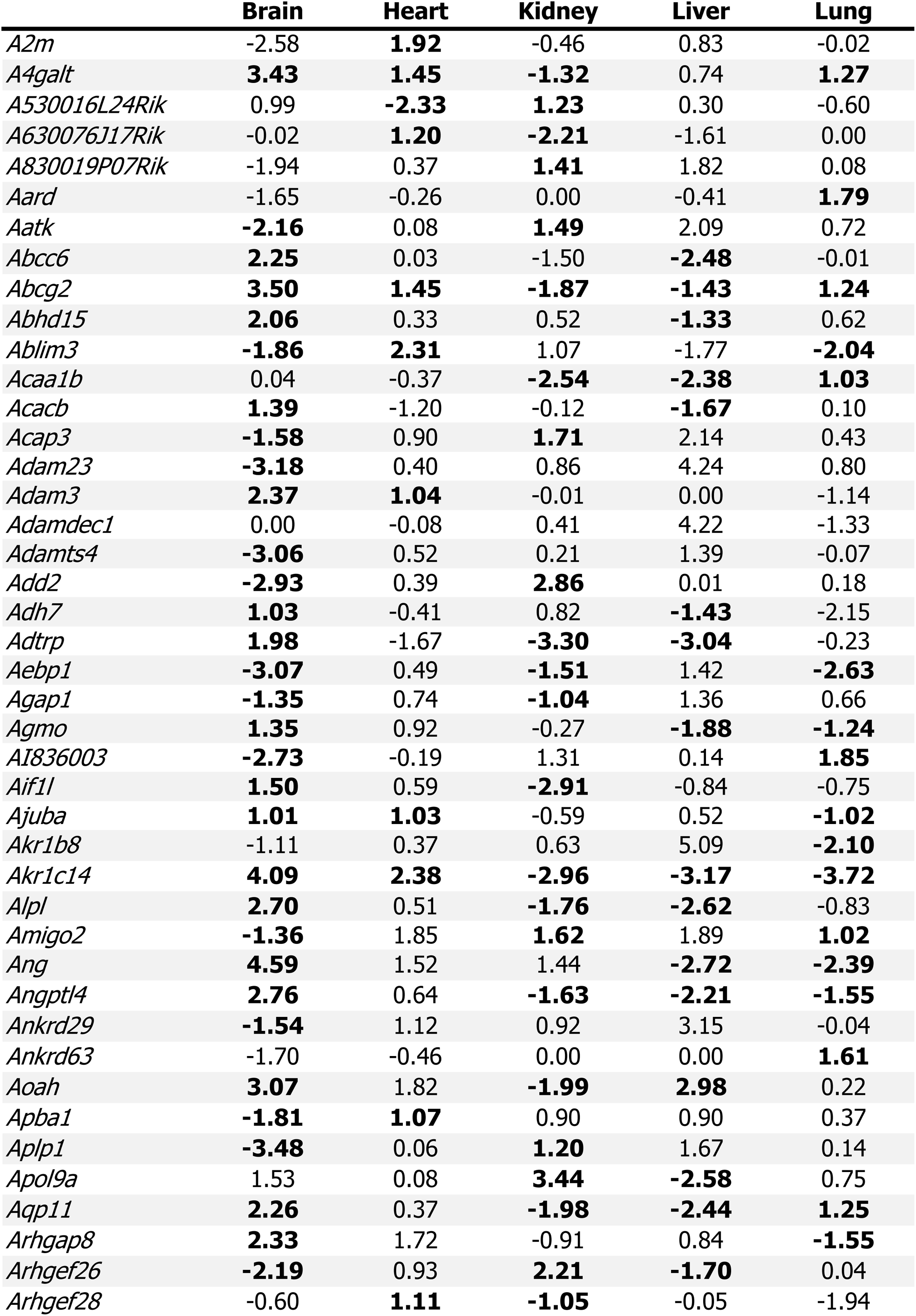
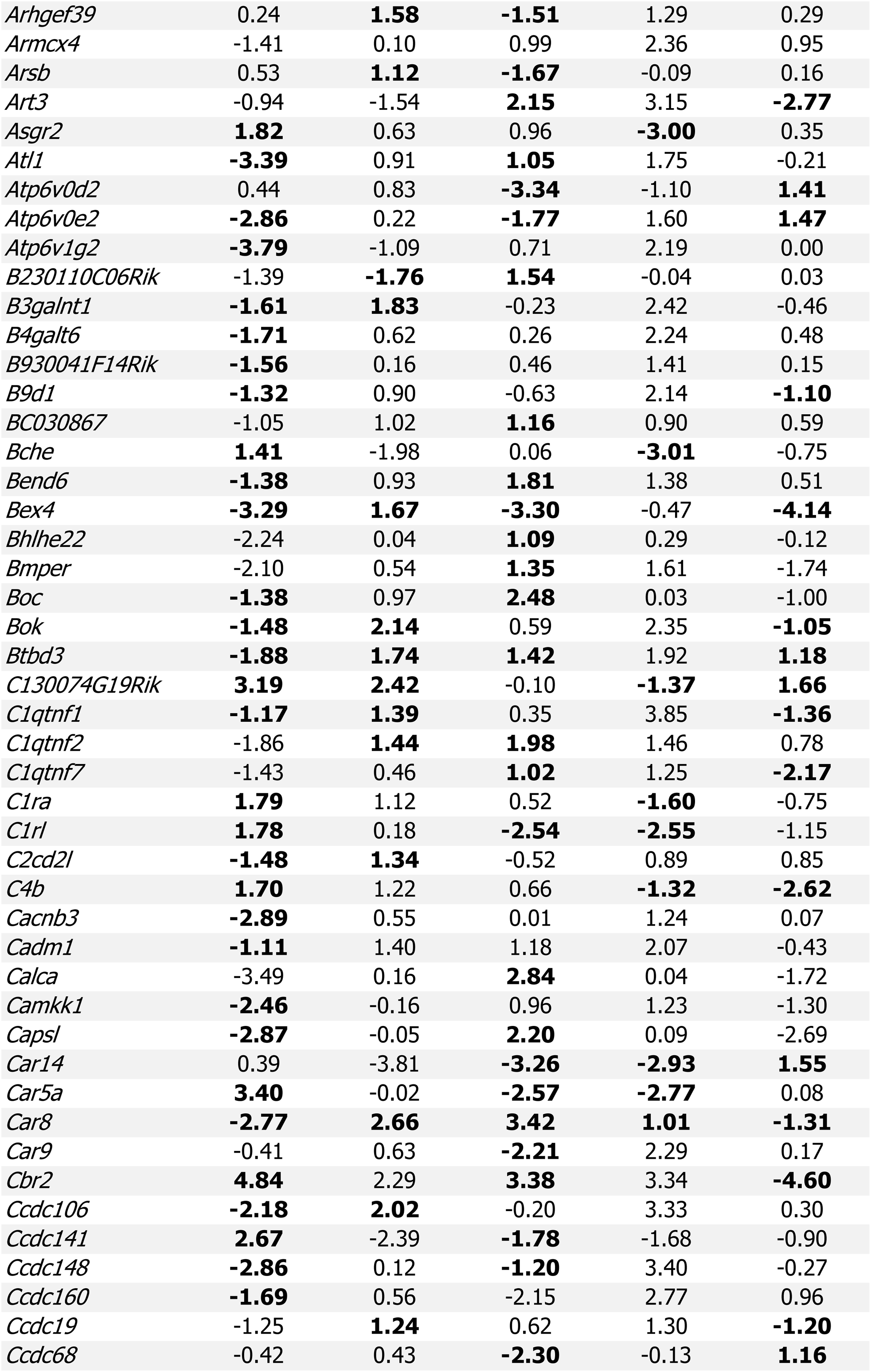
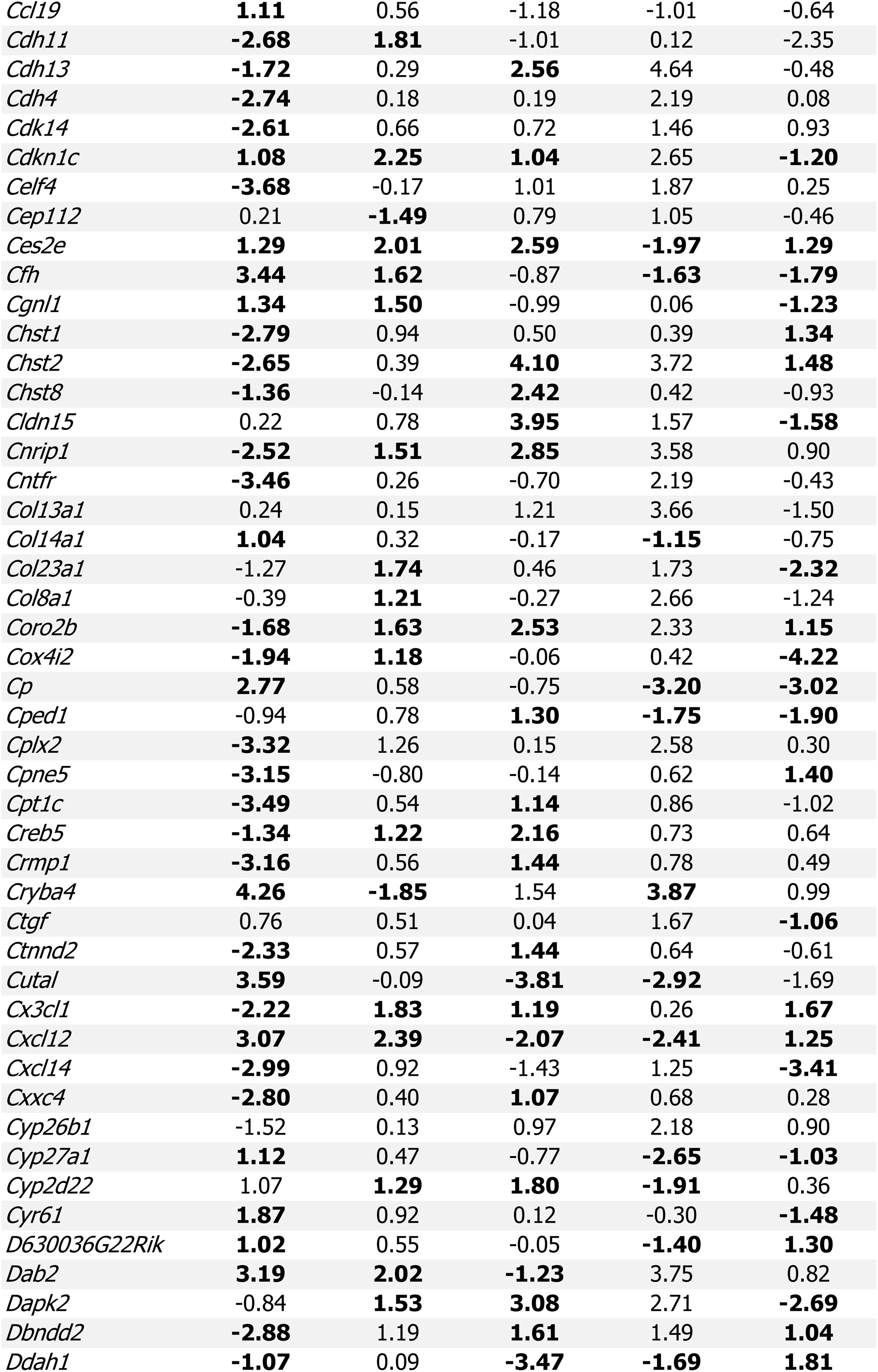
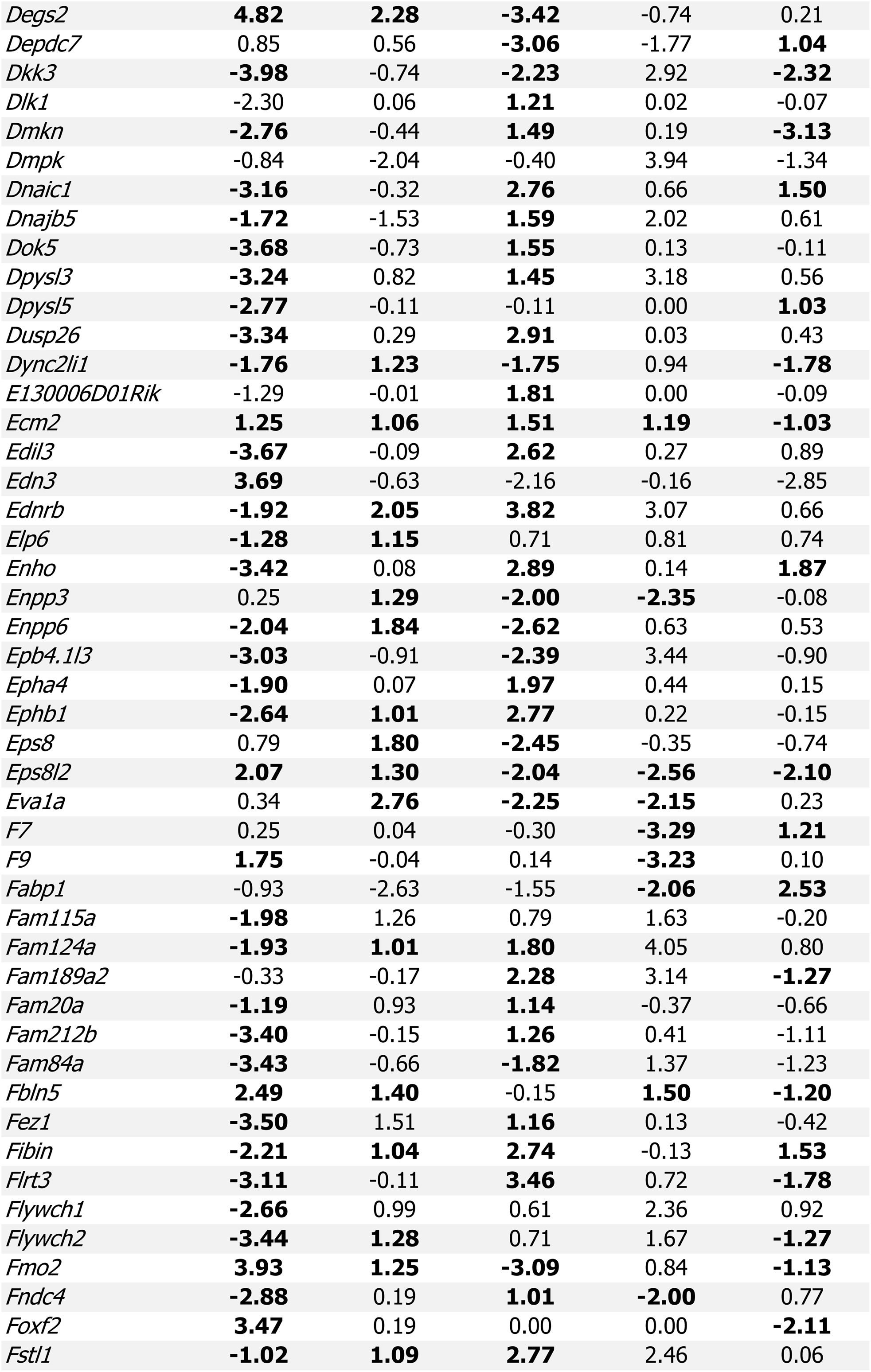
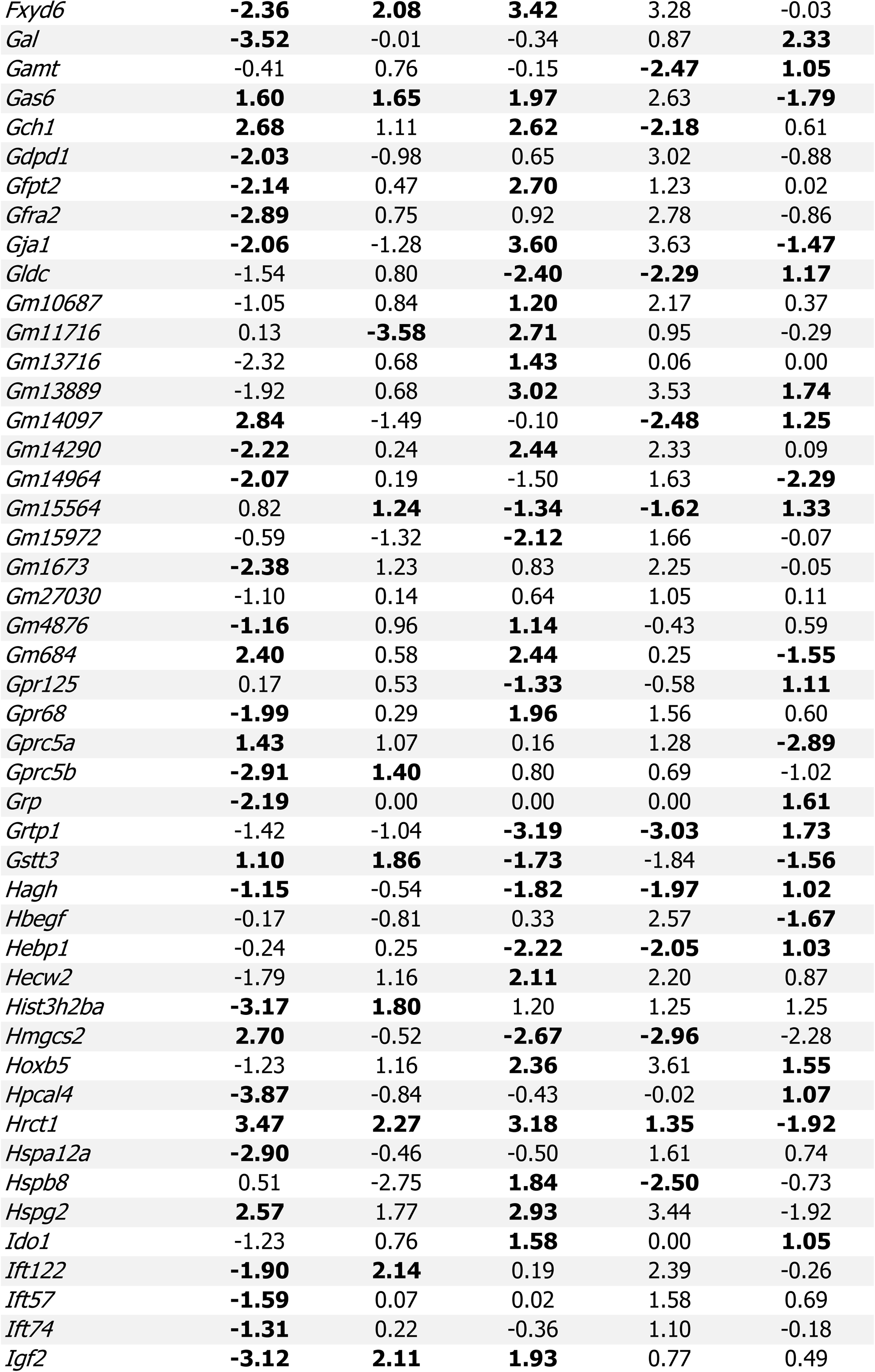
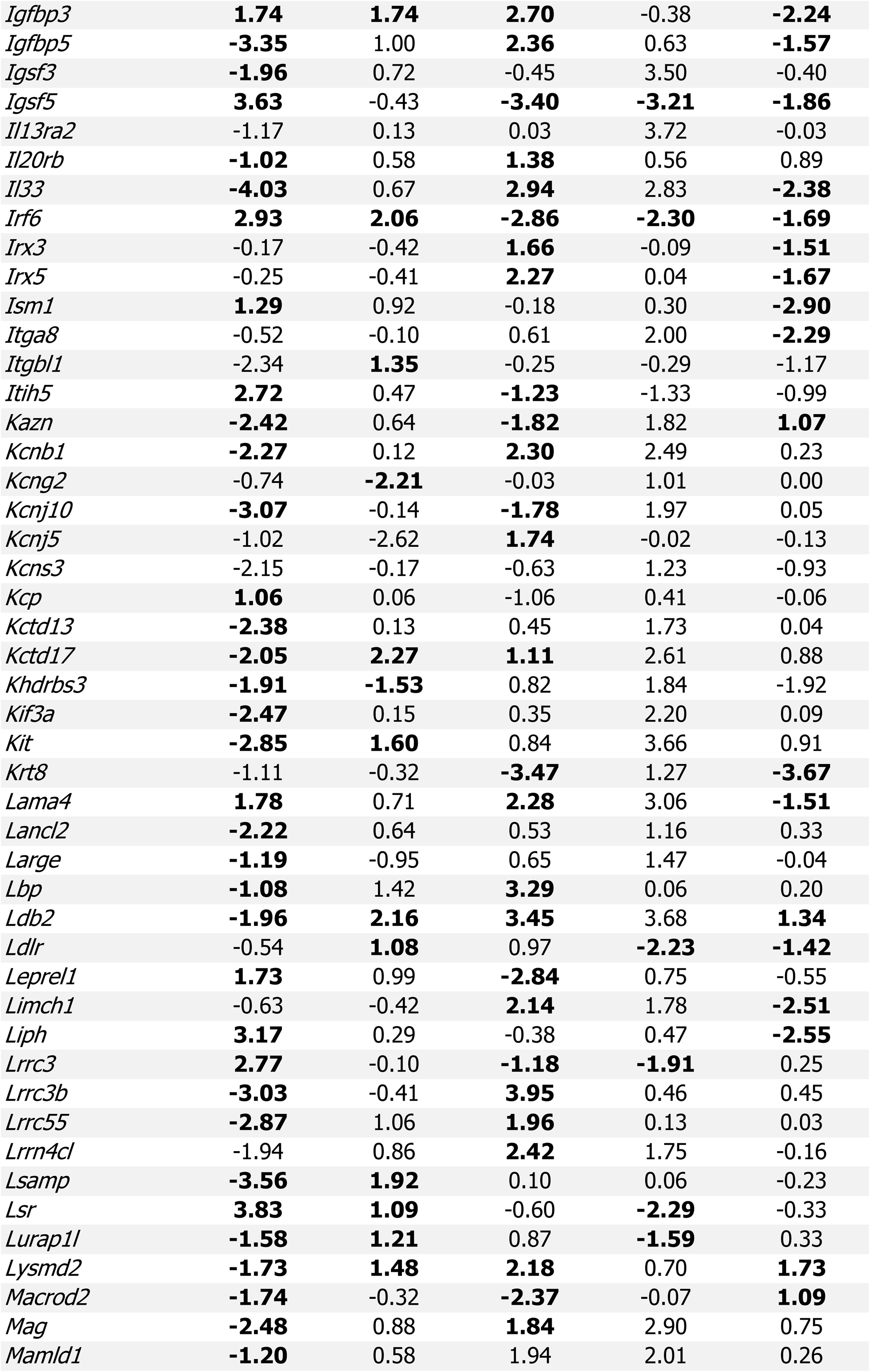
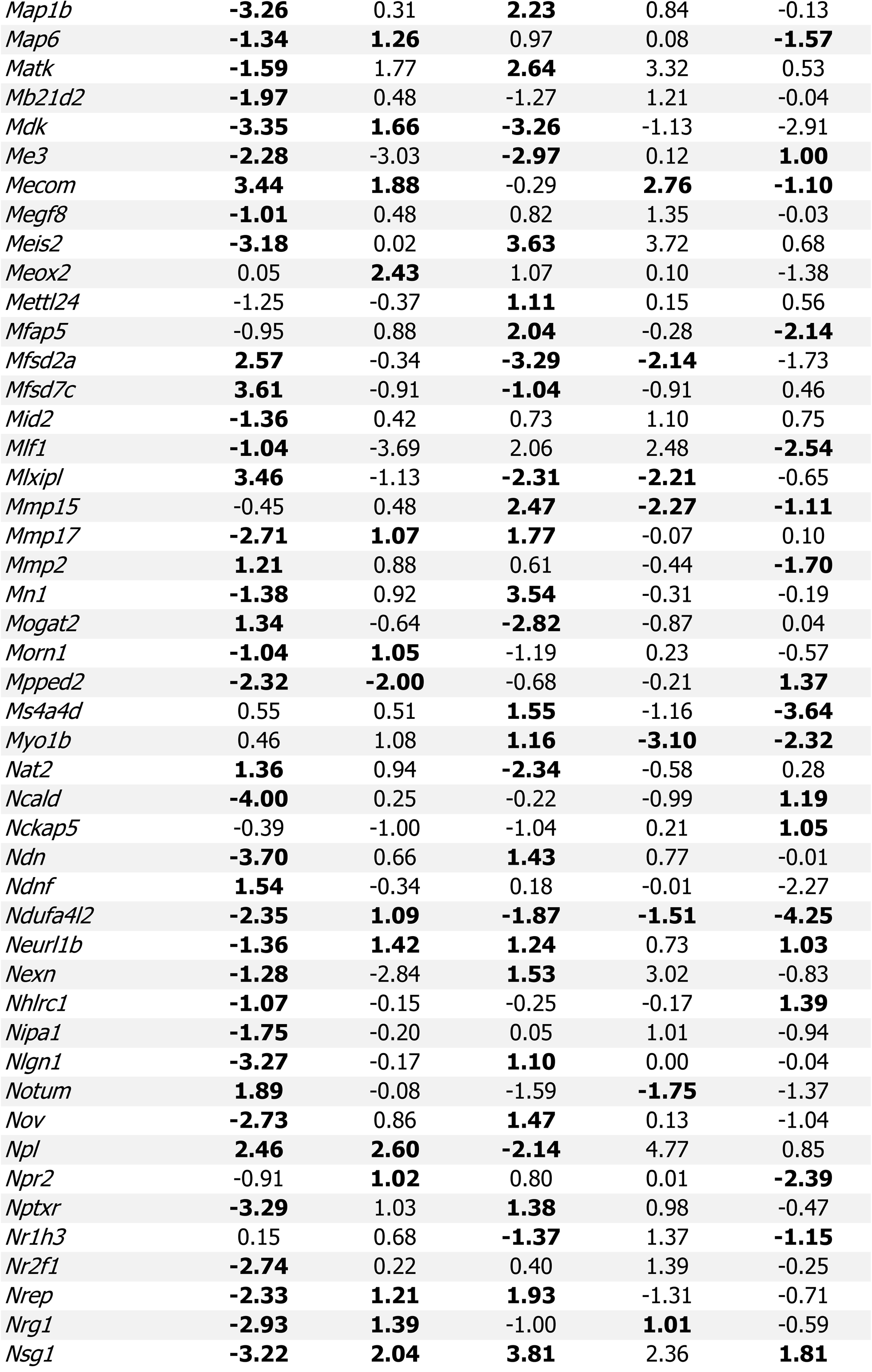
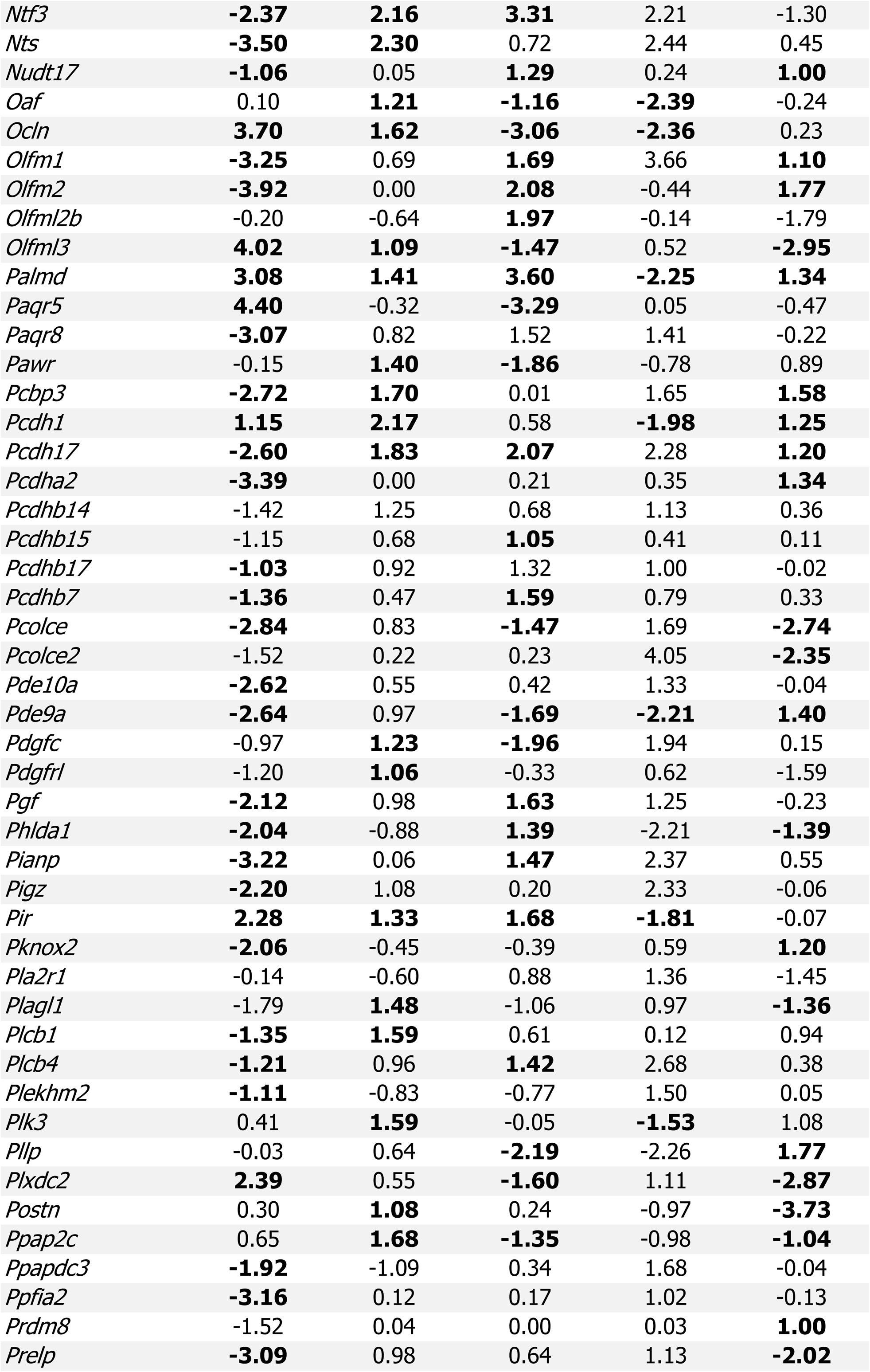
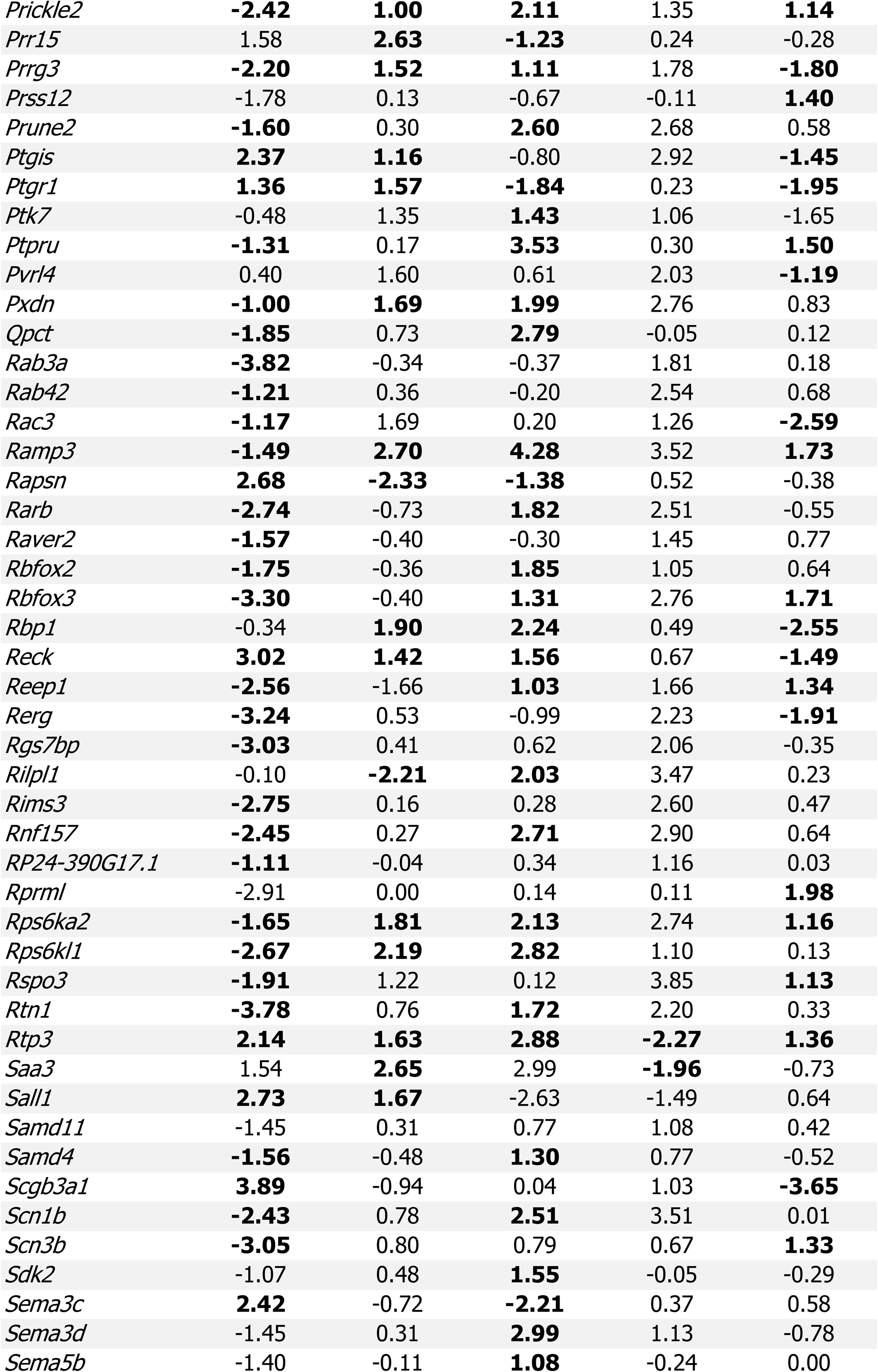
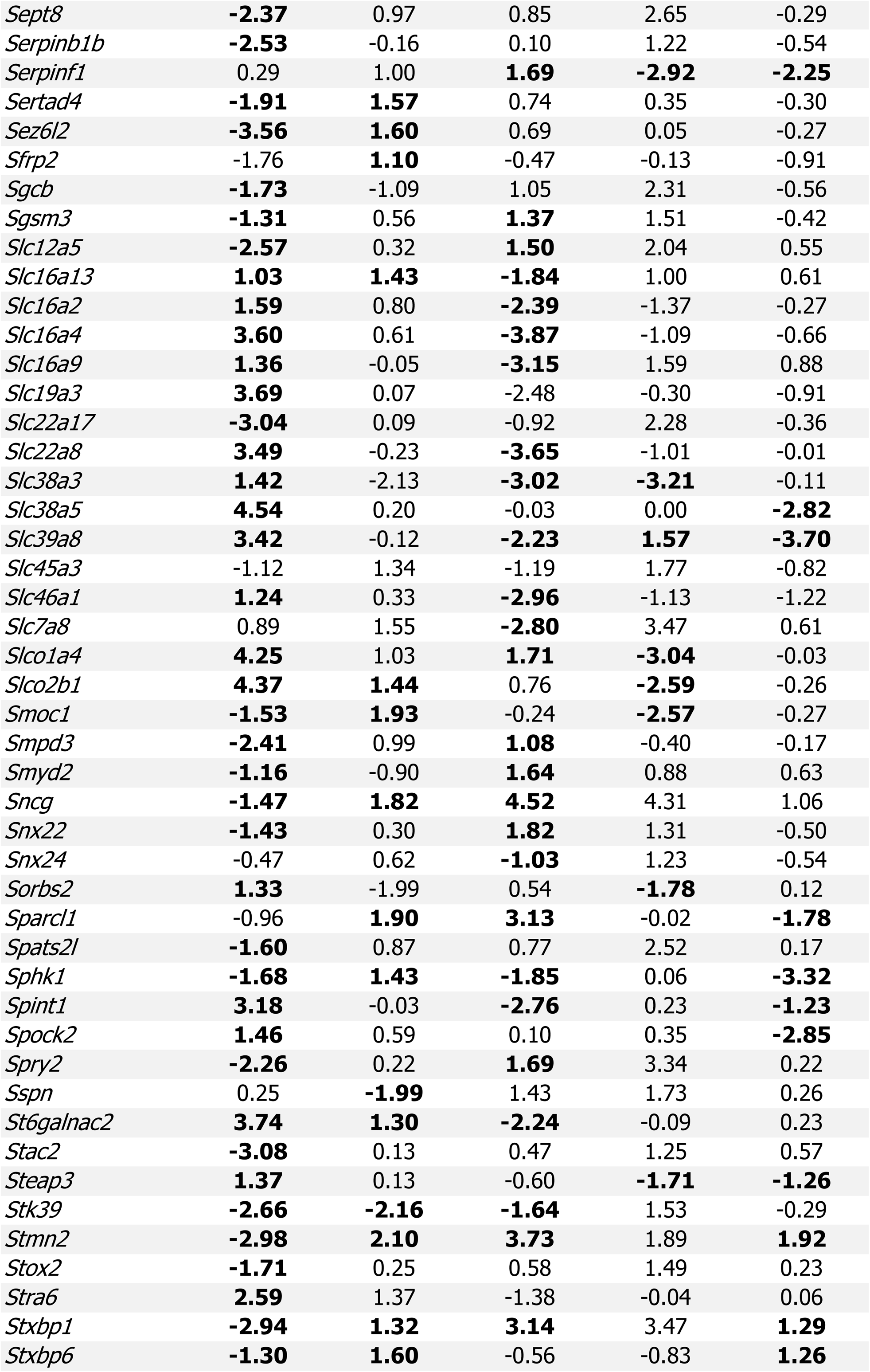
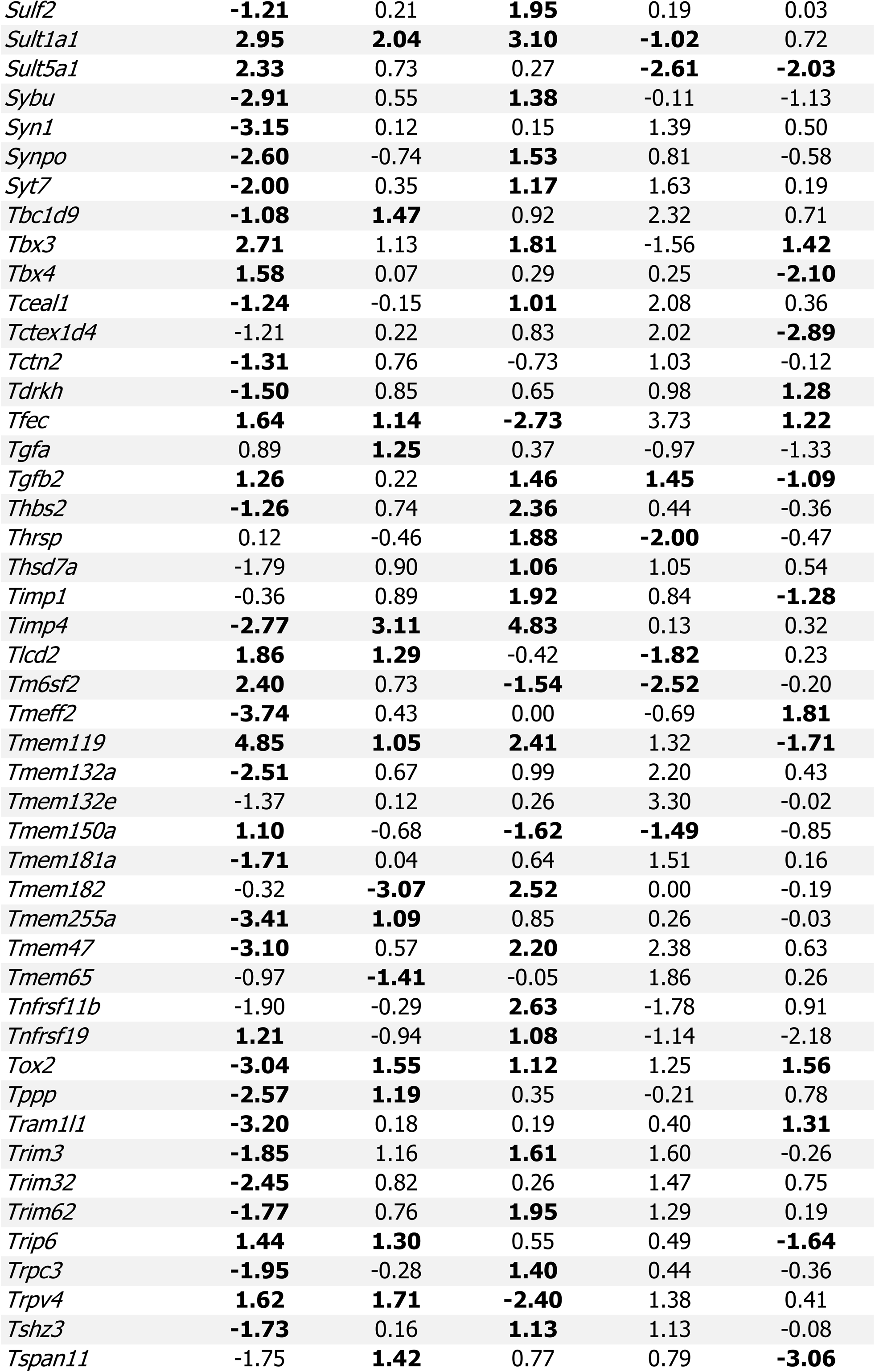
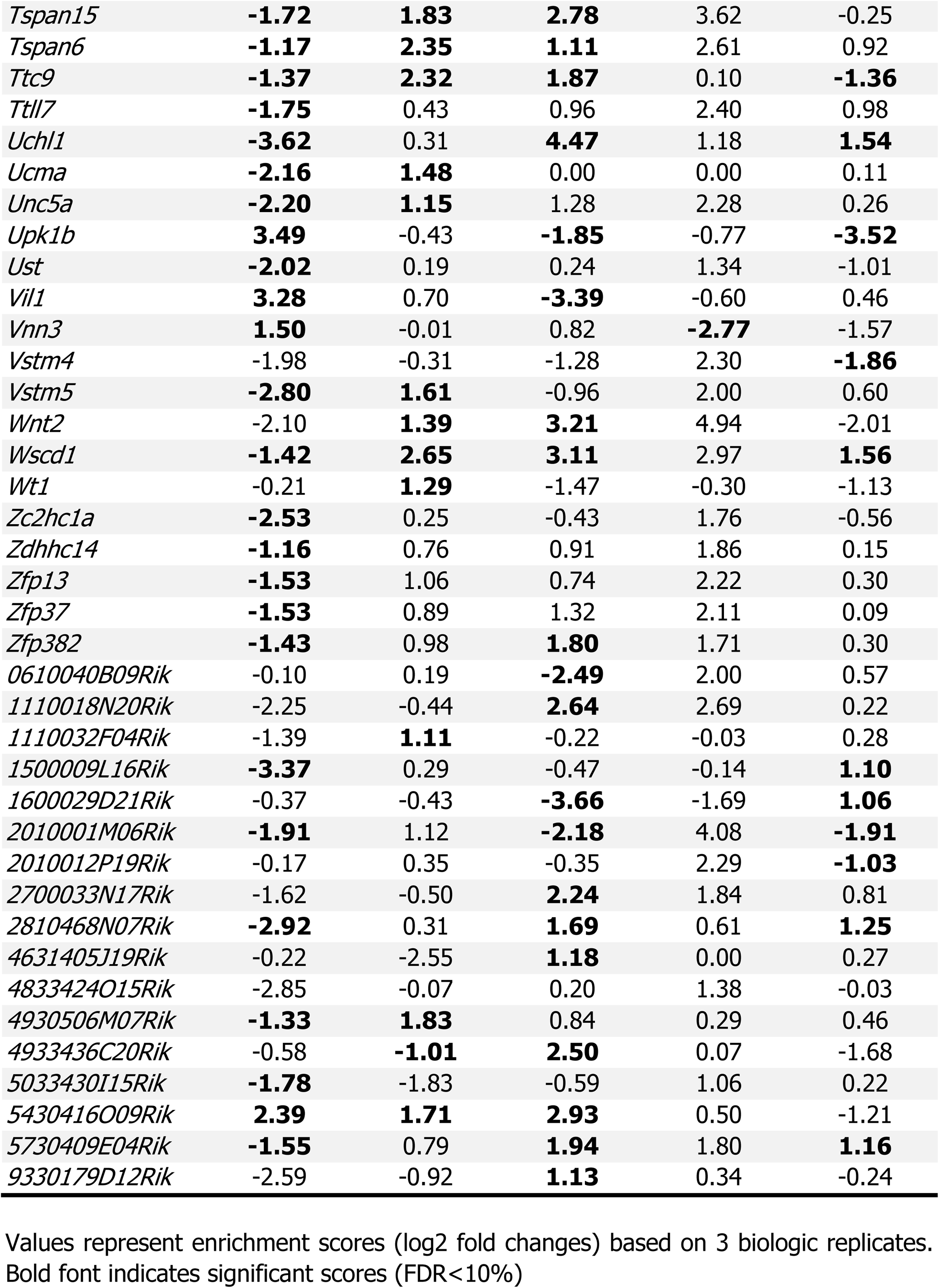
EC genes significantly enriched (log2 fold change >1) in at least 1 organ while also being significantly depleted (log2 fold change <-1) in at least 1 other organ.

**Supplemental Table S3:**
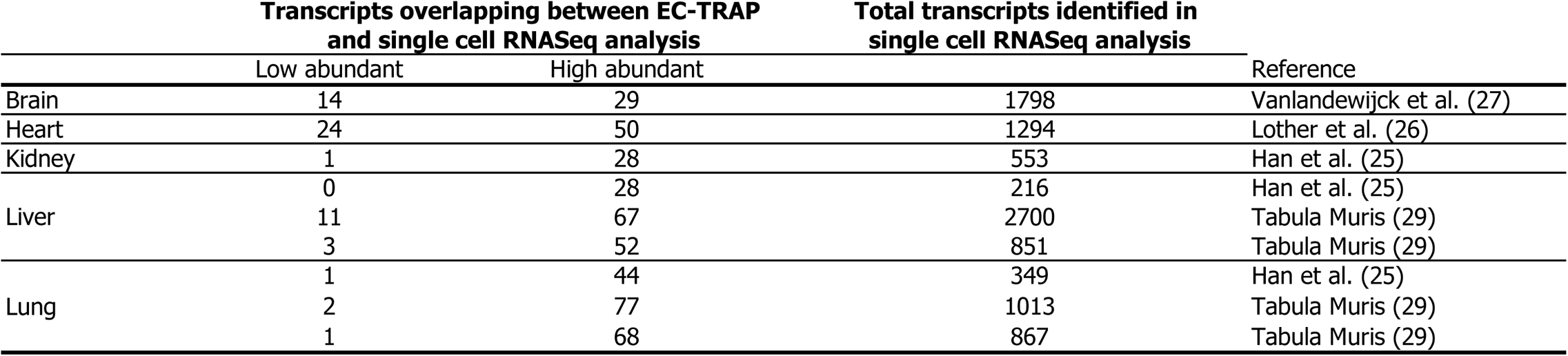
Comparison of high and low abundant EC-enriched genes identified by TRAP versus EC genes identified by single cell data RNASeq.

**Supplemental Table S4:**
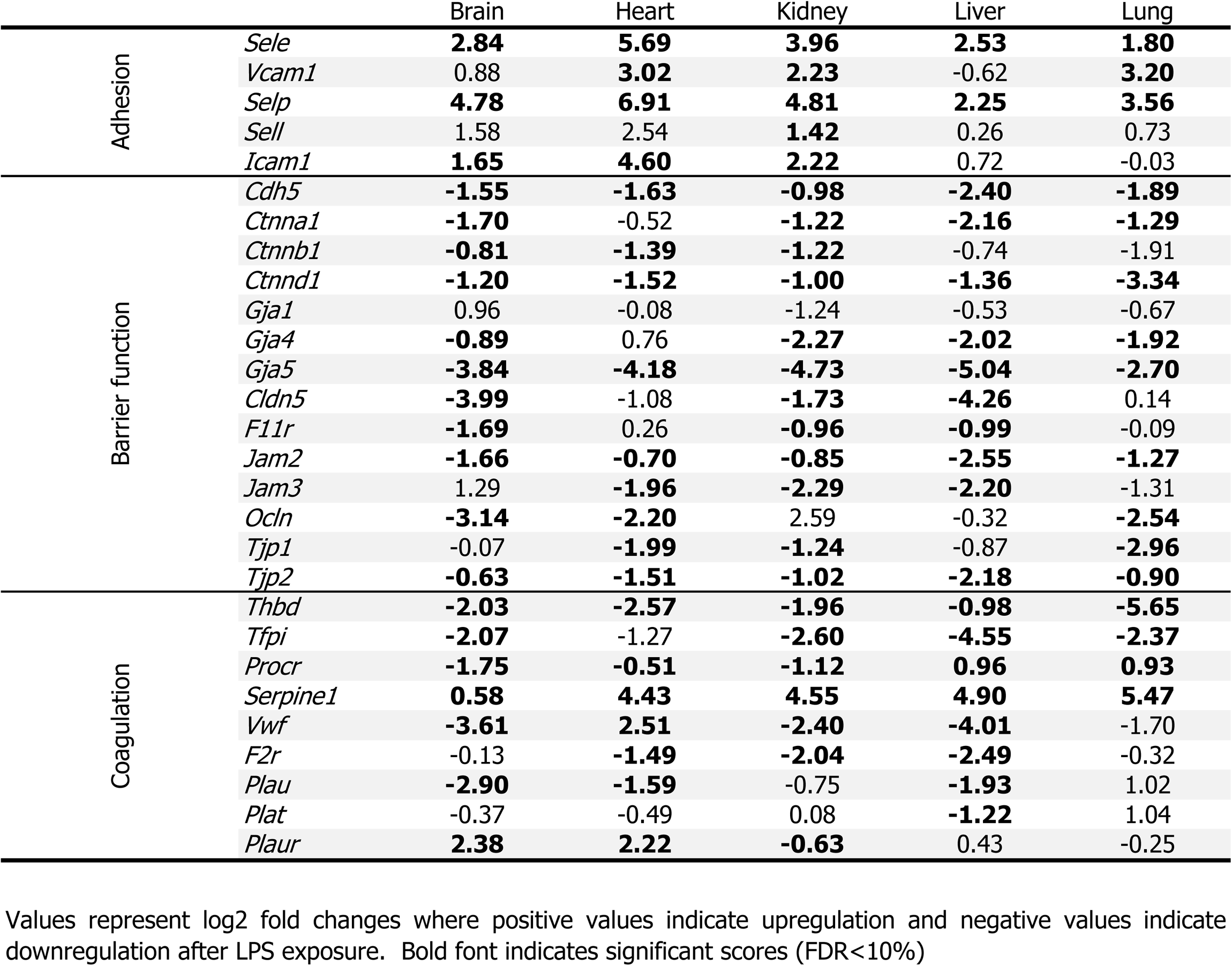
Selected genes previously identified to be affected in endotoxemia.

**Supplemental Table S5:**
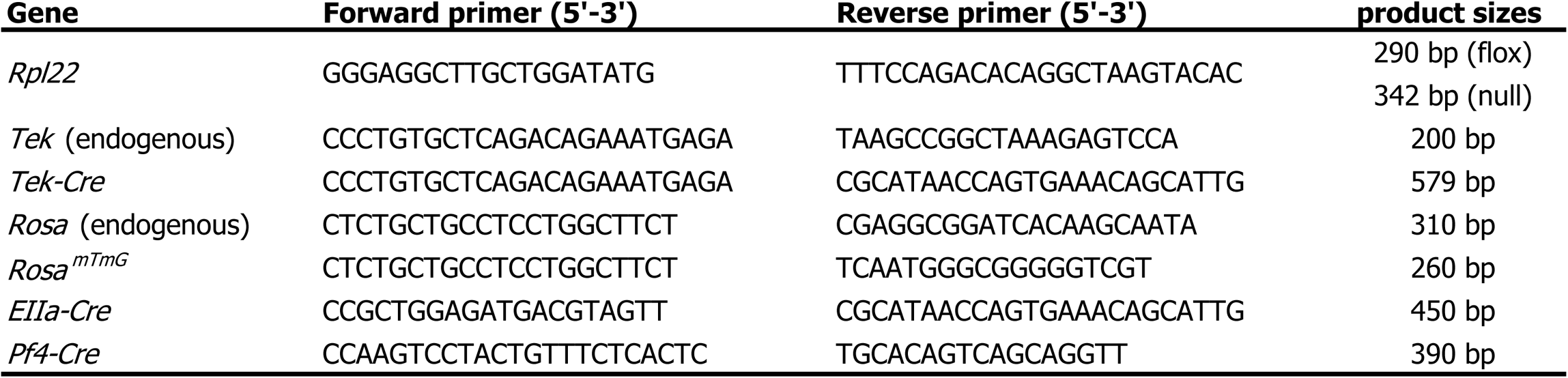
Genotyping primers and expected amplicon sizes.

**Supplemental Table S6:**
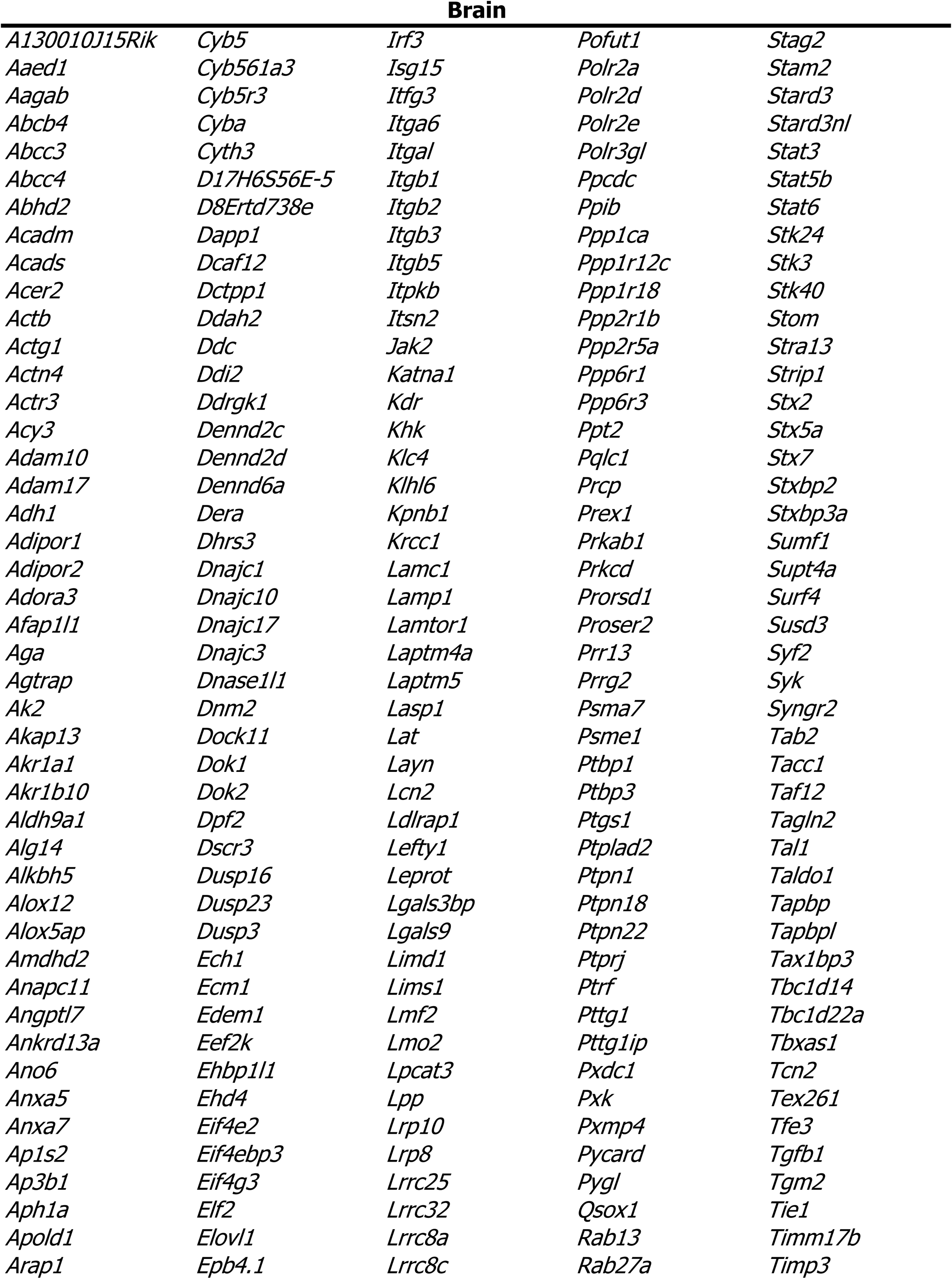
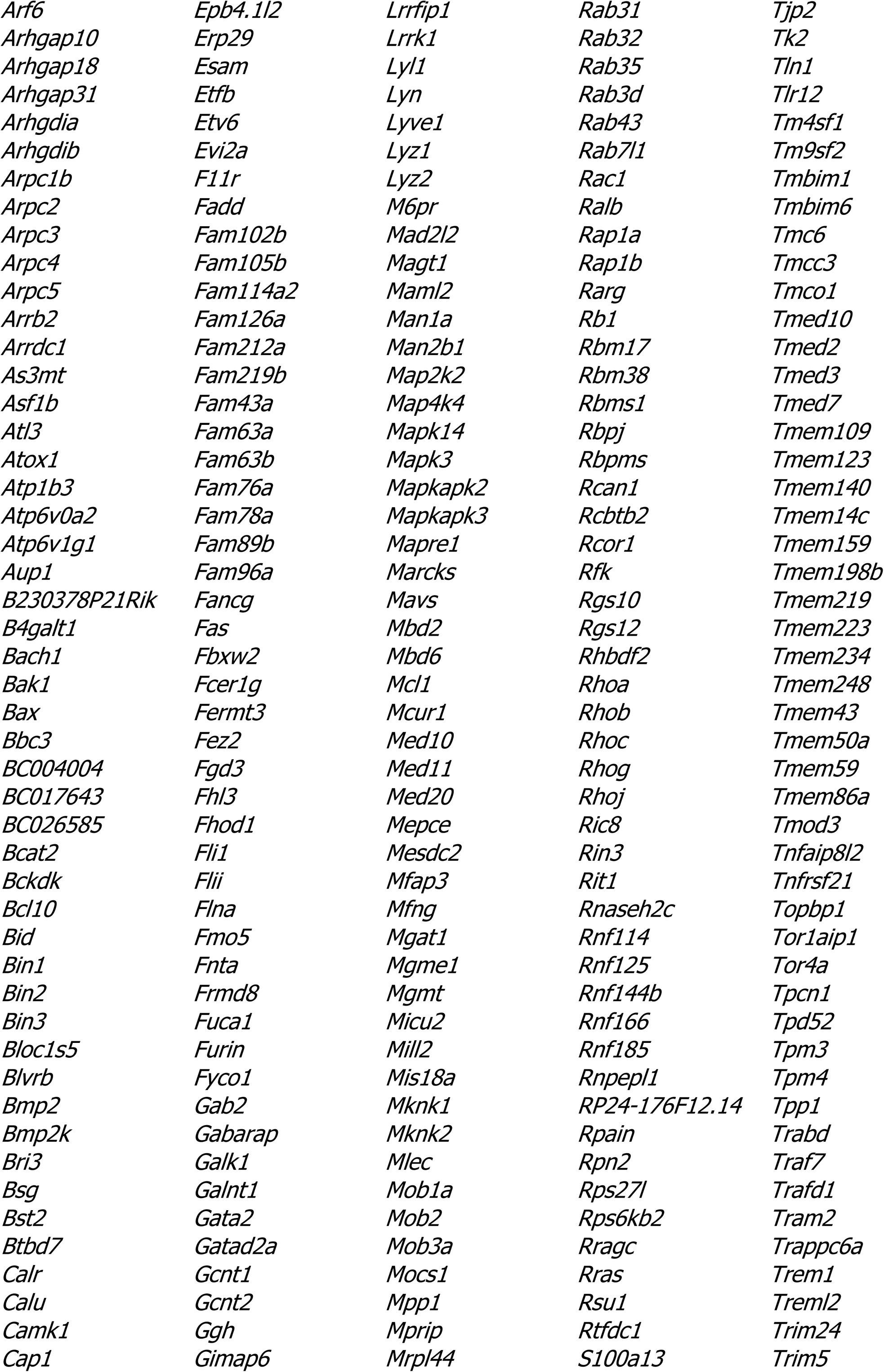
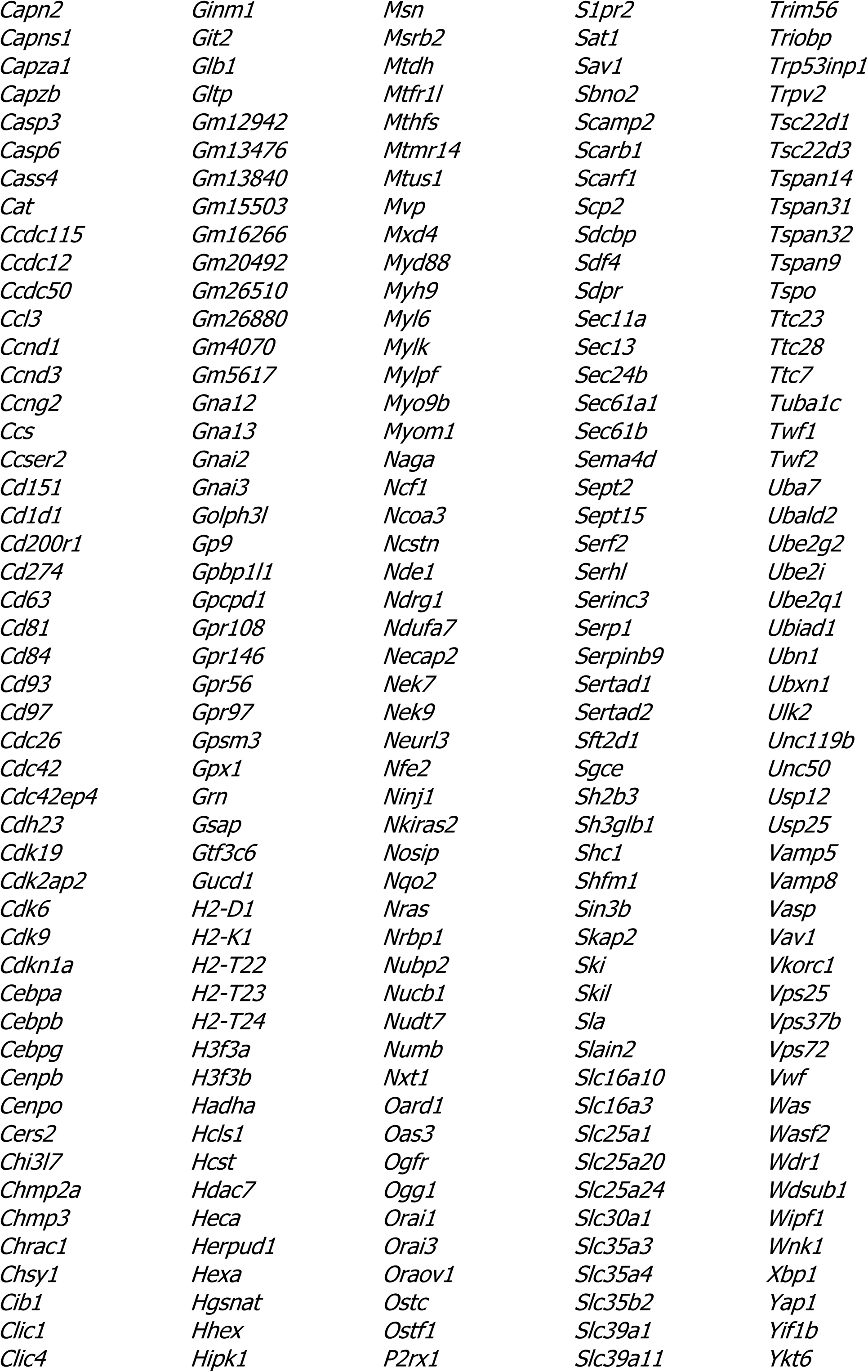
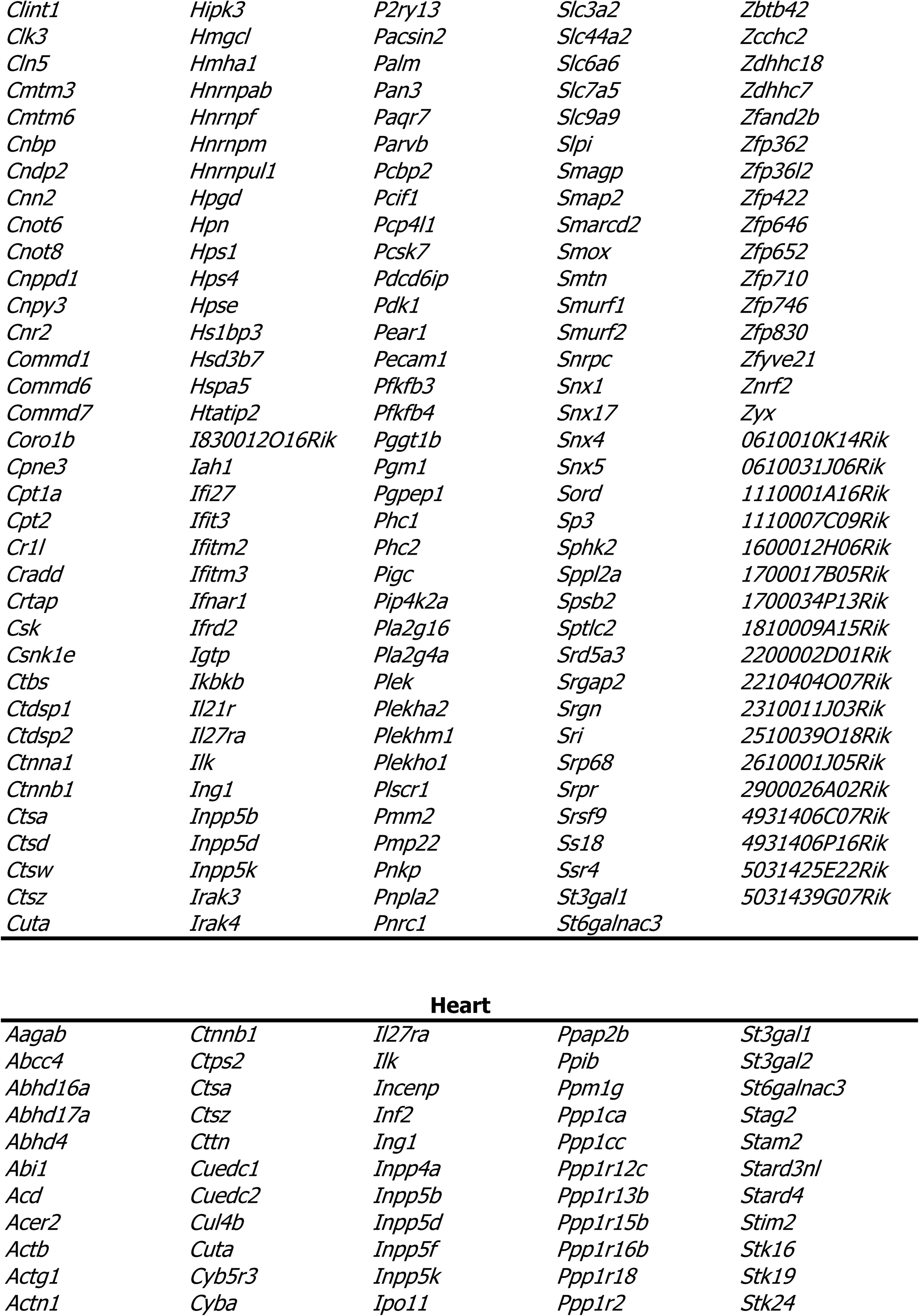
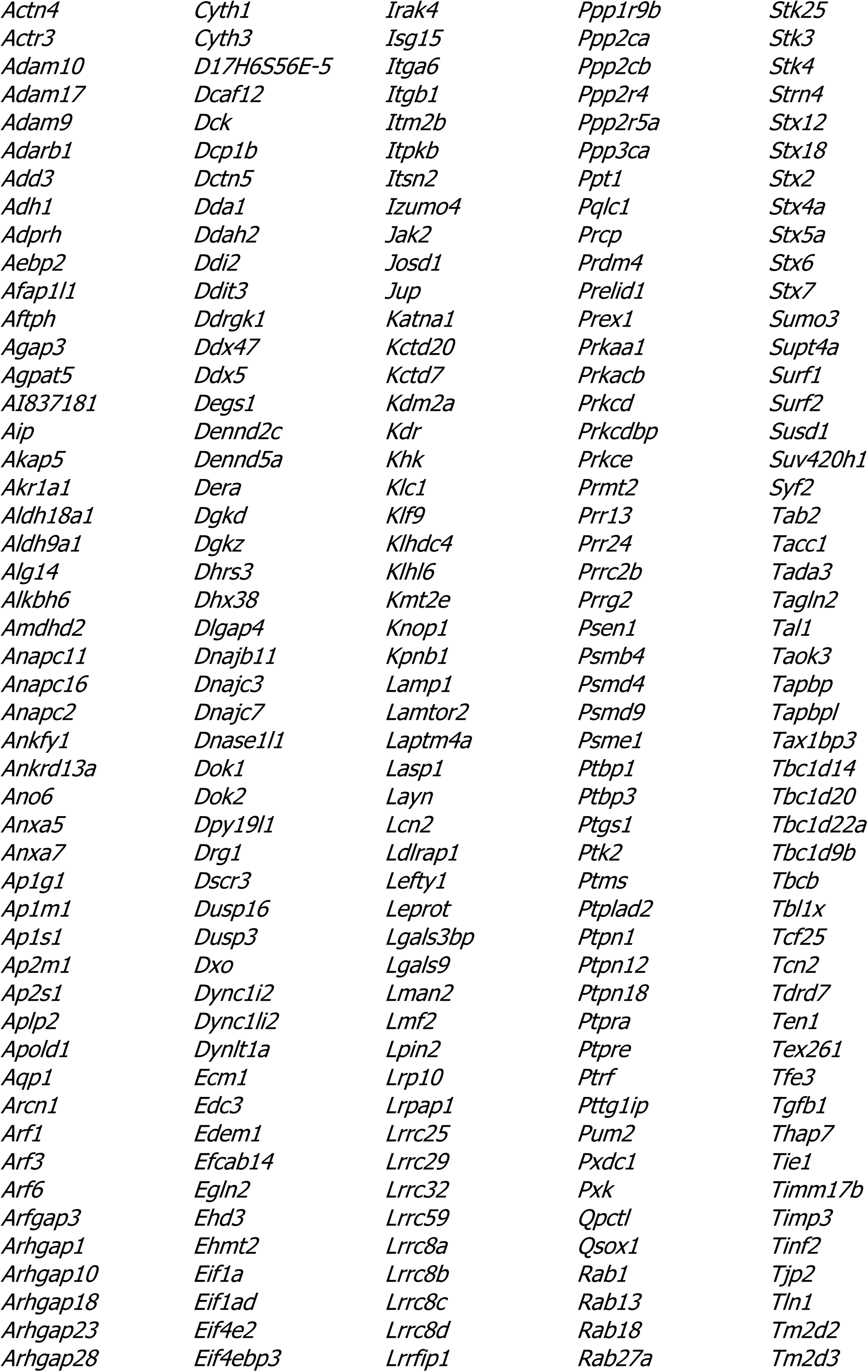
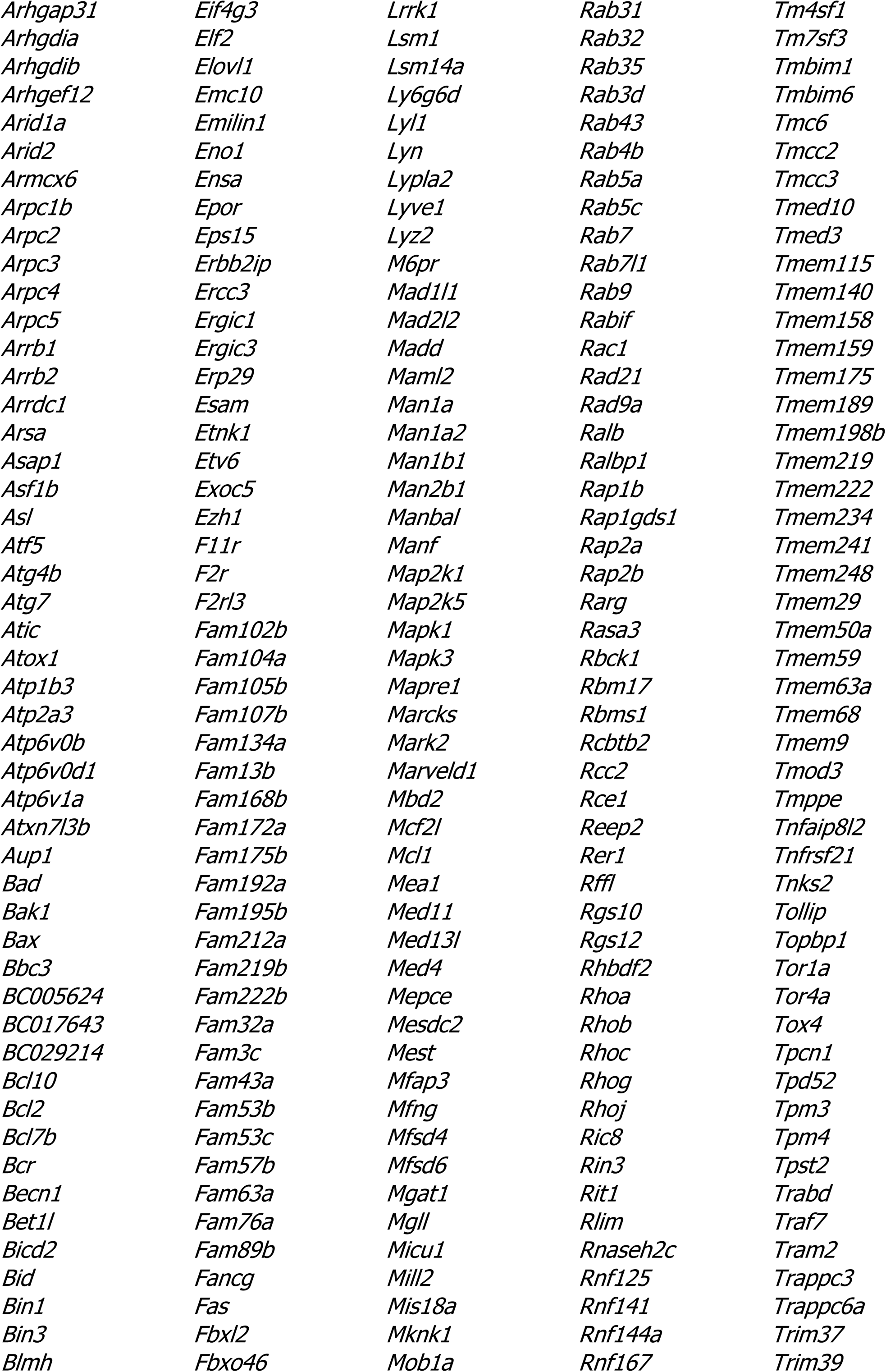
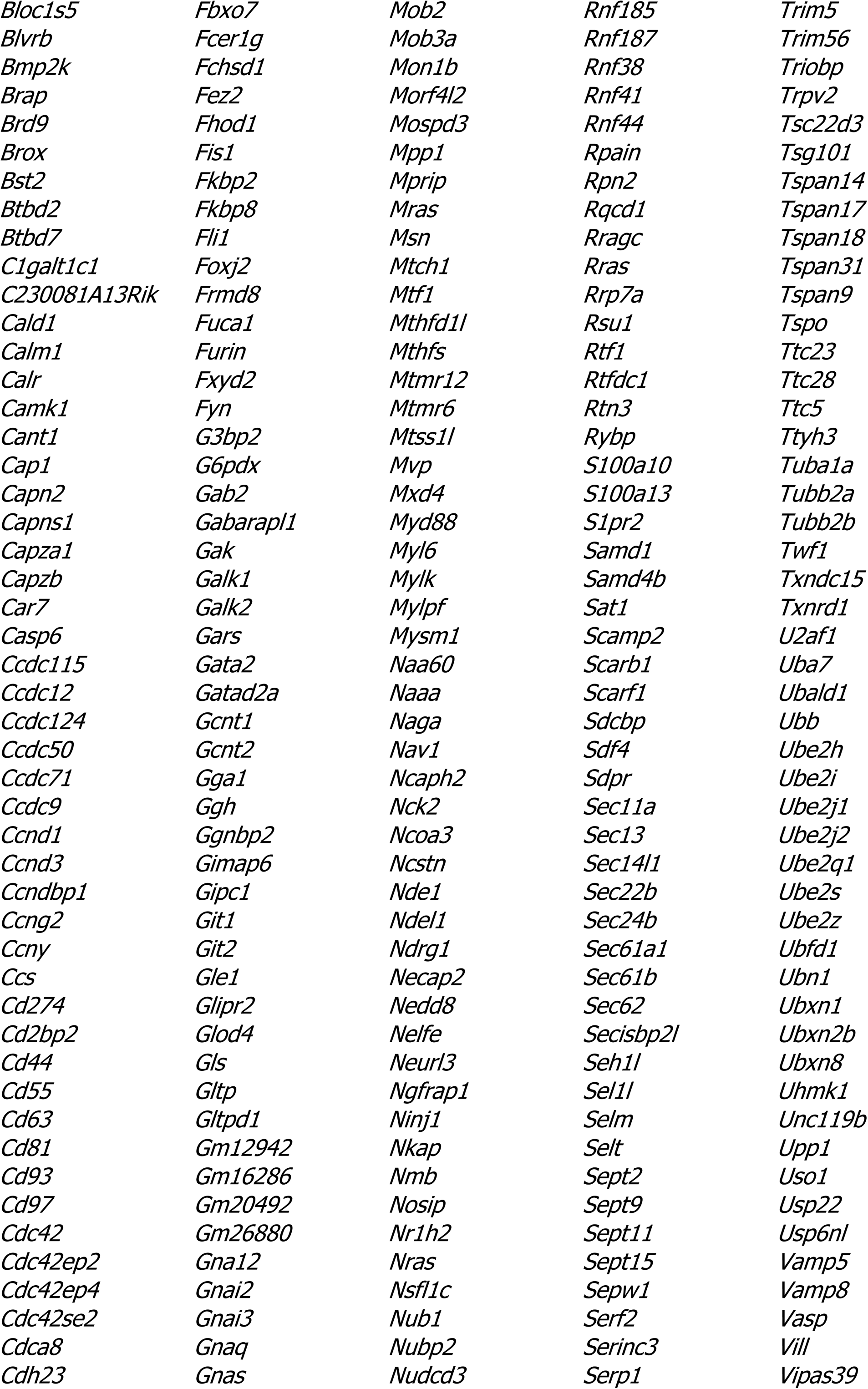
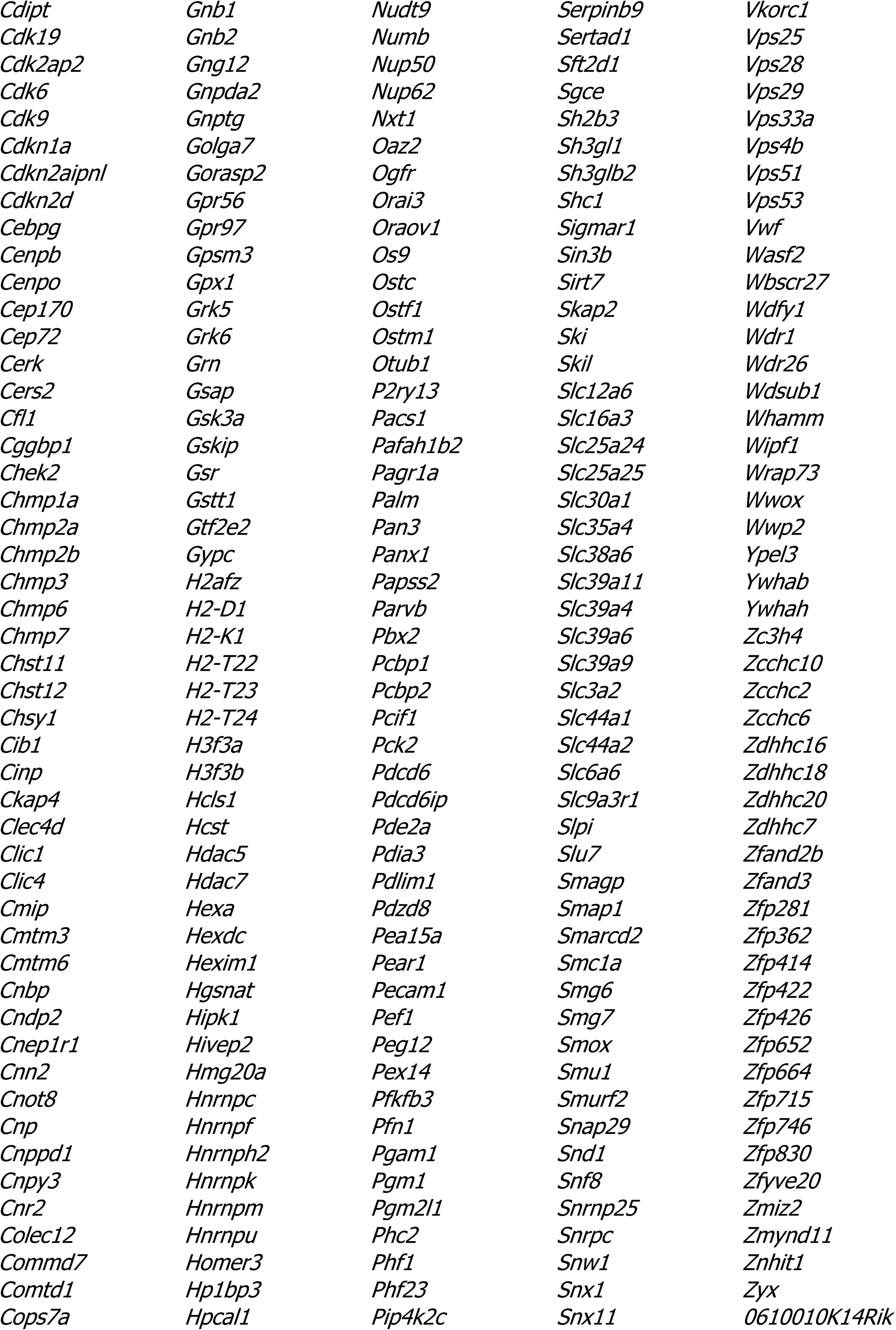
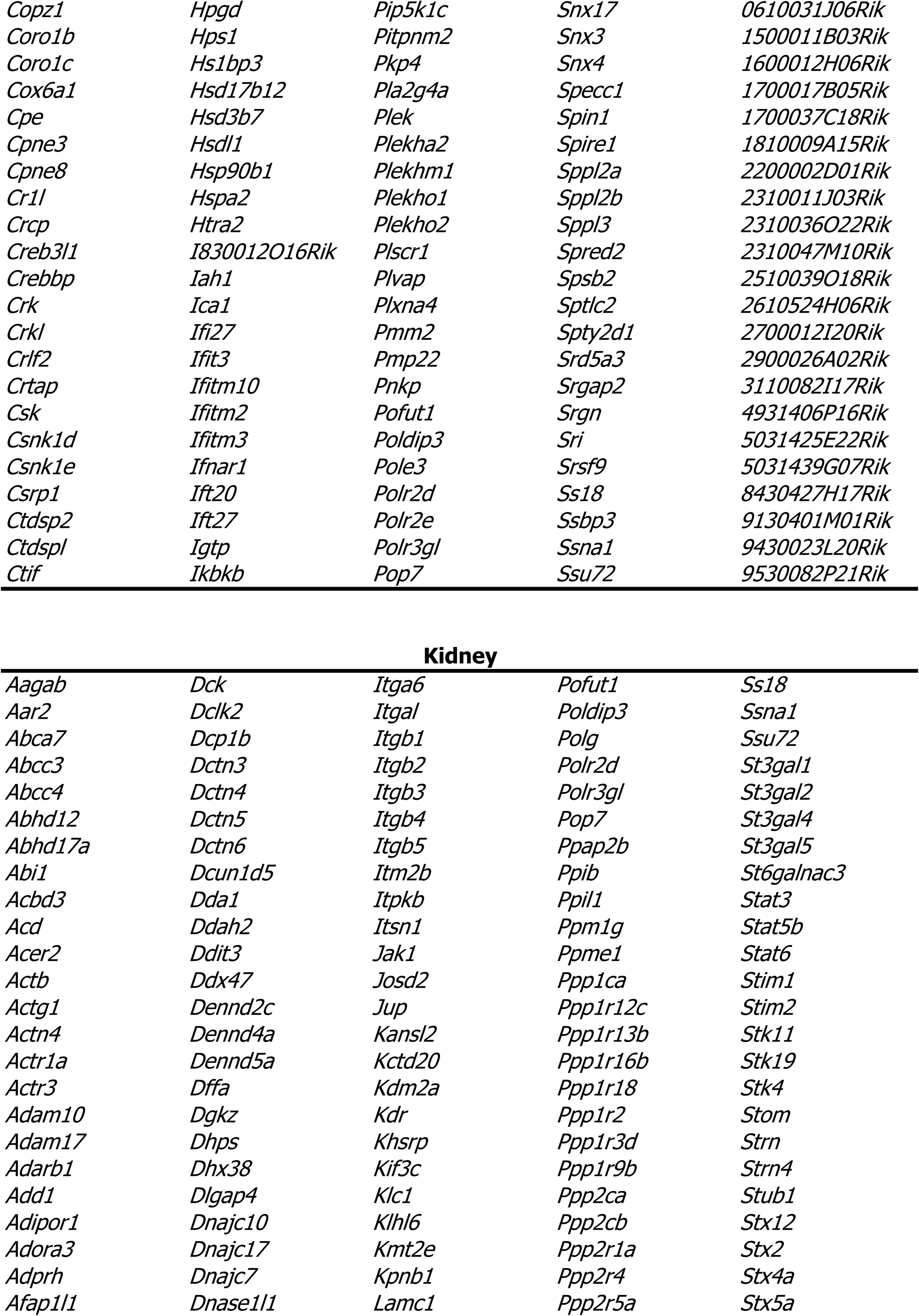
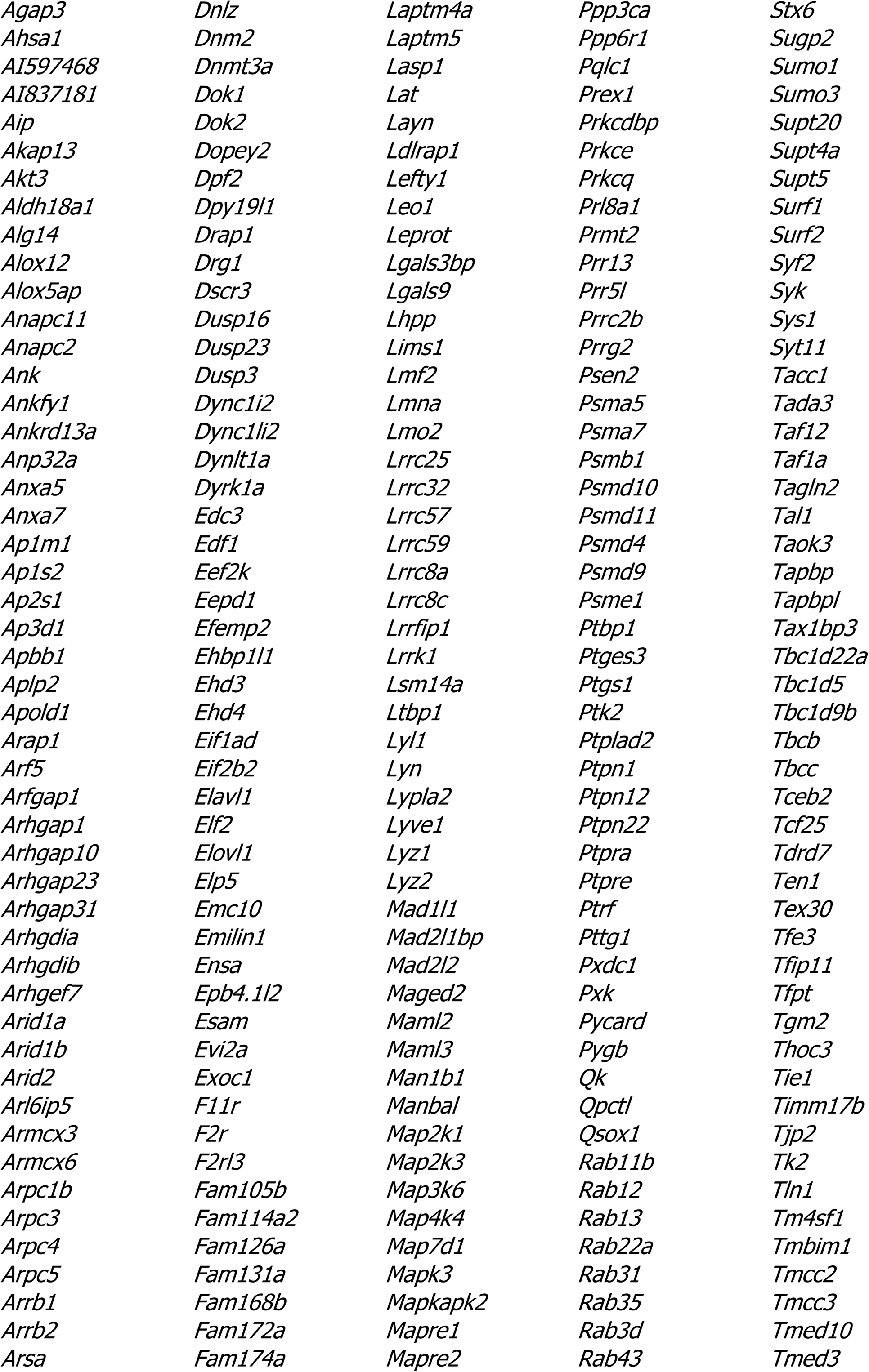
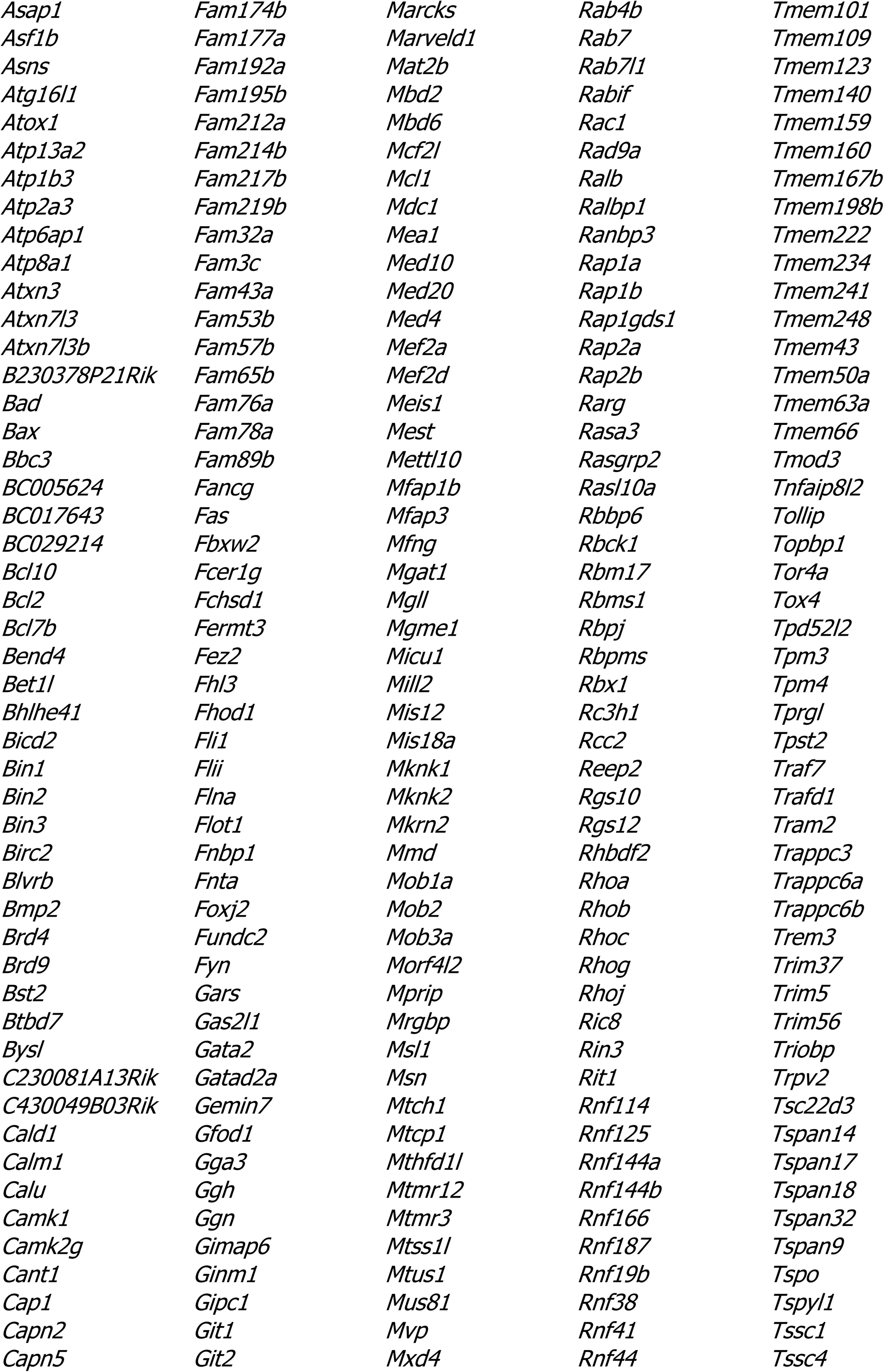
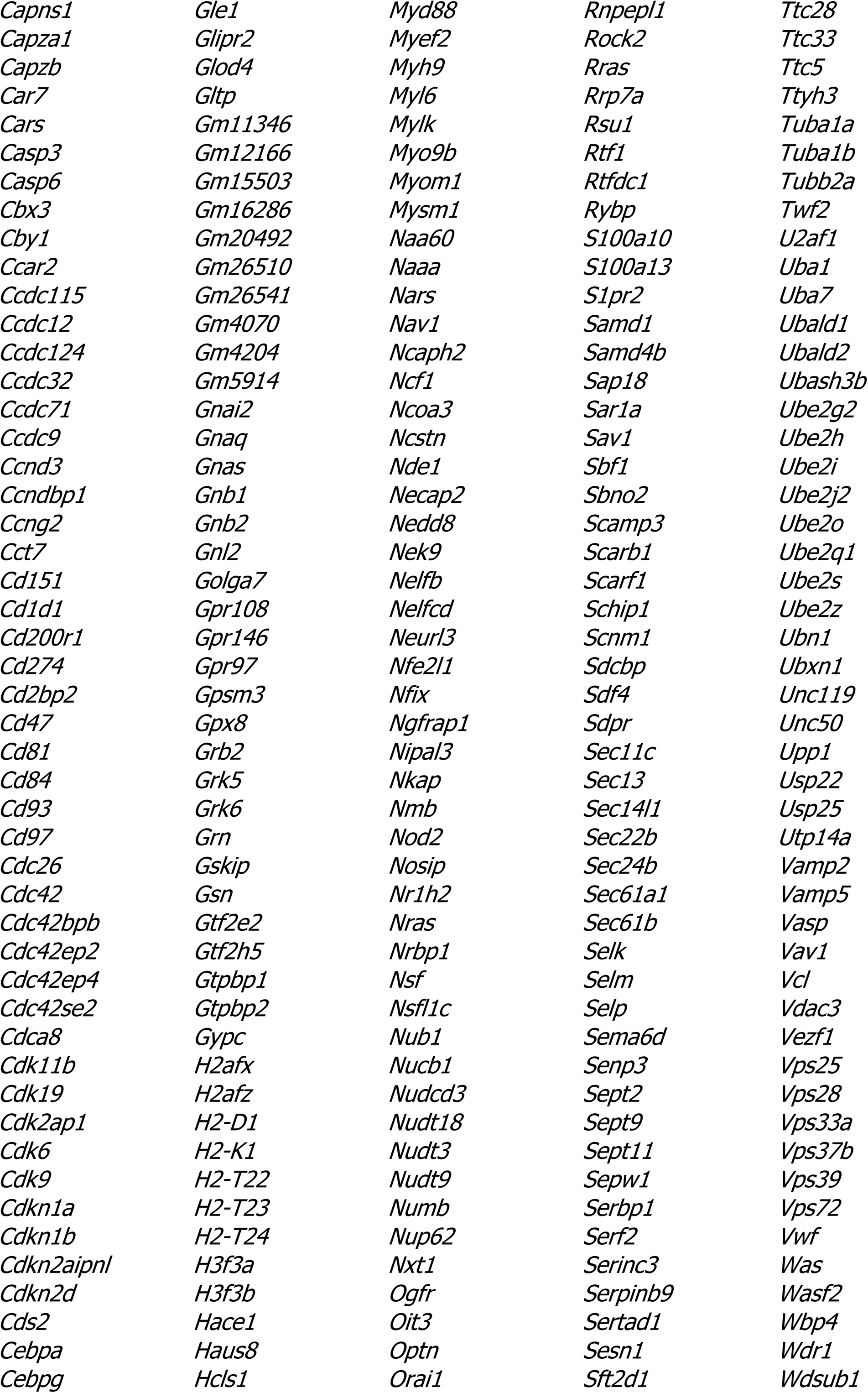
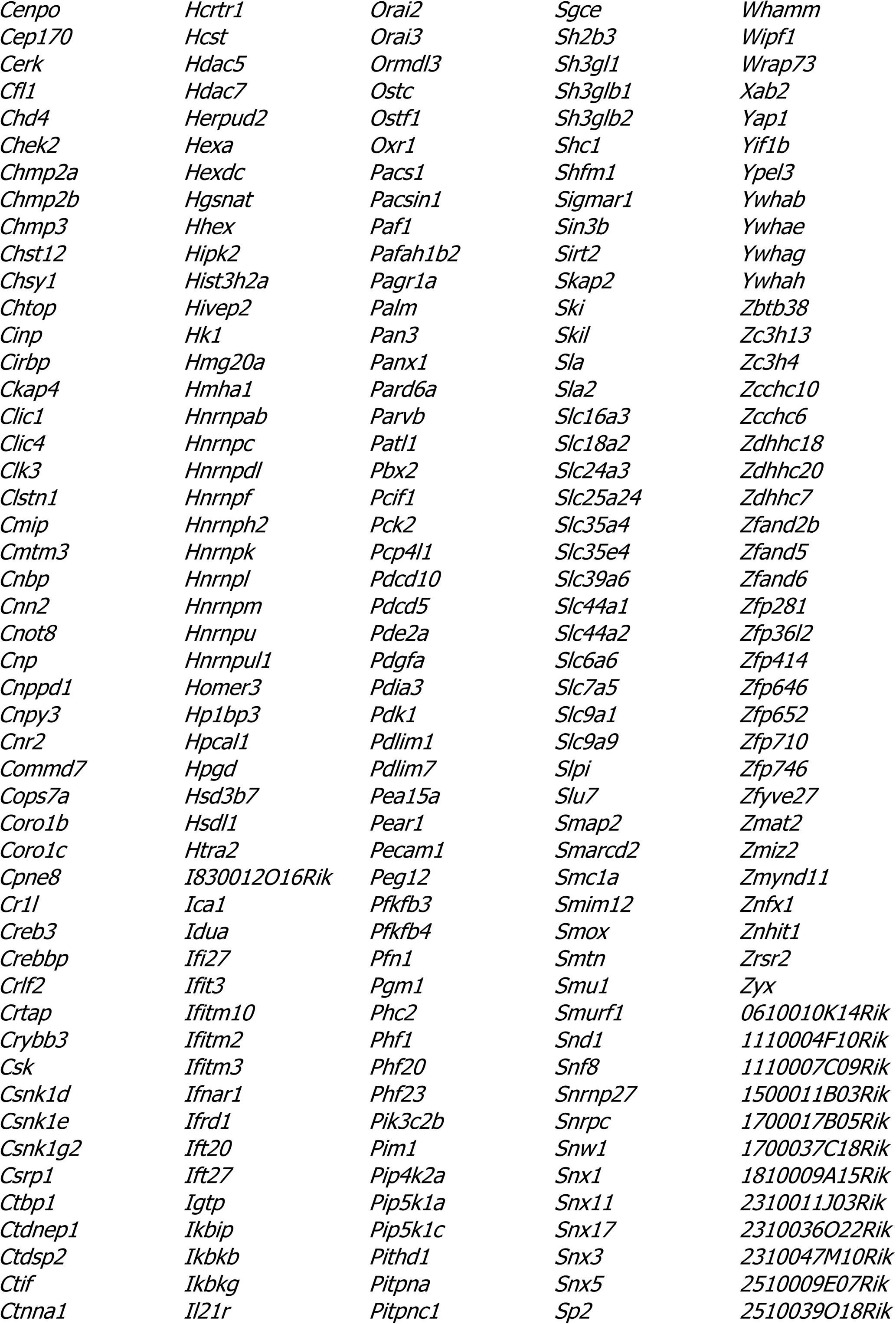
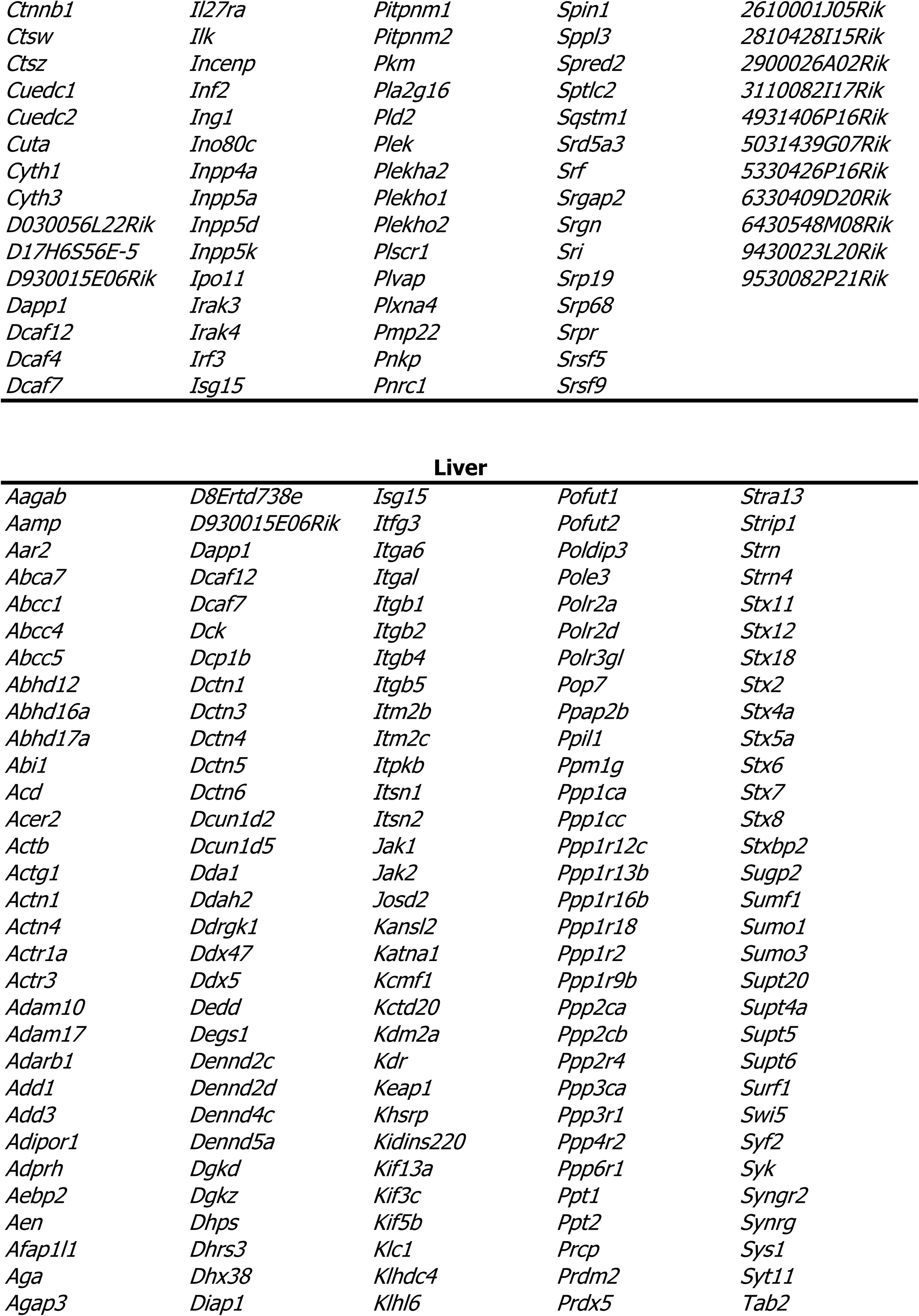
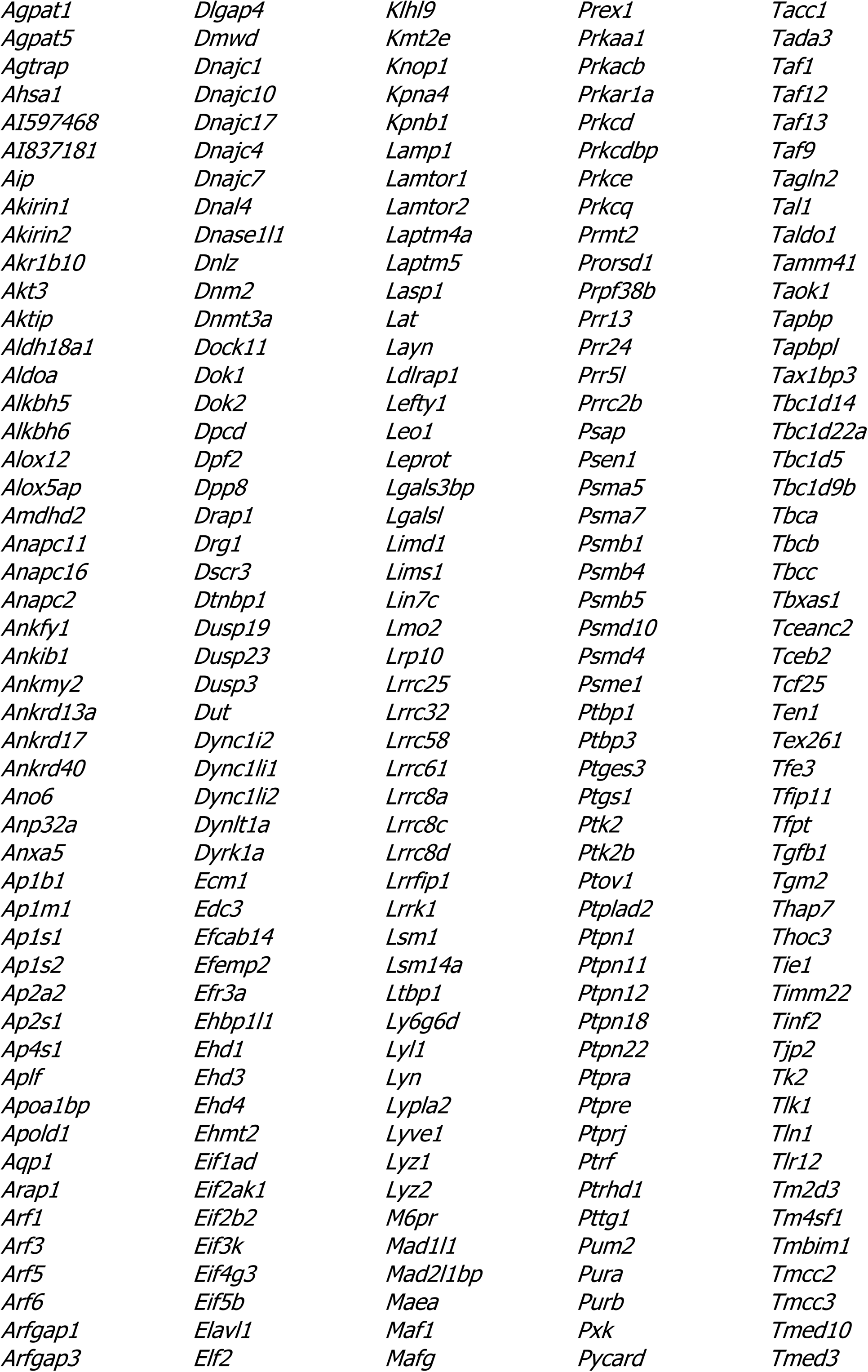
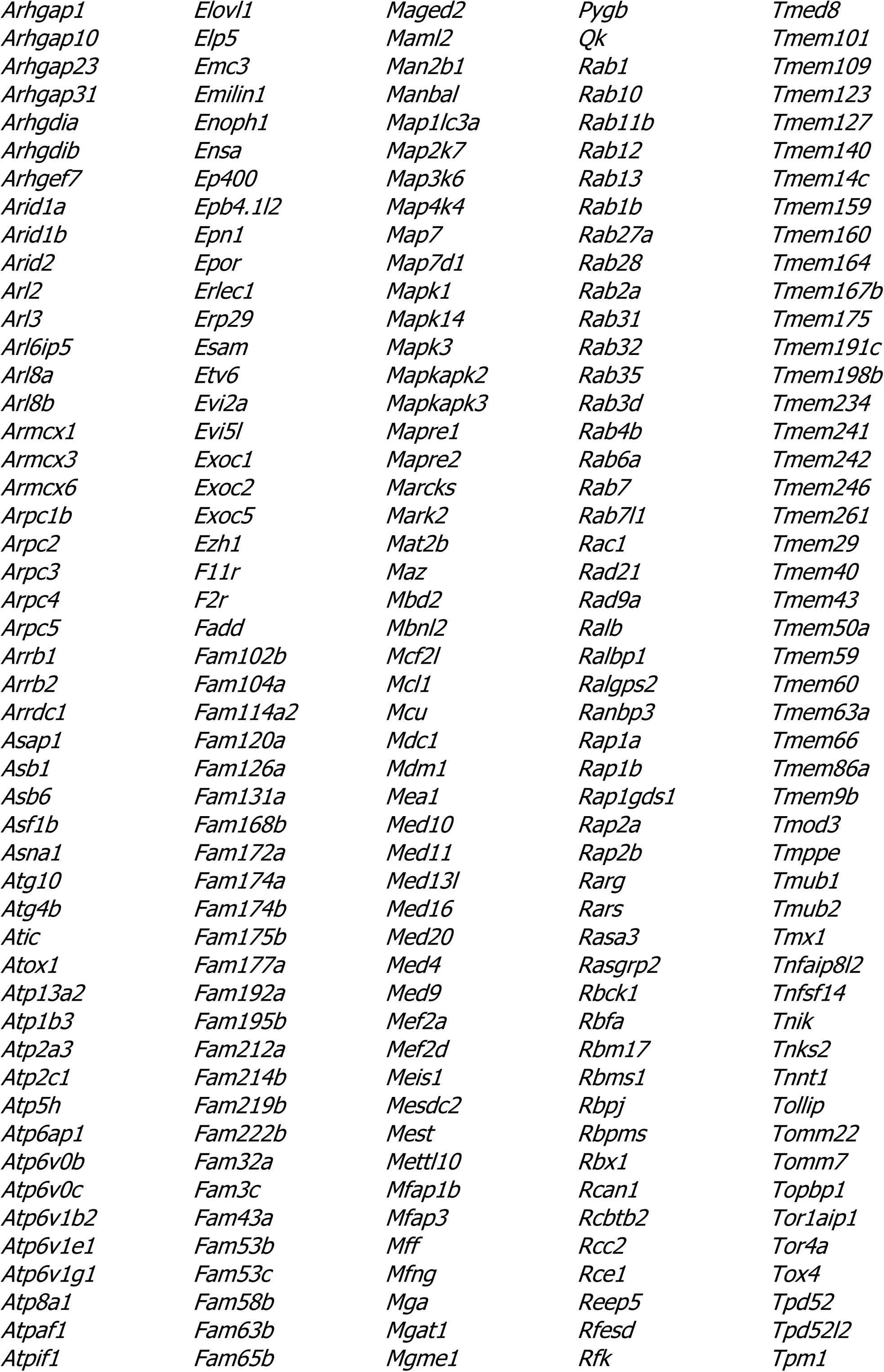
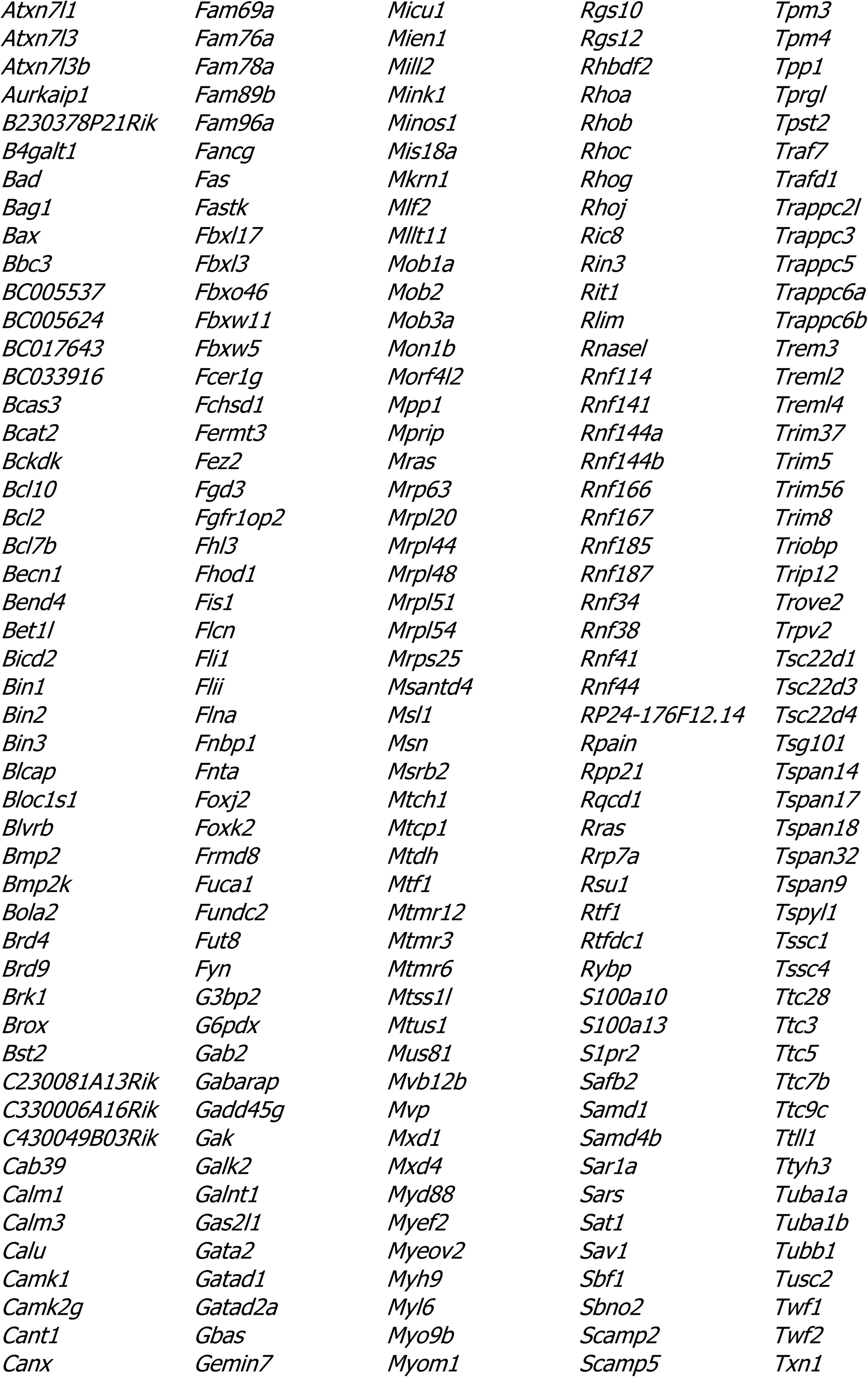
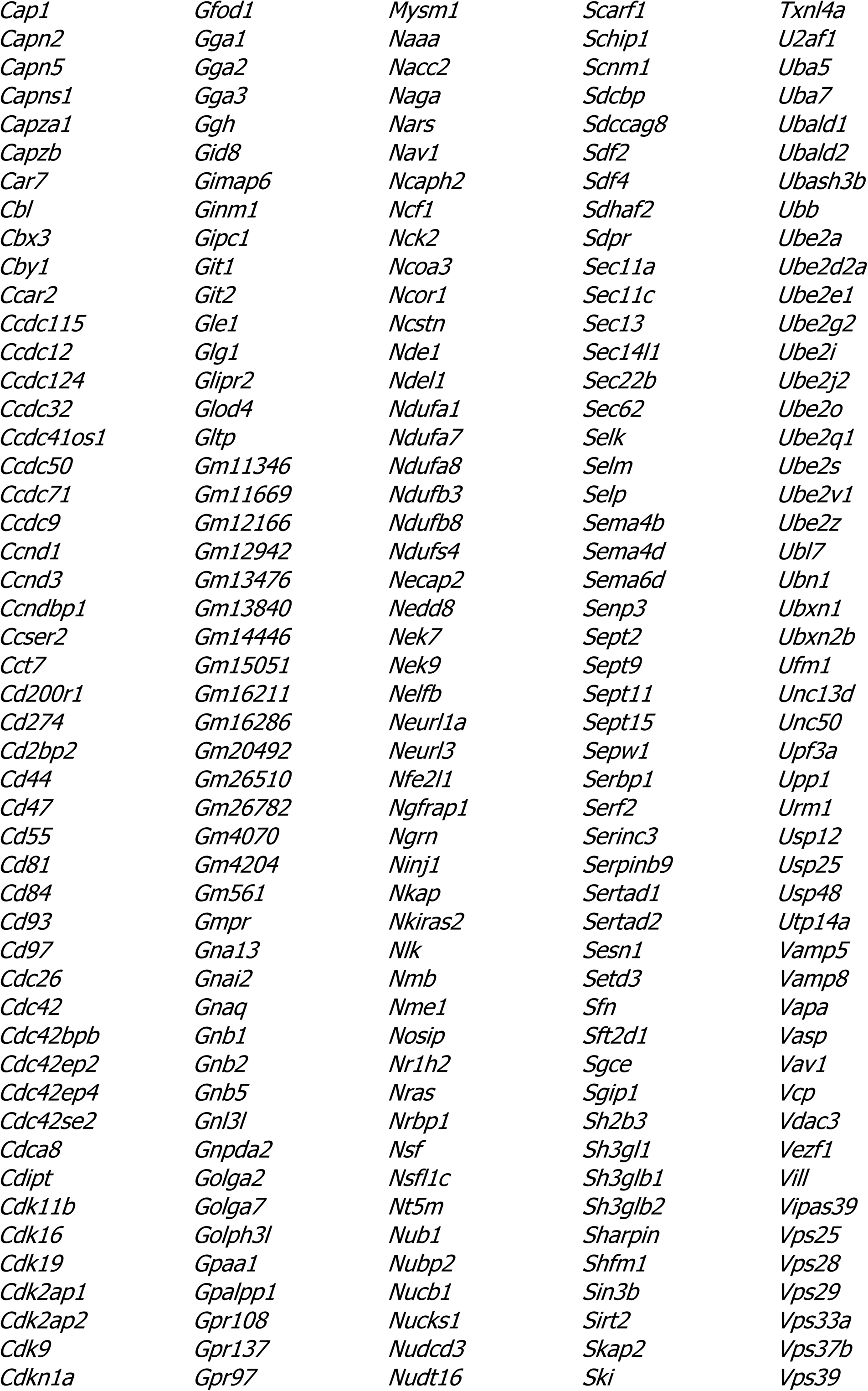
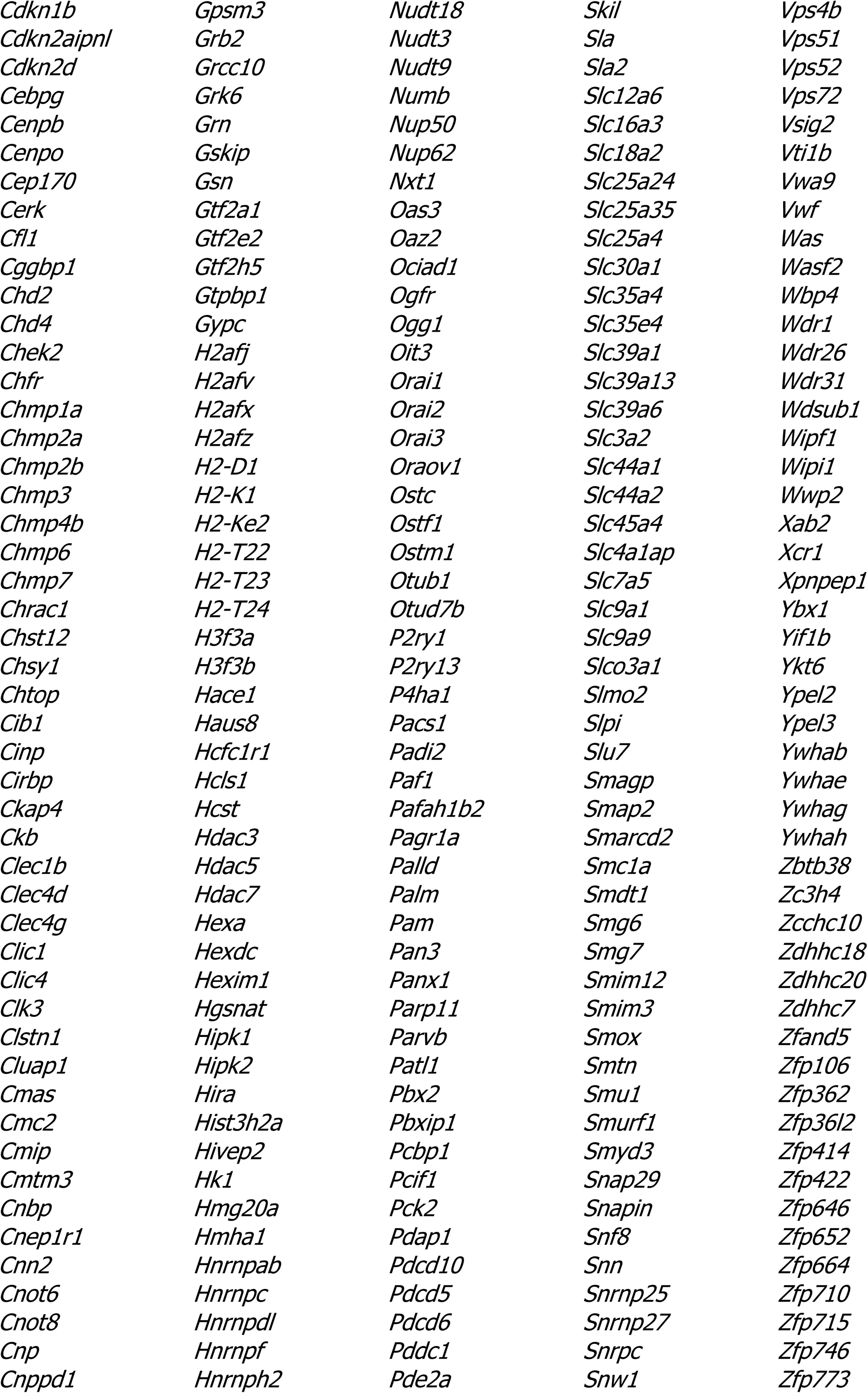
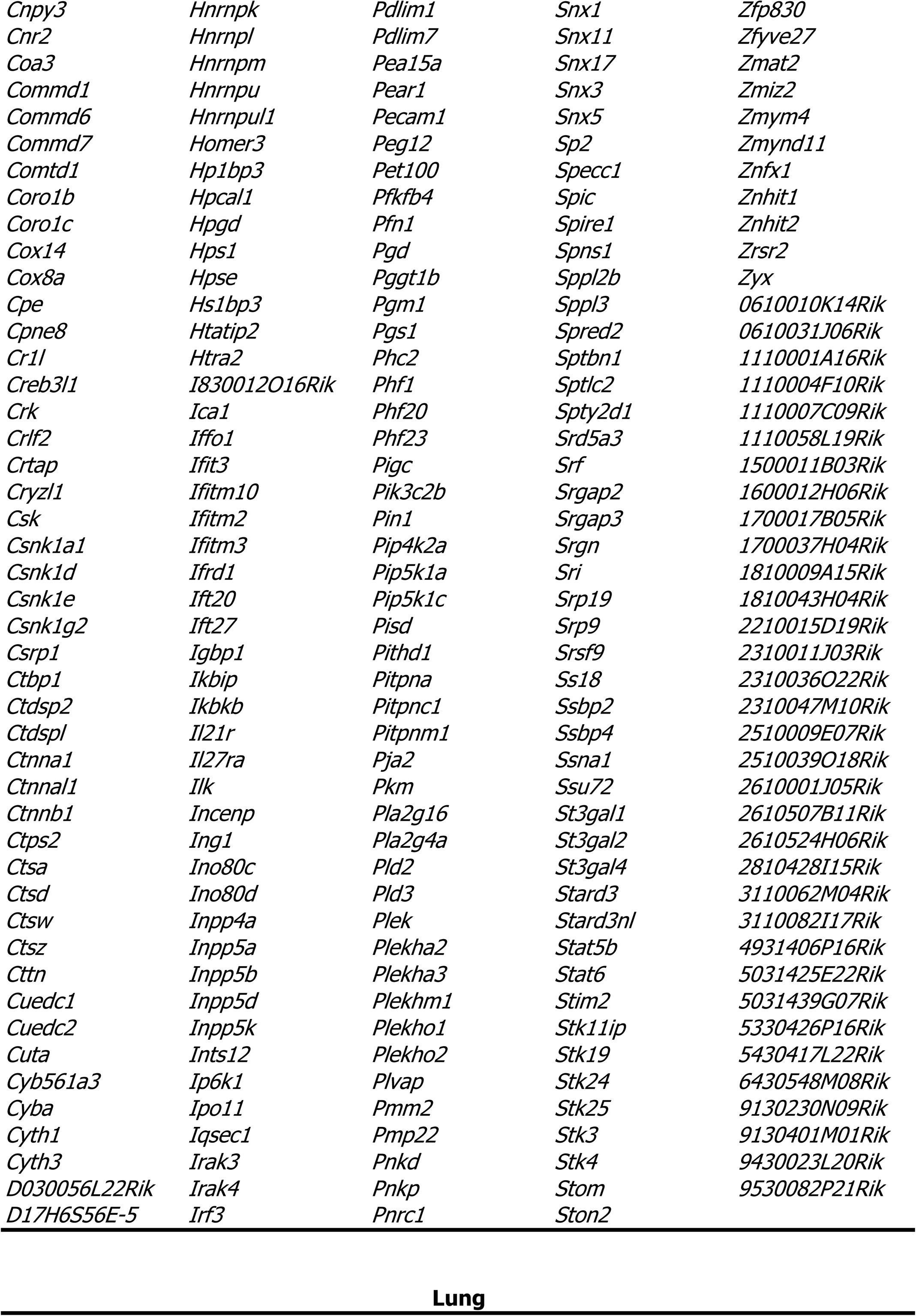
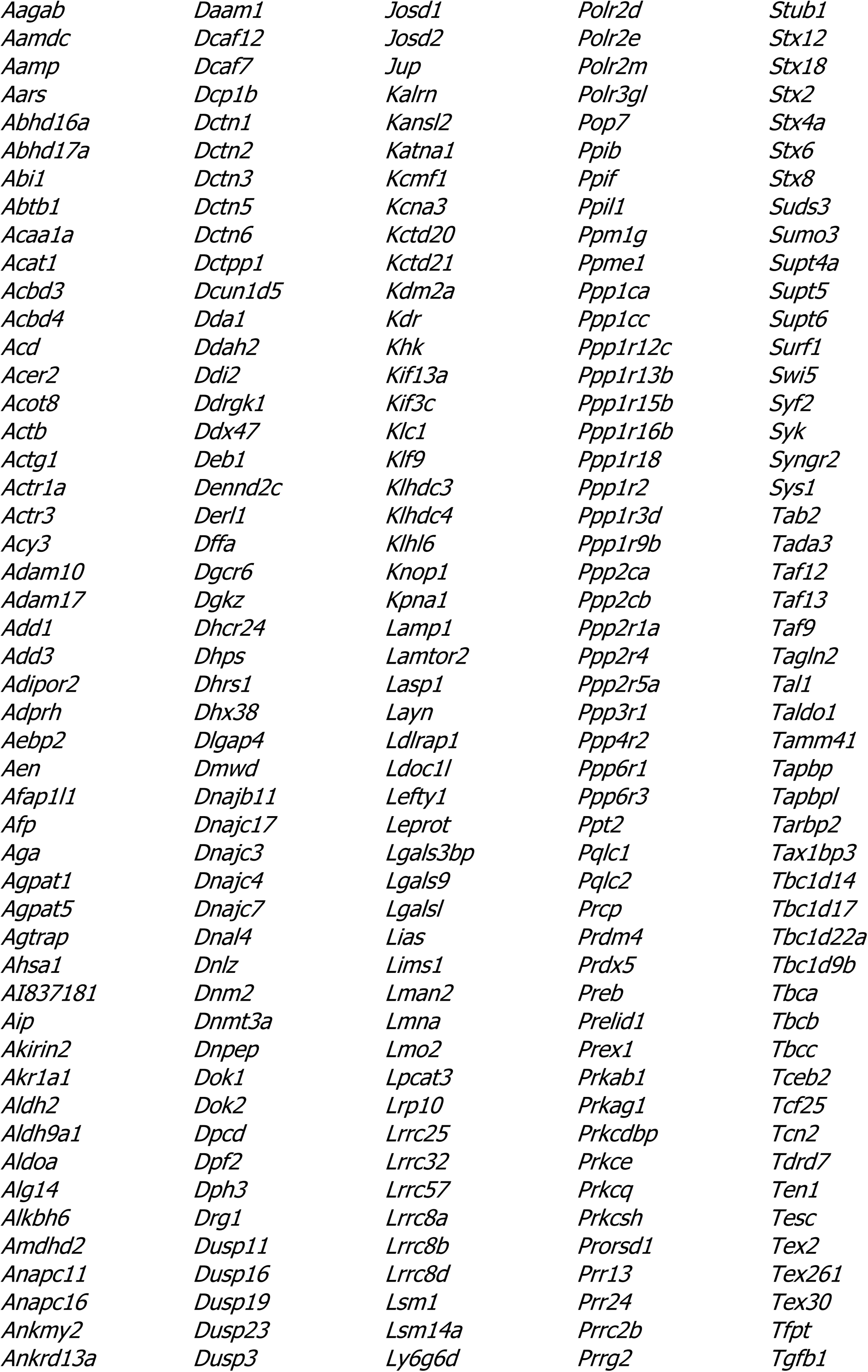
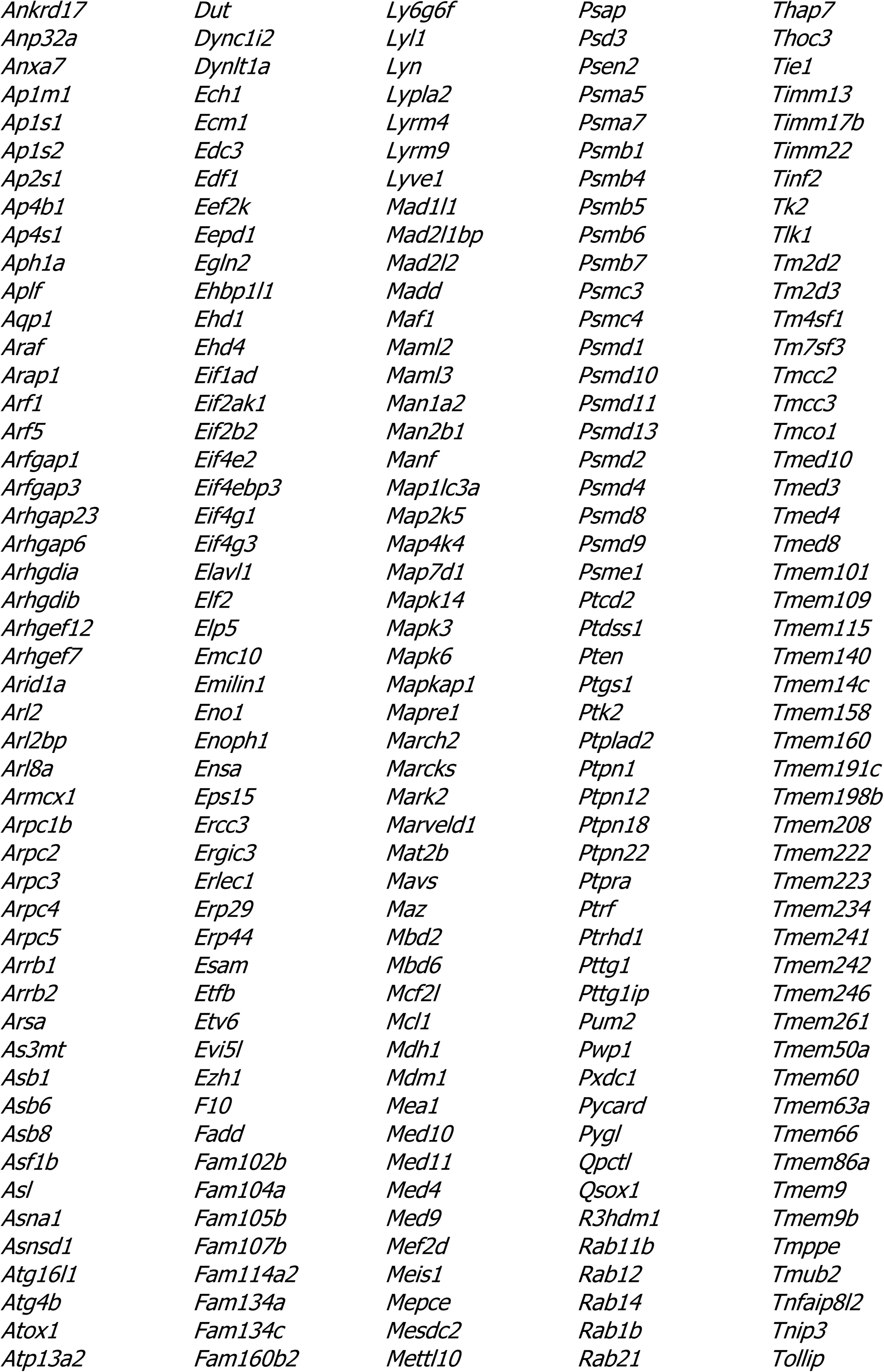
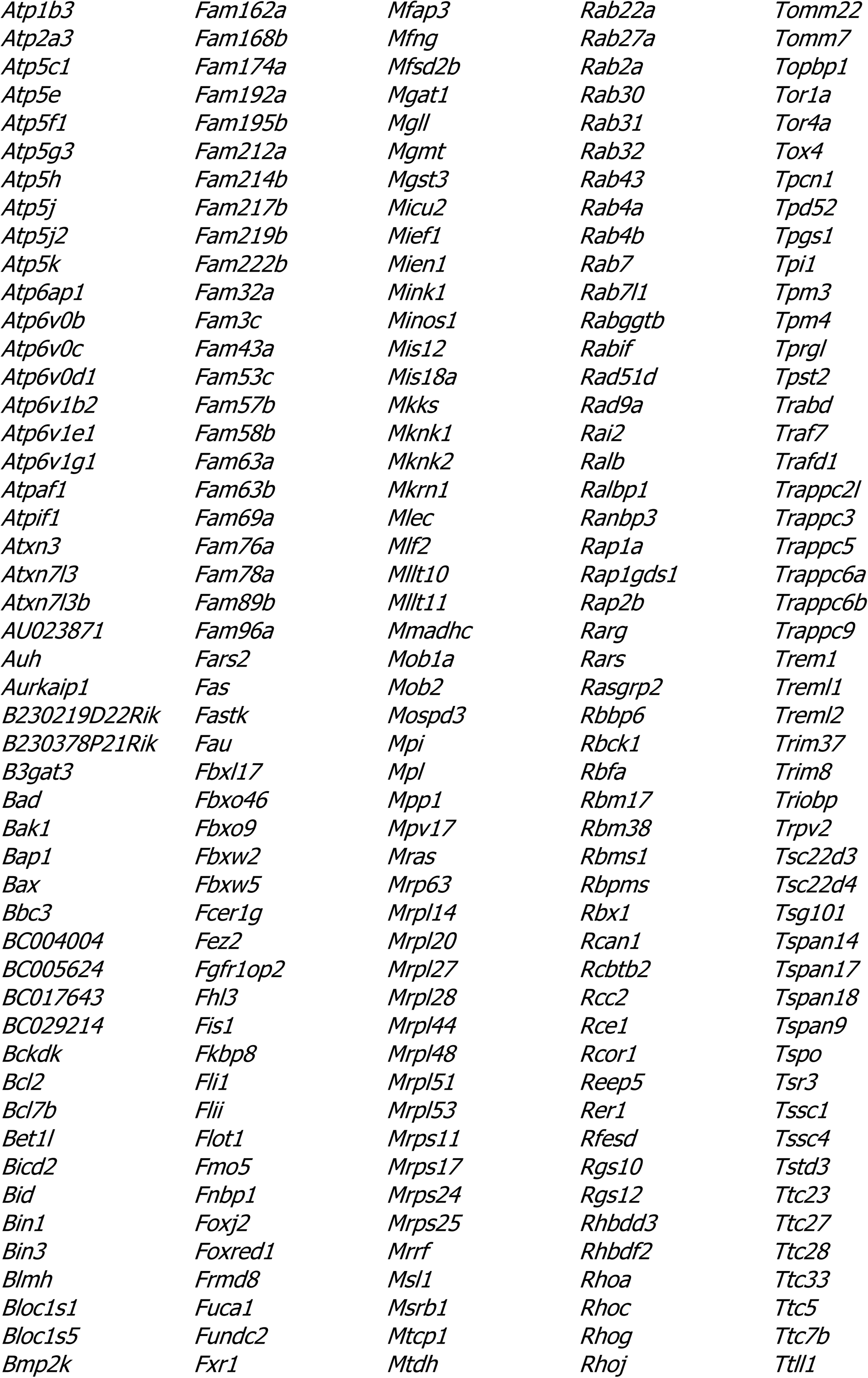
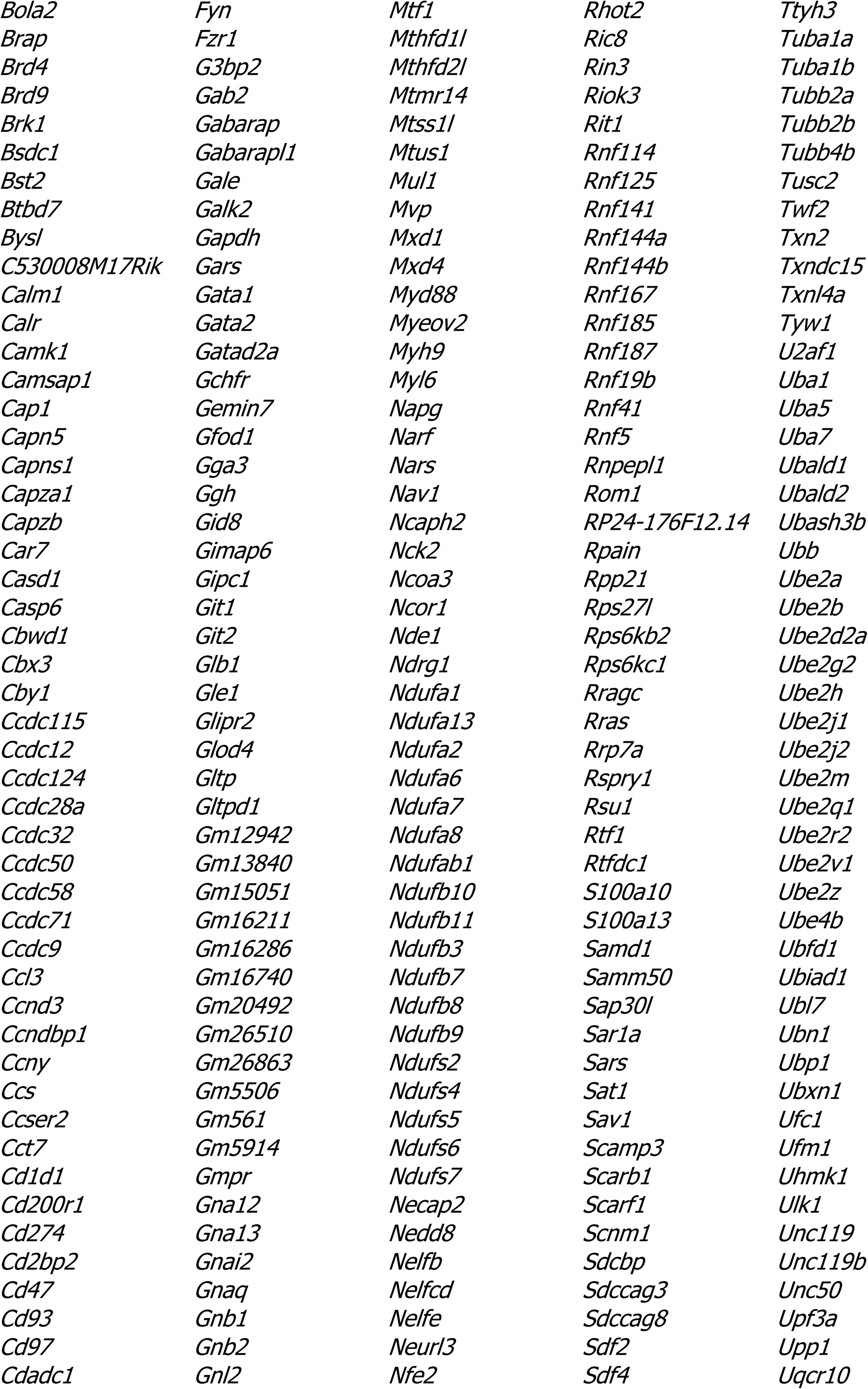
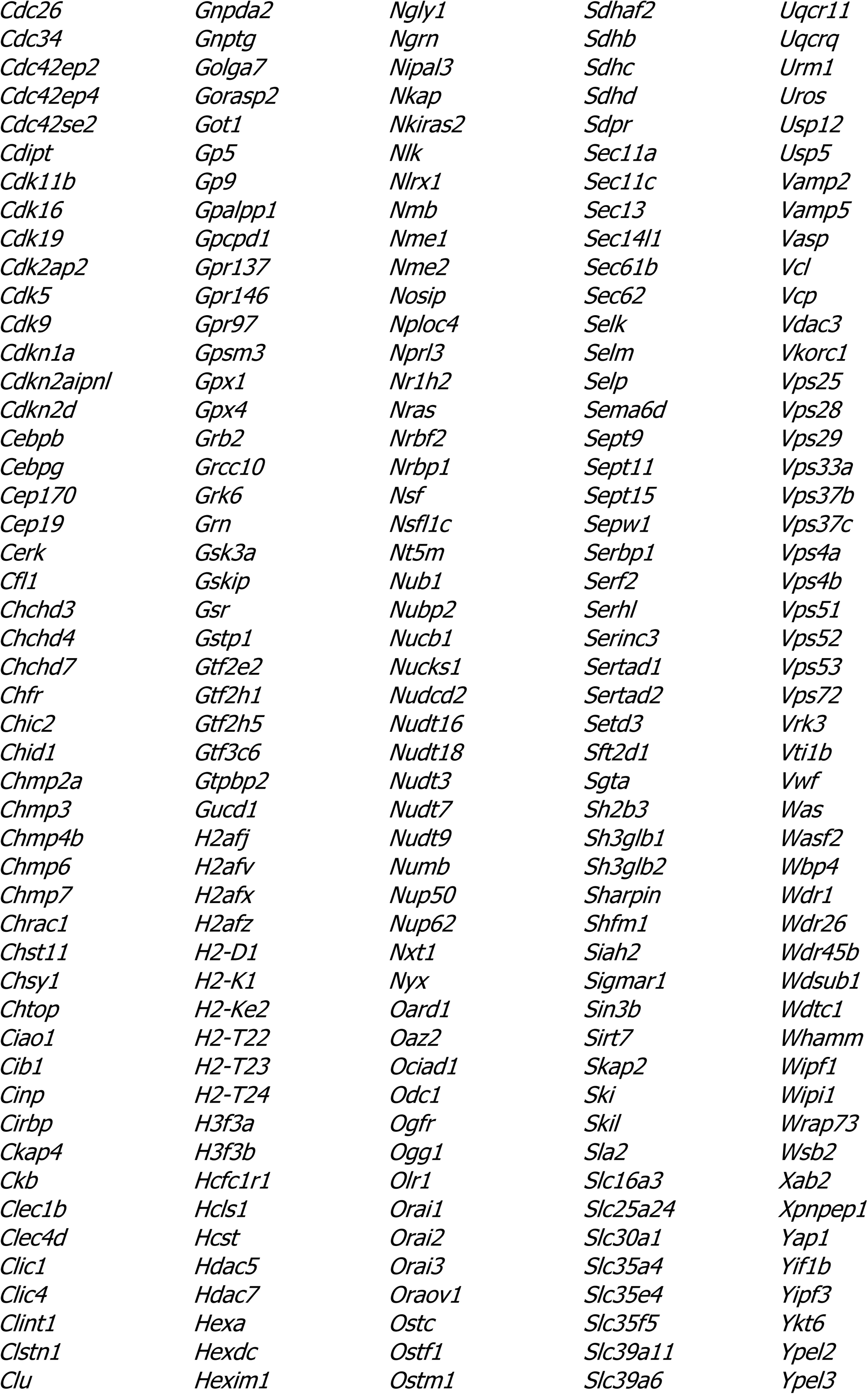
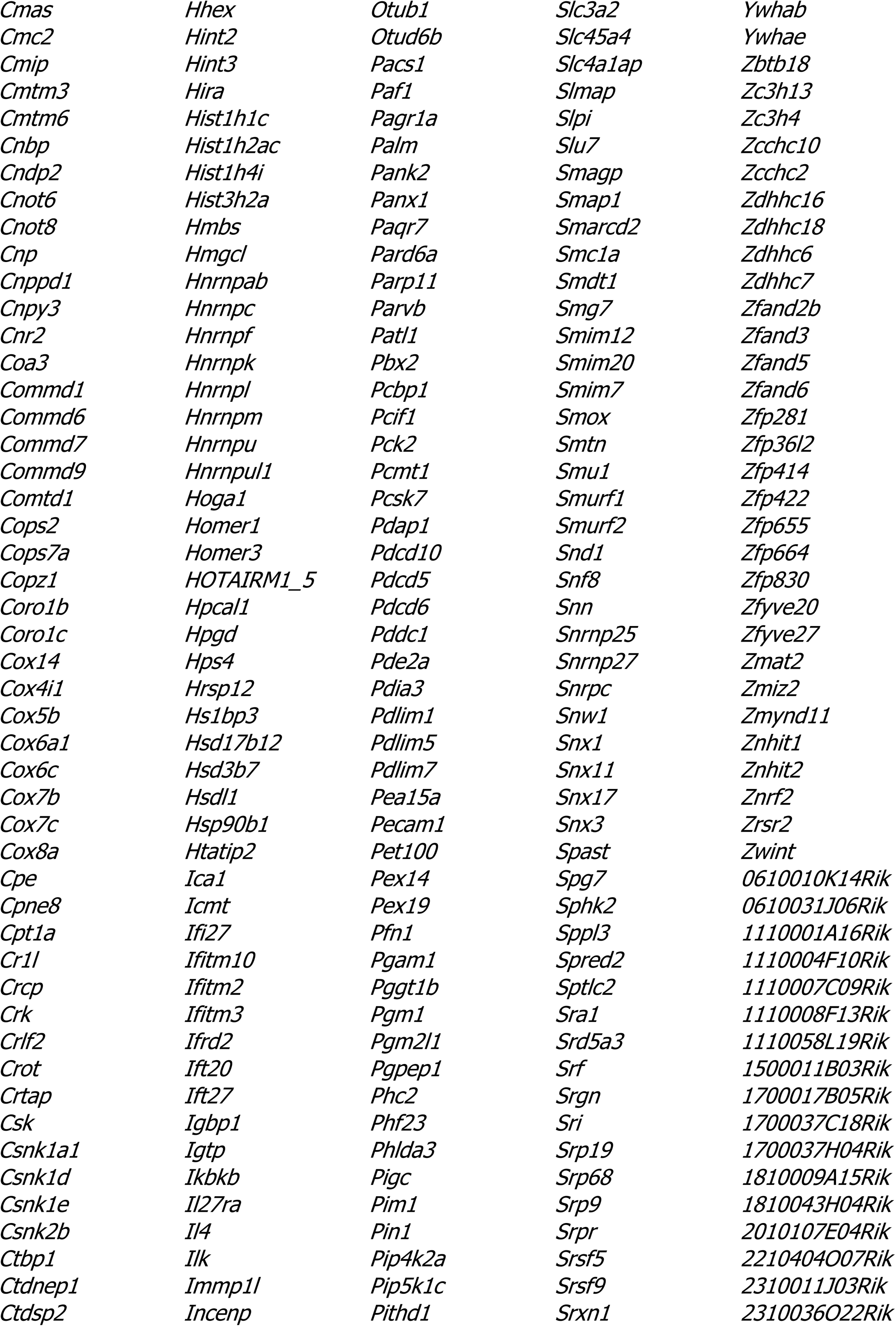
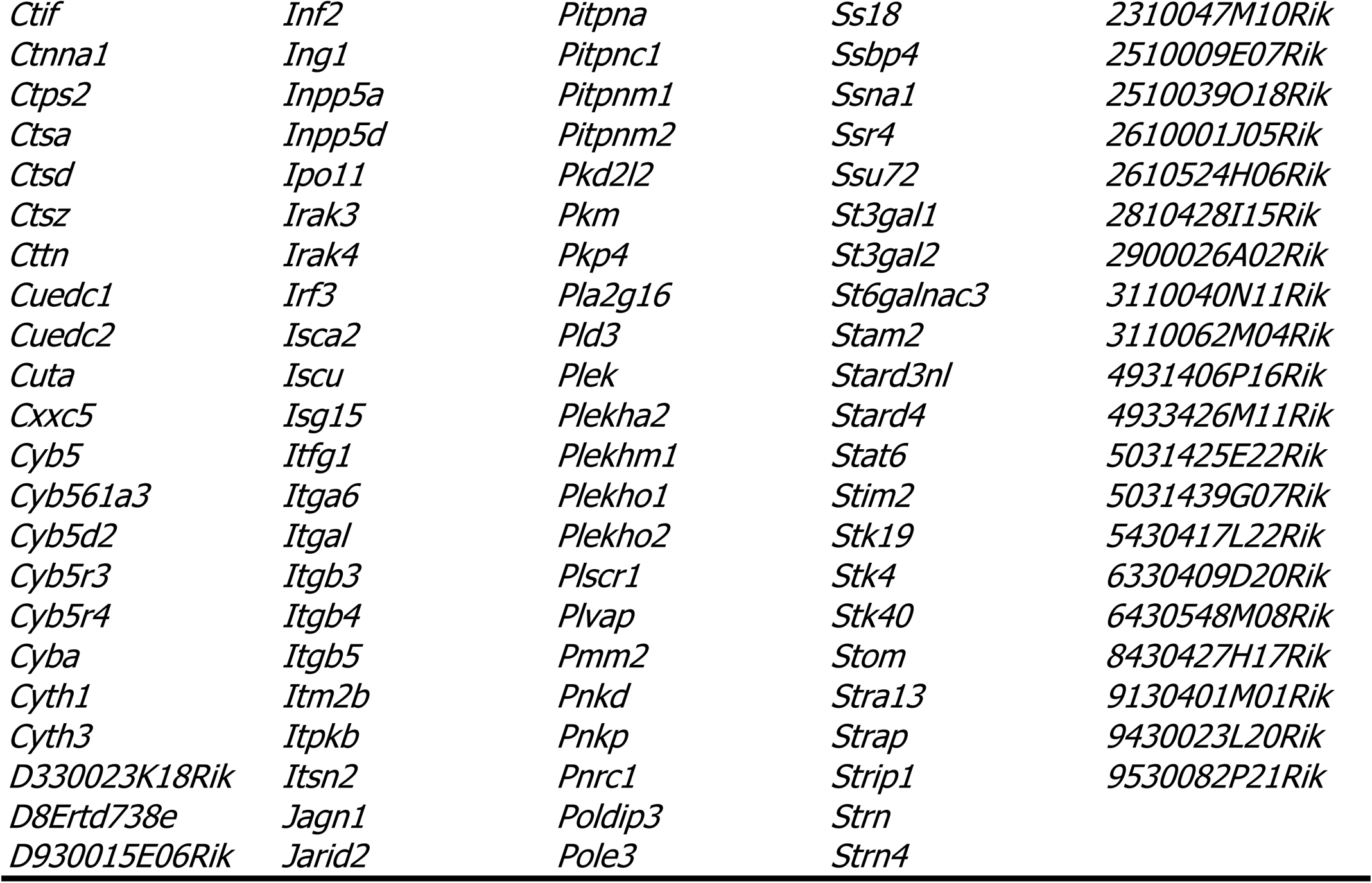
Tek-enriched genes also enriched in platelet TRAP of Rpl22<fl/fl>, Pf4-Cre<+/0> mice.

## References

1. Aird WC (2007) Phenotypic heterogeneity of the endothelium: II. Representative vascular beds. Circulation research 100(2):174–190.

2. Potente M & Makinen T (2017) Vascular heterogeneity and specialization in development and disease. Nature reviews. Molecular cell biology 18(8):477–494.

3. Ribatti D, Nico B, Vacca A, Roncali L, & Dammacco F (2002) Endothelial cell heterogeneity and organ specificity. Journal of hematotherapy & stem cell research 11(1):81–90.

4. Atkins GB, Jain MK, & Hamik A (2011) Endothelial differentiation: molecular mechanisms of specification and heterogeneity. Arteriosclerosis, thrombosis, and vascular biology 31(7):1476–1484.

5. McCarron JG, et al. (2019) Heterogeneity and emergent behaviour in the vascular endothelium. Current opinion in pharmacology 45:23–32.

6. Aird WC, et al. (1997) Vascular bed-specific expression of an endothelial cell gene is programmed by the tissue microenvironment. The Journal of cell biology 138(5):1117–1124.

7. Pusztaszeri MP, Seelentag W, & Bosman FT (2006) Immunohistochemical expression of endothelial markers CD31, CD34, von Willebrand factor, and Fli-1 in normal human tissues. The journal of histochemistry and cytochemistry : official journal of the Histochemistry Society 54(4):385–395.

8. Aman J, Weijers EM, van Nieuw Amerongen GP, Malik AB, & van Hinsbergh VW (2016) Using cultured endothelial cells to study endothelial barrier dysfunction: Challenges and opportunities. American journal of physiology. Lung cellular and molecular physiology 311(2):L453–466.

9. Helms HC, et al. (2016) In vitro models of the blood-brain barrier: An overview of commonly used brain endothelial cell culture models and guidelines for their use. Journal of cerebral blood flow and metabolism : official journal of the International Society of Cerebral Blood Flow and Metabolism 36(5):862–890.

10. Meyer J, Gonelle-Gispert C, Morel P, & Buhler L (2016) Methods for Isolation and Purification of Murine Liver Sinusoidal Endothelial Cells: A Systematic Review. PloS one 11(3):e0151945.

11. Amatschek S, et al. (2007) Blood and lymphatic endothelial cell-specific differentiation programs are stringently controlled by the tissue environment. Blood 109(11):4777–4785.

12. Durr E, et al. (2004) Direct proteomic mapping of the lung microvascular endothelial cell surface in vivo and in cell culture. Nature biotechnology 22(8):985–992.

13. Lacorre DA, et al. (2004) Plasticity of endothelial cells: rapid dedifferentiation of freshly isolated high endothelial venule endothelial cells outside the lymphoid tissue microenvironment. Blood 103(11):4164–4172.

14. Calabria AR & Shusta EV (2008) A genomic comparison of in vivo and in vitro brain microvascular endothelial cells. Journal of cerebral blood flow and metabolism : official journal of the International Society of Cerebral Blood Flow and Metabolism 28(1):135–148.

15. Pasqualini R & Ruoslahti E (1996) Organ targeting in vivo using phage display peptide libraries. Nature 380(6572):364–366.

16. Rajotte D, et al. (1998) Molecular heterogeneity of the vascular endothelium revealed by in vivo phage display. The Journal of clinical investigation 102(2):430–437.

17. Simonson AB & Schnitzer JE (2007) Vascular proteomic mapping in vivo. Journal of thrombosis and haemostasis : JTH 5 Suppl 1:183–187.

18. Tang FHF, et al. (2019) A ligand motif enables differential vascular targeting of endothelial junctions between brain and retina. Proceedings of the National Academy of Sciences of the United States of America 116(6):2300–2305.

19. Brunskill EW & Potter SS (2010) Gene expression programs of mouse endothelial cells in kidney development and disease. PloS one 5(8):e12034.

20. Daneman R, et al. (2010) The mouse blood-brain barrier transcriptome: a new resource for understanding the development and function of brain endothelial cells. The Journal of neuroscience : the official journal of the Society for Neuroscience 5(10):e13741.

21. Zhang Y, et al. (2014) An RNA-sequencing transcriptome and splicing database of glia, neurons, and vascular cells of the cerebral cortex. 34(36):11929–11947.

22. Schlereth K, Weichenhan D, & Bauer T (2018) The transcriptomic and epigenetic map of vascular quiescence in the continuous lung endothelium. Science (New York, N.Y.) 7.

23. Sabbagh MF, et al. (2018) Transcriptional and epigenomic landscapes of CNS and non-CNS vascular endothelial cells. eLife 7.

24. Nolan DJ, et al. (2013) Molecular signatures of tissue-specific microvascular endothelial cell heterogeneity in organ maintenance and regeneration. Developmental cell 26(2):204–219.

25. Han X, et al. (2018) Mapping the Mouse Cell Atlas by Microwell-Seq. Cell 172(5):1091–1107.e1017.

26. Lother A, et al. (2018) Cardiac Endothelial Cell Transcriptome. Arteriosclerosis, thrombosis, and vascular biology 38(3):566–574.

27. Vanlandewijck M, et al. (2018) A molecular atlas of cell types and zonation in the brain vasculature. Nature 554(7693):475–480.

28. He L & Vanlandewijck M (2018) Single-cell RNA sequencing of mouse brain and lung vascular and vessel-associated cell types. 5:180160.

29. Anonymous (2018) Single-cell transcriptomics of 20 mouse organs creates a Tabula Muris. Nature 562(7727):367–372.

30. Karaiskos N, et al. (2018) A Single-Cell Transcriptome Atlas of the Mouse Glomerulus. Journal of the American Society of Nephrology : JASN 29(8):2060–2068.

31. Heiman M, et al. (2008) A translational profiling approach for the molecular characterization of CNS cell types. Cell 135(4):738–748.

32. Hupe M, Li MX, Gertow Gillner K, Adams RH, & Stenman JM (2014) Evaluation of TRAP-sequencing technology with a versatile conditional mouse model. Science signaling 42(2):e14.

33. Liu J, et al. (2014) Cell-specific translational profiling in acute kidney injury. The Journal of clinical investigation 124(3):1242–1254.

34. Santhosh D & Huang Z (2016) A Tie2-driven BAC-TRAP transgenic line for in vivo endothelial gene profiling. Genesis (New York, N.Y. : 2000) 54(3):136–145.

35. Sanz E, et al. (2009) Cell-type-specific isolation of ribosome-associated mRNA from complex tissues. Proceedings of the National Academy of Sciences of the United States of America 106(33):13939–13944.

36. Zhou P, et al. (2013) Interrogating translational efficiency and lineage-specific transcriptomes using ribosome affinity purification. Proceedings of the National Academy of Sciences of the United States of America 110(38):15395–15400.

37. King HA & Gerber AP (2016) Translatome profiling: methods for genome-scale analysis of mRNA translation. Briefings in functional genomics 15(1):22–31.

38. Everett LA, Cleuren AC, Khoriaty RN, & Ginsburg D (2014) Murine coagulation factor VIII is synthesized in endothelial cells. Blood 123(24):3697–3705.

39. Fahs SA, Hille MT, Shi Q, Weiler H, & Montgomery RR (2014) A conditional knockout mouse model reveals endothelial cells as the principal and possibly exclusive source of plasma factor VIII. Blood 123(24):3706–3713.

40. de Lange WJ, Halabi CM, Beyer AM, & Sigmund CD (2008) Germ line activation of the Tie2 and SMMHC promoters causes noncell-specific deletion of floxed alleles. Physiological genomics 35(1):1–4.

41. Koni PA, et al. (2001) Conditional vascular cell adhesion molecule 1 deletion in mice: impaired lymphocyte migration to bone marrow. The Journal of experimental medicine 193(6):741–754.

42. Muzumdar MD, Tasic B, Miyamichi K, Li L, & Luo L (2007) A global double-fluorescent Cre reporter mouse. Genesis (New York, N.Y. : 2000) 45(9):593–605.

43. Hsieh PC, Davis ME, Lisowski LK, & Lee RT (2006) Endothelial-cardiomyocyte interactions in cardiac development and repair. Annual review of physiology 68:51–66.

44. Ingolia NT, Lareau LF, & Weissman JS (2011) Ribosome profiling of mouse embryonic stem cells reveals the complexity and dynamics of mammalian proteomes. Cell 147(4):789–802.

45. Tang Y, Harrington A, Yang X, Friesel RE, & Liaw L (2010) The contribution of the Tie2+ lineage to primitive and definitive hematopoietic cells. Genesis (New York, N.Y. : 2000) 48(9):563–567.

46. Lechauve C, et al. (2018) Endothelial cell alpha-globin and its molecular chaperone alpha-hemoglobin-stabilizing protein regulate arteriolar contractility. The Journal of clinical investigation 128(11):5073–5082.

47. Straub AC, et al. (2012) Endothelial cell expression of haemoglobin alpha regulates nitric oxide signalling. Nature 491(7424):473–477.

48. Rowley JW, et al. (2011) Genome-wide RNA-seq analysis of human and mouse platelet transcriptomes. Blood 118(14):e101–111.

49. Aitsebaomo J, Portbury AL, Schisler JC, & Patterson C (2008) Brothers and sisters: molecular insights into arterial-venous heterogeneity. Circulation research 103(9):929–939.

50. Podgrabinska S, et al. (2002) Molecular characterization of lymphatic endothelial cells. Proceedings of the National Academy of Sciences of the United States of America 99(25):16069–16074.

51. Bhasin M, et al. (2010) Bioinformatic identification and characterization of human endothelial cell-restricted genes. BMC genomics 11:342.

52. Wallgard E, et al. (2008) Identification of a core set of 58 gene transcripts with broad and specific expression in the microvasculature. Arteriosclerosis, thrombosis, and vascular biology 28(8):1469–1476.

53. Butler LM, et al. (2016) Analysis of Body-wide Unfractionated Tissue Data to Identify a Core Human Endothelial Transcriptome. Cell systems 3(3):287–301.e283.

54. Rosenberg AB & Roco CM (2018) Single-cell profiling of the developing mouse brain and spinal cord with split-pool barcoding. 360(6385):176–182.

55. Adam M, Potter AS, & Potter SS (2017) Psychrophilic proteases dramatically reduce single-cell RNA-seq artifacts: a molecular atlas of kidney development. 144(19):3625–3632.

56. Kang SS, Baker KE, Wang X, Kocher J-P, & Fryer JD (2017) Translational profiling of microglia reveals artifacts of cell sorting. bioRxiv.

57. van den Brink SC, Sage F, & Vertesy A (2017) Single-cell sequencing reveals dissociation-induced gene expression in tissue subpopulations. 14(10):935–936.

58. Hicks SC, Townes FW, Teng M, & Irizarry RA (2018) Missing data and technical variability in single-cell RNA-sequencing experiments. Biostatistics (Oxford, England) 19(4):562–578.

59. Cuevas-Diaz Duran R, Wei H, & Wu JQ (2017) Single-cell RNA-sequencing of the brain. Clinical and translational medicine 6(1):20.

60. Banks WA & Robinson SM (2010) Minimal penetration of lipopolysaccharide across the murine blood-brain barrier. Brain, behavior, and immunity 24(1):102–109.

61. Aird WC (2003) The role of the endothelium in severe sepsis and multiple organ dysfunction syndrome. Blood 101(10):3765–3777.

62. Ince C, et al. (2016) THE ENDOTHELIUM IN SEPSIS. Shock (Augusta, Ga.) 45(3):259–270.

63. Komarova YA, Kruse K, Mehta D, & Malik AB (2017) Protein Interactions at Endothelial Junctions and Signaling Mechanisms Regulating Endothelial Permeability. Circulation research 120(1):179–206.

64. von Drygalski A, Furlan-Freguia C, Ruf W, Griffin JH, & Mosnier LO (2013) Organ-specific protection against lipopolysaccharide-induced vascular leak is dependent on the endothelial protein C receptor. Arteriosclerosis, thrombosis, and vascular biology 33(4):769–776.

65. Pu W, et al. (2018) Genetic Targeting of Organ-Specific Blood Vessels. Circulation research 123(1):86–99.

## Supplemental references

1. Sanz E, et al. (2009) Cell-type-specific isolation of ribosome-associated mRNA from complex tissues. Proceedings of the National Academy of Sciences of the United States of America 106(33):13939–13944.

2. de Lange WJ, Halabi CM, Beyer AM, & Sigmund CD (2008) Germ line activation of the Tie2 and SMMHC promoters causes noncell-specific deletion of floxed alleles. Physiological genomics 35(1):1–4.

3. Koni PA, et al. (2001) Conditional vascular cell adhesion molecule 1 deletion in mice: impaired lymphocyte migration to bone marrow. The Journal of experimental medicine 193(6):741–754.

4. Muzumdar MD, Tasic B, Miyamichi K, Li L, & Luo L (2007) A global double-fluorescent Cre reporter mouse. Genesis (New York, N.Y. : 2000) 45(9):593–605.

5. Tiedt R, Schomber T, Hao-Shen H, & Skoda RC (2007) Pf4-Cre transgenic mice allow the generation of lineage-restricted gene knockouts for studying megakaryocyte and platelet function in vivo. Blood 109(4):1503–1506.

6. Lakso M, et al. (1996) Efficient in vivo manipulation of mouse genomic sequences at the zygote stage. Proceedings of the National Academy of Sciences of the United States of America 93(12):5860–5865.

7. Andreev DE, O’Connor PB, Loughran G, & Dmitriev SE (2017) Insights into the mechanisms of eukaryotic translation gained with ribosome profiling. 45(2):513–526.

8. Nolan DJ, et al. (2013) Molecular signatures of tissue-specific microvascular endothelial cell heterogeneity in organ maintenance and regeneration. Developmental cell 26(2):204–219.

9. Bodary PF, Westrick RJ, Wickenheiser KJ, Shen Y, & Eitzman DT (2002) Effect of leptin on arterial thrombosis following vascular injury in mice. Jama 287(13):1706–1709.

10. Mudge JM & Harrow J (2015) Creating reference gene annotation for the mouse C57BL6/J genome assembly. Mammalian genome : official journal of the International Mammalian Genome Society 26(9-10):366–378.

11. Langmead B & Salzberg SL (2012) Fast gapped-read alignment with Bowtie 2. Nature methods 9(4):357–359.

12. Salzman J, Jiang H, & Wong WH (2011) Statistical Modeling of RNA-Seq Data. Statistical science : a review journal of the Institute of Mathematical Statistics 26(1).

13. Pfeffer S, Woellhaf MW, Herrmann JM, & Forster F (2015) Organization of the mitochondrial translation machinery studied in situ by cryoelectron tomography. Nature communications 6:6019.

14. Anonymous (2015) Gene Ontology Consortium: going forward. Nucleic acids research 43(Database issue):D1049–1056.

15. Ashburner M, et al. (2000) Gene ontology: tool for the unification of biology. The Gene Ontology Consortium. Nature genetics 25(1):25–29.

16. Szklarczyk D, et al. (2017) The STRING database in 2017: quality-controlled protein-protein association networks, made broadly accessible. 45(D1):D362–d368.

17. Rowley JW, et al. (2011) Genome-wide RNA-seq analysis of human and mouse platelet transcriptomes. Blood 118(14):e101–111.

18. Jin E, et al. (2009) Differential roles for ETS, CREB, and EGR binding sites in mediating VEGF receptor 1 expression in vivo. Blood 114(27):5557–5566.

19. Du P, Kibbe WA, & Lin SM (2008) lumi: a pipeline for processing Illumina microarray. Bioinformatics (Oxford, England) 24(13):1547–1548.

20. Ritchie ME, et al. (2015) limma powers differential expression analyses for RNA-sequencing and microarray studies. Nucleic acids research 43(7):e47.

21. Wang F, et al. (2012) RNAscope: a novel in situ RNA analysis platform for formalin-fixed, paraffin-embedded tissues. The Journal of molecular diagnostics : JMD 14(1):22–29.

